# Endochondral bone in an Early Devonian ‘placoderm’ from Mongolia

**DOI:** 10.1101/2020.06.09.132027

**Authors:** Martin D. Brazeau, Sam Giles, Richard P. Dearden, Anna Jerve, Y.A. Ariunchimeg, E. Zorig, Robert Sansom, Thomas Guillerme, Marco Castiello

## Abstract

Endochondral bone is the main internal skeletal tissue of nearly all osteichthyans—the group comprising more than 60,000 living species of bony fishes and tetrapods. Chondrichthyans (sharks and their kin) are the living sister group of osteichthyans and have cartilaginous endoskeletons, long considered the ancestral condition for all jawed vertebrates (gnathostomes). The absence of bone in modern jawless fishes and the absence of endochondral ossification in early fossil gnathostomes appears to lend support to this conclusion. Here we report the discovery of extensive endochondral bone in *Minjinia turgenensis*, a new genus and species of ‘placoderm’-like fish from the Early Devonian (Pragian) of western Mongolia described using x-ray computed microtomography (XR-µCT). The fossil consists of a partial skull roof and braincase with anatomical details providing strong evidence of placement in the gnathostome stem group. However, its endochondral space is filled with an extensive network of fine trabeculae resembling the endochondral bone of osteichthyans. Phylogenetic analyses place this new taxon as a proximate sister group of the gnathostome crown. These results provide direct support for theories of generalised bone loss in chondrichthyans. Furthermore, they revive theories of a phylogenetically deeper origin of endochondral bone and its absence in chondrichthyans as a secondary condition.

---

The vertebrate skeleton comprises two main systems: the exoskeleton (external achondral dermal bones) and endoskeleton (internal chondral bones and cartilages, as well as some intramembranous bones)^1^. An ossified exoskeleton evolved at least 450 million years ago in jawless stem gnathostomes^2,3^, but the endoskeleton in these taxa is not endochondrally ossified. Endochondral bone, in which the cartilaginous endoskeletal precursor is invaded by and eventually replaced by bone, is widely considered an osteichthyan apomorphy^3-7^ and such a reliable identifying character gives the group its name. Extant chondrichthyans lack dermal bone and possess a mainly cartilaginous endoskeleton enveloped by a structurally diverse range of tessellate calcified cartilage^8^. Outgroups of the gnathostome crown also lack endochondral ossification. Galeaspids surround their cartilaginous skeleton in globular calcified cartilage{NianZhong:2005tj}, while osteostracan and ‘placoderm’ endoskeletons were sheathed in perichondral bone^3^. Consequently, the last common ancestor of jawed vertebrates was long thought to have been perichondrally ossified, but lacking endochondral ossification^3^.

In this paper, we describe a new genus and species of ‘placoderm’ from the Early Devonian of western Mongolia. Although Mongolia is known for some of the geologically oldest putative gnathostome fossils (isolated chondrichthyan-like scales ^9-12^), it remains a poorly sampled region of the world with respect to early vertebrates. ‘Placoderms’ were until now known from only a single fragmentary occurrence^13^ in the early Middle Devonian (Eifelian). Our new data highlight the importance of Mongolia as a key region for studies of early gnathostome evolution. We describe a braincase and partial skull roof representing the first substantial body fossil of an early gnathostome from Mongolia and displaying an unexpected occurrence of endochondral bone analysed using XR-µCT. We conducted phylogenetic analyses to reconstruct the evolutionary relationships of this new taxon. To explore the evolutionary history of endochondral bone in light of this new discovery, we used parsimony and maximum likelihood ancestral states reconstruction. Finally, we discuss these results in the context of earlier statements about endochondral bone in non-osteichthyans, new developments in understanding the complexity and diversity of chondrichthyan endoskeletal tissues, and current uncertainties about early gnathostome phylogenetic relationships.

## Systematic palaeontology

> Gnathostomata Gegenbaur, 1874^14^
>
> *Minjinia turgenensis* gen. et sp. nov.

### Etymology

Generic name honours the memory of Chuluun Minjin for his extensive contributions to the Palaeozoic stratigraphy of Mongolia, his enthusiastic support of this work, and introducing us to the Yamaat River locality. Specific name recognises the provenance of the fossil from the Turgen region, Uvs aimag of western Mongolia.

### Holotype

Institute of Paleontology, Mongolian Academy of Sciences MPC-FH100/9.1, a partial braincase and skull roof.

### Type locality

Turgen Strictly Protected Area, Uvs province, western Mongolia; near the top of the stratigraphic sequence that occurs between the Tsagaan-Salaat and Yamaat Rivers.

### Formation and age

Upper part of Tsagaansalaat Formation, Pragian (Early Devonian) ^15,16^.

### Diagnosis

‘Placoderm’-grade stem gnathostome with endochondral bone, deep epaxial muscle cavities flanking a slender occipital ridge, and the following possible autapomorphies: dermal bones covered in sparsely placed tubercles, penultimate spino-occipital nerve canal substantially larger in diameter than others.

## Description

MPC-FH100/9.1 consists of a partial braincase and skull roof (Fig. 1). The skull roof is ornamented with sparsely distributed finely ridged tubercles resembling those of the Siberian ‘placoderm’ *Dolganosteus*^17^; the tubercles become more broadly separated towards the midline of the skull. They are distinct from those of *Dolganosteus* in that towards the midline of the skull roof, the tubercles are larger and more pointed. The specimen shows signs of extensive post-mortem transport, with angles of the braincase worn off and much of the skull roof and some of the braincase preserved as a mould. Individual skull roof ossifications cannot be identified, although this may be due to the dominantly mouldic preservation. There appears to have been a prominent nuchal plate eminence comparable to certain acanthothoracids such as *Romundina*^18^ and *Arabosteus*^19^.

**Fig. 1.**
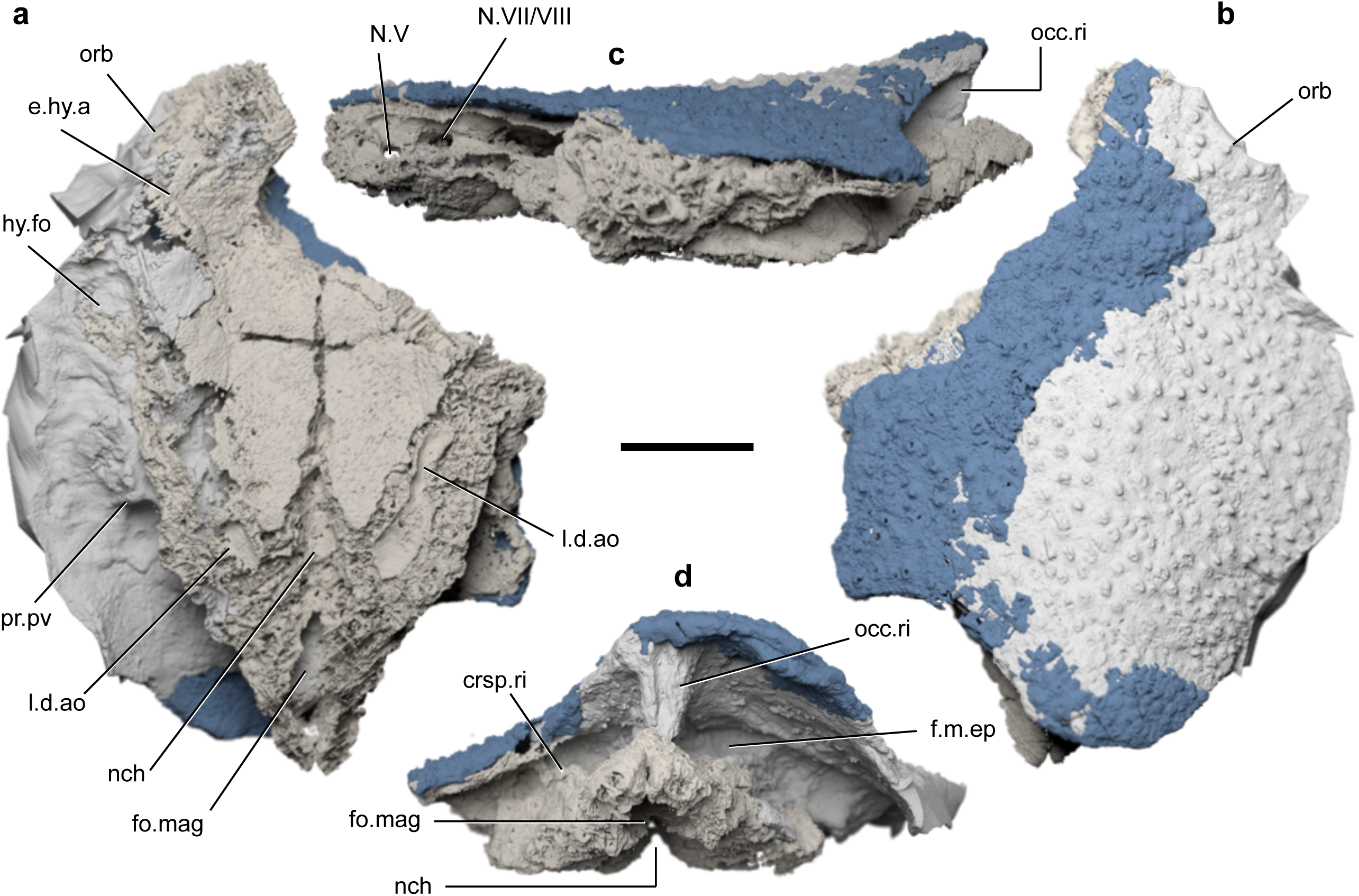
MPC-FH100/9.1 a ‘placoderm’ skull roof and braincase from the Early Devonian of Mongolia. **a**, Ventral view. **b**, Dorsal view. **c**, Left lateral view. **d**, Posterior view. **e**, Braincase endocavity in dorsal view. Taupe: endoskeleton; grey: mould; blue: exoskeleton. crsp.ri, craniospinal ridge; e.hy.a, sulcus for the efferent hyoid artery; f.m.ep, epaxial muscle fossa; fo.mag., foramen magnum; hy.fo, hyodean fossa; l.d.ao, sulcus for the lateral dorsal aorta; N.V, trigeminal nerve canal; N.VII, facial nerve canal; N.VIII, acoustic nerve canal; nch, notochordal canal; occ.ri, occipital ridge; orb, orbit; pr.pv, paravagal process. Scale bar, 10 mm.

### Endoskeletal tissue

The braincase of MPC-FH100/9.1 is well ossified, comprising an external bony sheath filled with an extensive matrix of spongy tissue (Fig. 2a-b; Extended Data Fig. 1; Supplementary Video 1). The trabecles forming this tissue are irregular and branching, less than 1 mm thick and often curved, and resemble most closely the endochondral tissue of osteichthyans (Fig. 2c-d; Supplementary Video 2). As such, we interpret this as endochondral bone. Notably, this is found in all preserved regions of the braincase, in contrast to the isolated trabeculae previously identified as endochondral bone in *Boreaspis*^20^ and *Bothriolepis*^21^. The margins of the braincase, the endocranial walls, and the boundaries of nerve and blood canals, are formed from a thicker tissue which we interpret as perichondral bone. This suggests that the endoskeleton of *Minjinia* comprises osteichthyan-like endochondral bone, with an ossified perichondrium. To address the possible alternative explanation that it is an aberrant instance of calcified cartilage, we compared the structure of this tissue with rarely-preserved mineralized cartilage in the stem chondrichthyan *Diplacanthus crassismus* (National Museums of Scotland specimen NMS 1891.92.334; Fig. 2e-f) observed using synchrotron tomography. The cancellae within the endochondral tissue of *Minjinia* are irregular, with a diameter of approximately 1-2 mm. This tissue is distinctly unlike the calcified cartilage of *Diplacanthus* in appearance, which consists of a densely packed matrix of irregularly stacked chondrons between 20-60 µm in diameter.

**Fig. 2.**
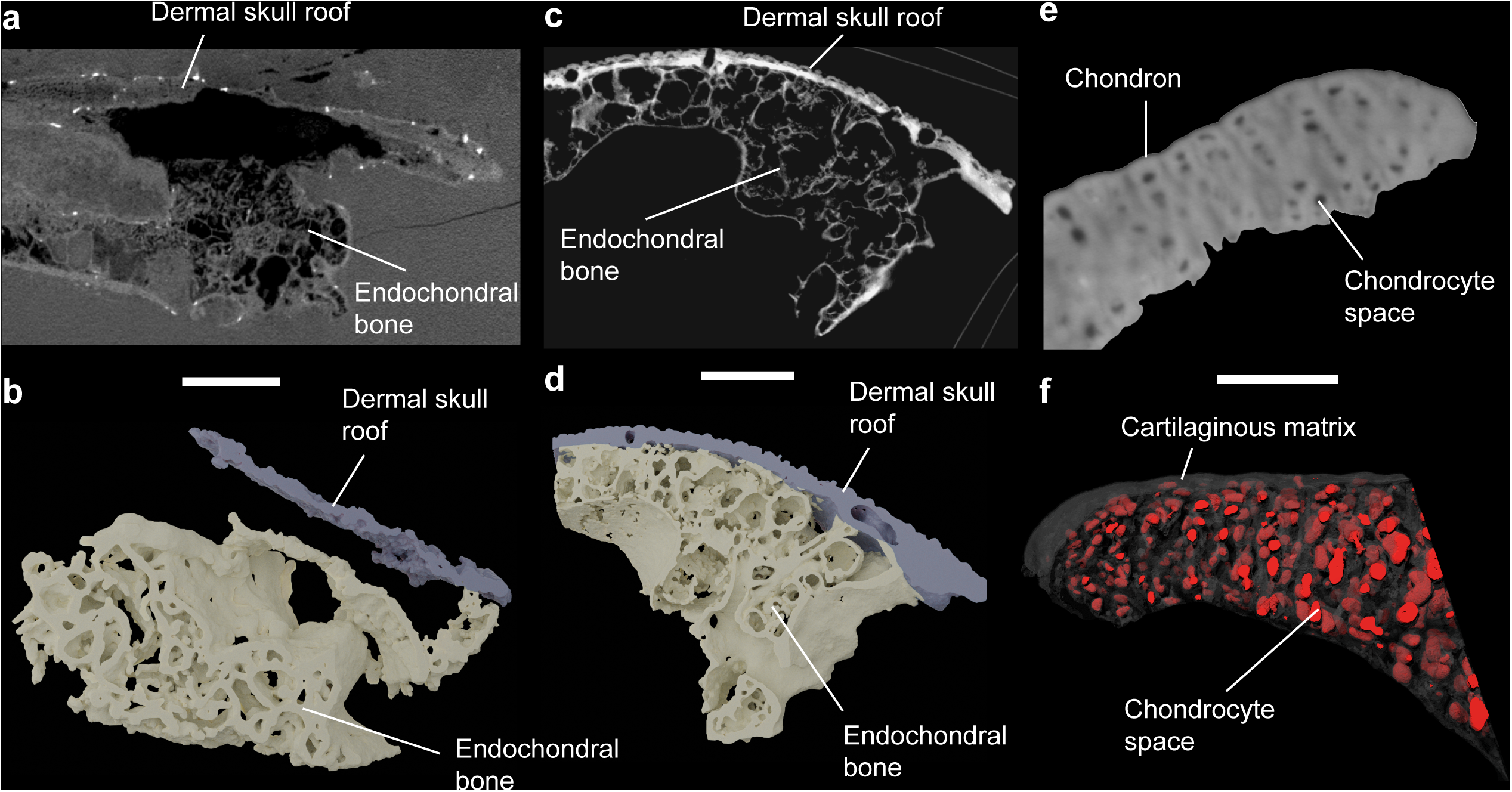
Endoskeletal mineralisation in fossil gnathostomes. **a**, Transverse tomographic slice through MPC-FH100/9.1. **b**, Three-dimensional rendering of trabecular bone structure. **c**, Transverse tomographic section through the braincase of the osteichthyan *Ligulalepis*. **d**, Three-dimensional rendering of the trabecular bone in *Ligulalepis* (**c** and **d** use data from^56^). **e**, Synchrotron tomography image of the calcified cartilage of the certatohyal of the stem-group chondrichthyan *Diplacanthus crassisimus* specimen NMS 1891.92.334. **f**, Semi-transparent three-dimensional structure of calcified cartilage of NMS 1891.92.334. Scale bars, **a** and **b**, 10 mm; **c** and **d**, 1 mm; **e** and **f**, 150 µm.

### Neurocranium

The braincase is preserved from the level of the right posterior orbital wall to the posterior end of the occipital ridge. Occipital glenoid condyles are not preserved, but much of the rest of the broad, flat parachordal region is present, separated by a midline groove that accommodated a relatively narrow notochordal tunnel. An asymmetric transverse fissure spans the basicranial surface at about mid-length of the preserved portion. It appears to demarcate the anterior margin of the parachordal plates and may correspond to the ventral cranial fissure of crown-group gnathostomes. However, unlike in crown gnathostomes, it is traversed by a substantial anterior extension of the cranial notochord. The courses of the lateral dorsal aortae are marked by a pair of sulci on the lateral margins of the parachordal plates, though only a short part of the canal is preserved on the right side of the specimen. A narrow, shallow sulcus for the efferent hyoid artery is present on the preserved right side of the specimen, immediately behind the level of the orbit (Fig. 1a).

The lateral surface of the braincase is preserved on the right side as a mouldic impression in the matrix (Fig. 1). A sharply demarcated hyoid fossa is present on the lateral wall of the otic region (Fig. 1). Posterior to this, a stout but pronounced vagal process with a pair of rounded eminences likely corresponds to the branchial arch articulations. There is no evidence for a pair of anterior and posterior divisions to the vagal process, which are typically seen in other ‘placoderms’. A well-developed ‘placoderm’-like craniospinal process is absent; its homologous position is instead covered in perichondral bone and marked by a low ridge (Fig. 1).

In posterior view, a tall, narrow median occipital ridge is evident and resembles the morphology of *Romundina*^22^ and *Arabosteus*^19^. Similar to these taxa, the median otic ridge is flanked by two large occipital fossae for the epaxial musculature. The notochordal tunnel is approximately the same size as or smaller than the foramen magnum, as in ‘placoderms’ and in contrast with crown-group gnathostomes. A metotic fissure is absent.

### Endocast

A partial cranial endocast is preserved, consisting of the hindbrain cavity, partial midbrain cavity, labyrinth cavities, and posteromedial corner of the orbital region. The two primary trunk canals of the trigeminal nerve (N.V_1_ and N.V_2,3_) are preserved (Fig. 3). The acoustic (N.VIII) and facial nerve (N.VII) canals share a common trunk canal behind the trigeminal nerves, as in many other ‘placoderms’ ^22-25^. The facial nerve canal branches into palatal and hyomandibular branches between the saccular chamber and rear orbit wall (Fig. 3; Extended Data Fig. 2), indicating this division was internal (deep) to the otic process. The supraophthalmic branch opens into the rear wall of the orbit and part of its supraorbital course is preserved (Fig. 3; Extended Data Fig. 2). A slender branch extends below the labyrinth and divides into palatine and hyomandibular branches (Fig 3; Extended Data Fig. 2). As in other ‘placoderm’-grade taxa, the vagus nerve (N. X) trunk canal is very large in diameter and exits from immediately behind the labyrinth cavity (Fig. 3). The spino-occipital region resembles other ‘placoderms’ in being extended. At least four spino-occipital nerve canals are present in a linear series, and the penultimate canal is largest in diameter (Fig. 3). Intercalating these is a network of occipital artery canals branching from the dorsal aortae.

**Fig. 3.**
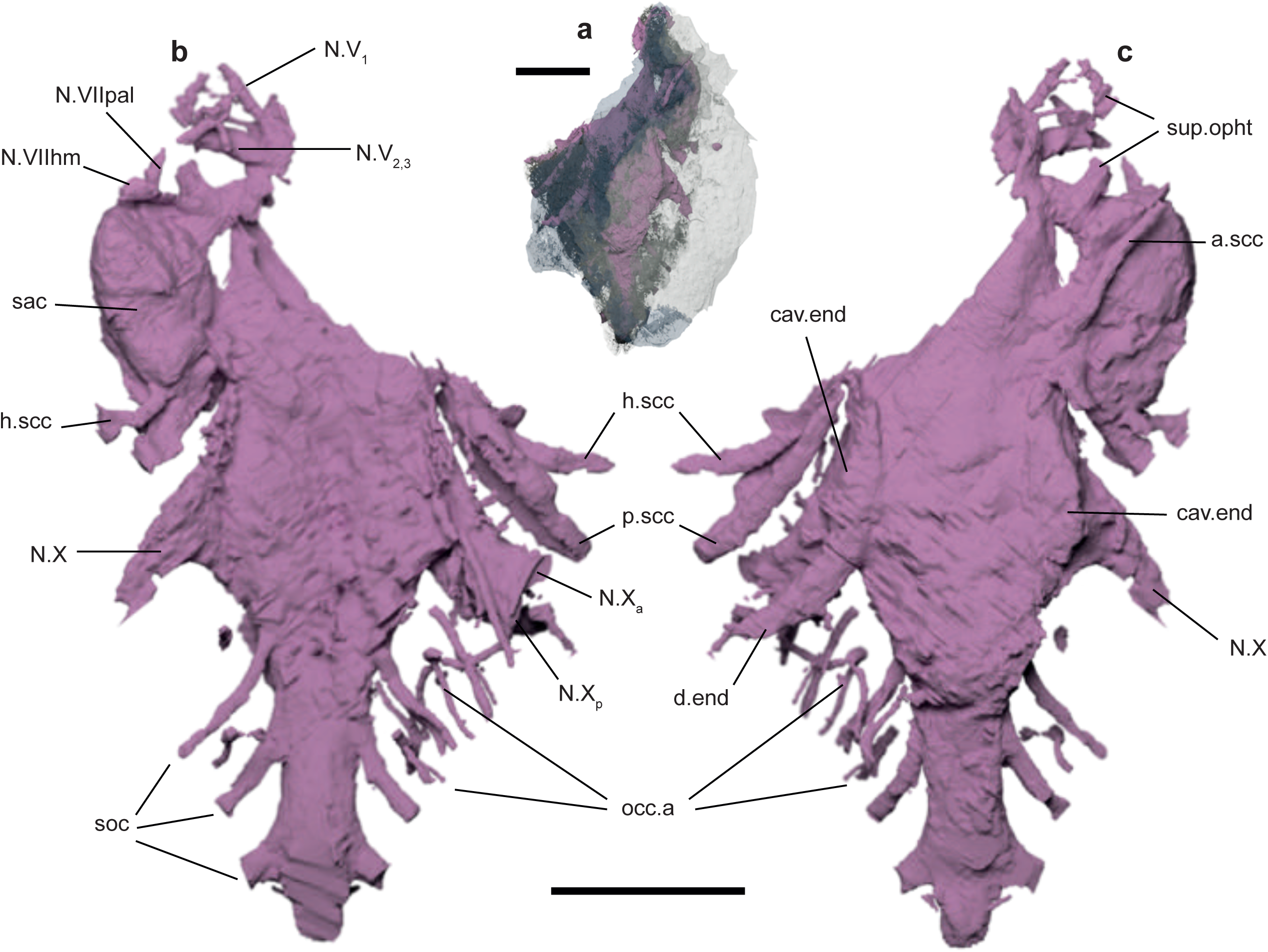
Braincase endocavity of *Minjinia*. **a**, Semi-transparent rendering of skull roof and braincase (grey and blue) showing extent of endocavity (pink). **b**, Ventral view. **c**, Dorsal view. a.scc, anterior semicircular canal; cav.end, endolymphatic cavity; d.end, endolymphatic duct; h.scc, horizontal semicicular canal; N.V, trigeminal nerve canal; N.VIIhm, hyomandibular branch of facial nerve canal; N.VIIpal, palatine branch of facial nerve canal; N.VIII, acoustic nerve canal; N.X, vagus nerve canal, N.X_a_, anterior branch of vagus nerve canal; N.X_p_, posterior branch of vagus nerve canal; occ.a, occipital artery canals; p.scc, posterior semicircular canal; sac, sacculus; soc, spino-occipital nerve canals; sup.opth, canal for supra-ophtalmic nerve. Scale bars, 10 mm (upper scale bar associates with **a**, lower scale bar associates with **b** and **c**).

The skeletal labyrinth is not complete on either side of the specimen, but can mostly be reconstructed according to the assumption of bilateral symmetry. The most significant feature is that the labyrinth and endolymphatic cavity are joined to the main endocavity chamber (Fig. 3). This is a striking contrast to other ‘placoderms’ and closely resembles crown-group gnathostomes^26^. The endolymphatic canals are elongate and tubular, extending posterolaterally to reach the skull roof, though external openings cannot be clearly identified. The anterior semi-circular canal follows the saccular cavity closely as in petalichthyids^27^(Fig. 3). However, the horizontal and posterior canals appear to extend well away from the saccular chamber (Fig. 3). The dorsal junctions of the anterior and posterior canals are joined in a crus commune, as in *Romundina*^22^ and *Jagorina*^23^. A sinus superior is absent.

## Phylogenetic analyses

We conducted phylogenetic analyses under four different protocols: equal weights parsimony, implied weights parsimony, an unpartitioned Bayesian analysis, and a Bayesian analysis with characters partitioned by fit determined under implied weights parsimony^28^ (see Extended Data Figs. 3-6). All phylogenetic analyses consistently place *Minjinia* as a stem-group gnathostome, proximate to the gnathostome crown (Fig. 4, Extended Data Figs 3, 4). *Minjinia* is recovered in a position crownward of arthrodires but outside of a grade consisting of *Entelognathus, Ramirosuarezia*, and *Janusiscus*. Under implied weights parsimony, these three taxa move onto the osteichthyan stem and *Minjinia* is placed as the immediate sister taxon of the gnathostome crown. Under parsimony, the crownward position of *Minjinia* is unambiguously supported by the skeletal labyrinth and endolymphatic duct being confluent with the main cranial cavity^26^ (Supplementary Information). In common with arthrodires and the gnathostome crown^29^, *Minjinia* possesses a division of the facial nerve(Fig. 3; Extended Data Fig. 2) deep to the transverse otic process. However, *Minjinia* is excluded from the gnathostome crown group due to the absences of a metotic fissure and a posterior dorsal fontanelle, and presence of broad, flat parachordal plates expanded behind the saccular cavity (Fig. 3, Supplementary Information).

**Fig. 4.**
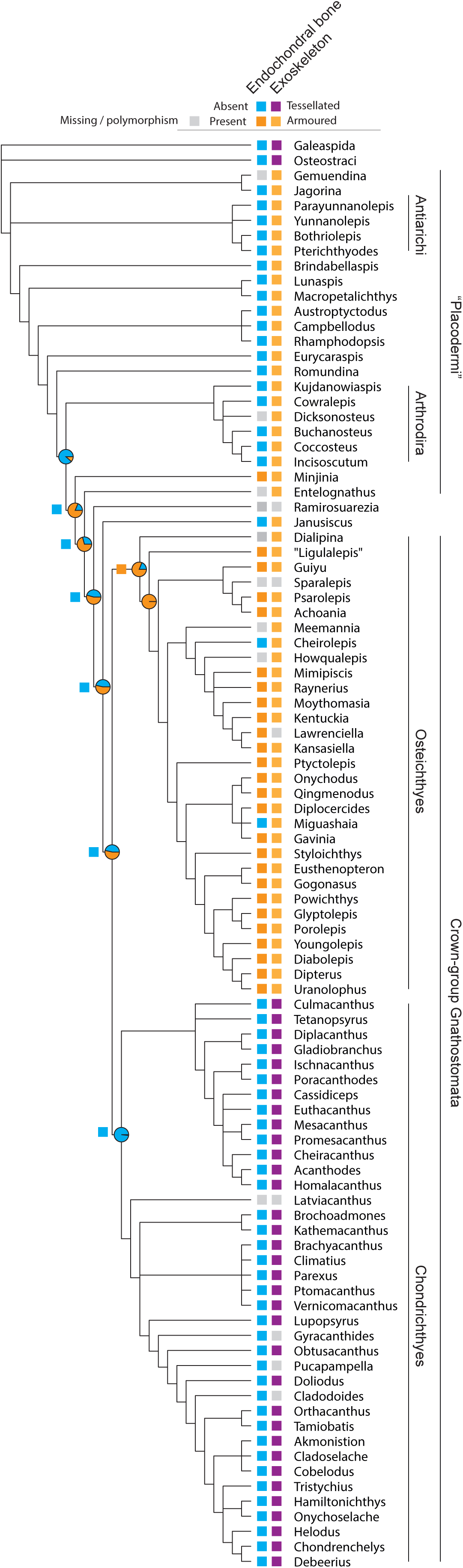
Strinct consensus tree from parsimony analysis of early gnathostomes showing distribution of endochondral bone and exoskeletal armour. Squares at nodes indicate parsimony reconstruction for endochondral bone. Pie charts at nodes show likelihood reconstructions for the same character under the all-rates-different model (see Extended Data Figs 6 & 7 for competing reconstructions). Grey box indicates uncertainty. Loss of endochondral bone maps closely with generalised loss of bone in chondrichthyans where exoskeletal armour and perichondral bone are also absent.

We undertook ancestral states reconstructions to assess the evolutionary history of endochondral bone (Fig. 4; Extended Data Figs. 5 & 6; Supplementary Information). Interestingly, parsimony analysis fails to recover secondary homology of this trait between *Minjinia* and osteichthyans. The crownward placement of *Minjinia* is, in fact, based on independent evidence relating to anatomical features of the braincase and endocast. However, the resolution becomes ambiguous if missing data in either *Entelognathus* or *Ramirosuarezia* are resolved as having endochondral bone. The reconstruction becomes similarly ambiguous if *Janusiscus* is moved a single branch (requiring only two additional steps) onto the chondrichthyan stem. The strict precision of parsimony reconstructions makes it insensitive to this underlying uncertainty. To explore this, we used likelihood reconstructions and compared the ancestral state reconstructions under equal rates (ER) and all rates different (ARD) variants of the Mkv model on branch-length-rescaled parsimony trees and Bayesian trees. Both models show substantial non-zero marginal likelihoods if endochondral bone is assumed present in the common node of *Minjinia* and Osteichthyes, with ARD strongly favouring its presence (0.33 for ER; 0.81 for ARD; Fig. 3, Table 1, Extended Data Figs. 5, 6, Supplementary Table 1). Under the ARD model, there is nearly equivocal support for presence or absence of endochondral bone at the gnathostome crown node (Table 1). The ARD model shows the best fit for endochondral bone (likelihood ratios 4.75 for parsimony [p = 0.029] and 5.26 for Bayesian, [p = 0.022]) (Table 1, Supplementary Table 1), favouring repeated losses of this tissue over multiple gains (see Discussion).

**Table 1.** Tree distribution (n=100) ancestral states estimation results. ER = Equal rates model; ARD = All Rates Different model. The columns AIC and log.like represent the median AIC and log.lik across the 100 parsimony and Bayesian trees (for both models). The like.ratio column is the likelihood ratio for the models compared on these trees. The columns Absent and Present represent the median scaled likelihood for the endochondral bone state.

| trees | model | log.like. | like.ratio | AIC | node | Absent | Present |
| --- | --- | --- | --- | --- | --- | --- | --- |
| Parsimony | ER | -28.91 | 4.74 | 59.82 | Minjinia:Gnathostomes | 0.67 | 0.33 |
| (equal weights) | ARD | -26.54 |  | 57.09 |  | 0.19 | 0.81 |
|  | ER |  |  |  | Crown Gnathostomes | 0.91 | 0.09 |
|  | ARD |  |  |  |  | 0.46 | 0.54 |
| Bayesian | ER | -29.66 | 5.26 | 61.32 | Minjinia:Gnathostomes | 0.73 | 0.27 |
| (unpartitioned) | ARD | -27.03 |  | 58.06 |  | 0.17 | 0.83 |
|  | ER |  |  |  | Crown Gnathostomes | 0.79 | 0.21 |
|  | ARD |  |  |  |  | 0.22 | 0.78 |

## Discussion

*Minjinia turgenensis* presents an unexpected discovery of extensive endochondral bone in a ‘placoderm’-grade fish, with repercussions for the phylogenetic origin of this tissue and the problem of early gnathostome relationships more generally. The prevailing hypothesis has been that endochondral bone is an osteichthyan apomorphy^3,7,29^. However, recent discoveries have cast doubt on this assertion. The recognition that dermal bone is secondarily lost in chondrichthyans^30,31^ (Fig. 4) is consonant with prior knowledge of the loss of perichondral bone in this same lineage^32^. Taken together, this has revived uncertainty about the true phylogenetic timing of the origin of endochondral ossification^33^. *Minjinia* provides direct corroboration for a more ancient origin.

*Minjinia* does not represent the first report of endochondral bone outside of Osteichthyes. However, it is by far the most extensive and unequivocal example and raises explicit questions in light of the proximity of *Minjinia* to the gnathostome crown (Fig. 4; Extended Data Figs. 3, 4). Isolated examples of trabecular endoskeletal bone have historically been reported in boreaspid osteostracans^20,34^, a rhenanid^35^, arthrodires^36^, a ptyctodont^37^, and a petalichthyid^38,39^. However, these reports are nearly all unillustrated statements; they have all been considered tenuous^3^ or dismissed as misidentifications^5^. In line with these assessments, we found no evidence of endochondral bone in material of *Buchanosteus* held in the Natural History Museum, London, or indeed in any other ‘placoderms’ we have examined. The *Epipetalichthys* holotype (Museum für Naturkunde, Berlin specimen MB.f.132.1-3) shows an apparently spongiose infilling in the anterior region of the braincase, but the identity of this structure, or even whether it is biological, cannot be determined. The *Epipetalichthys* tissue figured by Stensiö^38^ was very superficial, and possibly represents the retreat of perichondral bone deposited during cartilage growth^39^. Most recently, trabeculae in supposed endoskeletal pelvic bones of *Bothriolepis* have been termed endochondral bone^21^, although the small scale of these is in line with ‘superficial’ perichondral trabeculae seen elsewhere^38^. The reported examples in boreaspid osteostracans have also been dismissed by later authors^3,5^. Although they warrant further study, their tissue structures are unlikely to be homologous to osteichthyans owing to their phylogenetic remoteness and nested position in the Osteostraci^40^.

Among chondrichthyans, endochondral bone has been mentioned in ‘acanthodians’^3,41^ and superficial bone-like tissues have been reported in the skeletons of extant chondrichthyans. We are unable to substantiate statements about acanthodians: no authors have cited primary sources or specimens. One possible source is Watson’s^42^ description of “massive ossification” of the endoskeleton of *Diplacanthus*. However, our synchrotron data of this same specimen (Fig. 2) shows that this tissue is undoubtably calcified cartilage. Some authors have speculated that the superficial mineralised tissue in the jaws of acanthodians or chondrichthyans may have developed in an endochondral position^39^. Histological studies show that endoskeletal mineralization in the jaws of acanthodians is globular calcification and occasionally ‘sub-tessellate’^8,43^. Recent comparative histology and development in extant chondrichthyans has shown the presence of an extensive canalicular network in the tesserae^44^ and a trabecular tesseral network in some vertebral elements^45^, both resembling bone. Whether these represent homologues of osteichthyan examples remains open to debate; future works could employ synchrotron microtomography of stem-chondrichthyan cartilages to address these questions.

Does endochondral bone have a deep origin within the gnathostome stem group? This would imply repeated losses of this tissue. We do find statistical support for this hypothesis (Fig. 4, Table 1, Extended Data Figs. 5, 6, Supplementary Table 1), and the model is well justified on prior phylogenetic and biological grounds. Endochondral bone has long been known to be inconsistently developed across ‘primitive’ bony fishes: incomplete, polymorphic, or entirely absent ossification of the endoskeleton is known in both Palaeozoic actinopterygians^41,46,47^ and sarcopterygians{Cloutier:wm}, as well as more recent taxa^48^. The frequent absence of endochondral bone in osteichthyans is considered secondary, and other controlling factors such as body size, maturity, mechanical stress, and buoyancy can determine its degree of development^1^. Our findings are also in agreement with studies establishing a genetic basis for secondary loss of all bone types within chondrichthyans^49-51^, with the failure to produce endochondral bone likely representing arrested development of chondrocytes as opposed to a primary lack of ability^52^.

Another confounding factor in this question is the problem of ‘placoderm’ relationships. Although currently resolved in most analyses as a deeply pectinate grade along the gnathostome stem (Fig. 4), the backbone of this arrangement has poor statistical support, even in the present analysis (Extended Data Figs. 3). There is a lack of consistency in the arrangement of plesia and Bayesian tip-dating methods have even recovered a monophyletic Placodermi^53^. *Minjinia* itself highlights this uncertainty, given its highly unexpected character combinations. Notwithstanding its endochondral bone and crown-gnathostome-like inner ear structure, it resembles ‘acanthothoracids’—the ‘placoderms’ widely considered among the most removed from the gnathostome crown (i.e. most ‘primitive’): it possesses deep epaxial fossae either side of a prominent occipital ridge and a nuchal eminence otherwise seen only in acanthothoracids such as *Romundina*^18^ and *Arabosteus*^19^. This apparent character conflict could perhaps be more easily reconciled with a more coherent (though not necessarily monophyletic) ‘placoderm’ assemblage. Indeed, the highly pectinate structure of the ‘placoderm’ grade seems symptomatic of an overemphasis on characters and taxa resembling the crown group, thereby undersampling characters that could stabilise a clear picture of ‘placoderm’ interrelationships.

*Minjinia turgenensis* reveals new data on ‘placoderm’ endoskeleton and tissue diversity recorded from Mongolia—an otherwise extremely poorly known biogeographic realm for early gnathostomes. The phylogenetic placement of this ‘acanthothoracid’-like taxon crownward of all non-maxillate ‘placoderms’, in conjunction with possession of extensive endochondral bone, highlights the importance of material from traditionally undersampled geographic areas. The presence of endochondral bone renews the hypothesis that this tissue is evolutionarily ancient and was lost secondarily in chondrichthyans^6,33^. This view is overall consistent with evidence of generalised bone loss in chondrichthyans, potentially as a result of the suppression of bone-generating molecular genetic pathways^51,52^. Continued work in Mongolia and re-evaluation of phylogenetic datasets will be necessary to address this, with the results likely to lead to substantial re-evaluation of gnathostome phylogeny.

## Methods

### X-ray computed microtomography

We scanned MPC-FH100/9.1 using the Nikon XT 225s at the Museum of Paleontology, University of Michigan with the following parameters: 200kV, 140*µ*A, over 3123 projections and a voxel size of 32.92*µ*m. We conducted segmentation using Mimics 19.0 (http://biomedical.materialise.com/mimics; Materialise, Leuven, Belgium) and we imaged models for publication using Blender (https://www.blender.org).

### Synchrotron light propagation phase contrast tomography

We imaged *Diplacanthus crassismus* specimen NMS 1891.92.334 on Beamline 19 of the European Synchrotron Radiation Facility, using propagation phase-contrast synchrotron microtomography. We performed a spot scan with an energy of 116keV, achieving a voxel size of 0.55 *μ*m. We processed the resulting tomograms using VG StudioMax 2.2 (Volume Graphics, Germany), and prepared images in Blender.

### Phylogenetic analysis

We conducted a parsimony analysis using TNT 1.5^54^ and Bayesian analysis using MrBayes v 3.2.7^55^. The dataset consisted of 95 taxa and 284 discrete characters based on a pre-existing dataset^56^. We employed Osteostraci and Galeaspida as composite outgroups. We conducted parsimony analysis using both equal weights and implied weights methods. Global settings were 1000 search replicates and a hold of up to 1 million trees. Equal weights parsimony analyses were conducted using the ratchet with default settings. Implied weights parsimony used a concavity parameter of 3 and the search was without the ratchet. Command lists are included in Supplementary Information. We conducted Bayesian analysis using both a partitioned and unpartitioned dataset. We used the Mkv model^57^ and gamma rate distribution. We ran the analyses for 5 million generations with a relative burn-in fraction of 0.25. Runs were checked for convergence using Tracer^58^. We partitioned the dataset using a newly proposed method^28^ that partitions the data according to homoplasy levels. Using the results of implied weights parsimony conducted in TNT, we created a text table of character fit values. We wrote an R^59^ script to generate a list of partition commands for MrBayes.

We assessed parsimony ancestral states visually using Mesquite^60^. Likelihood and Bayesian ancestral states were estimated in R using the castor package^61^ version 1.5.7. Prior to calculating likelihood ancestral states on parsimony trees, we scaled branch lengths using PAUP*^62^ and calculated the likelihood scores for all of the trees under the Mkv model with gamma rate parameter. The trees were then exported with branch lengths. To account for overall uncertainty in tree estimates, we estimated ancestral states on 100 trees randomly selected from the fundamental set of most parsimonious trees and two times 50 trees selected from the 75% last trees of each posterior tree distribution from the Bayesian analysis. We then run an ancestral states estimation Mk model (using the castor R package) using both the Equal Rates (ER) and All Rates Different (ARD) models. This resulted in 400 ancestral states estimations. For each estimation we extracted the overlap log likelihood, the AIC (counting one parameter for the ER model and two for the ARD model) and the scaled log likelihood (probability) for the presence and absence of the endochondral bone character (character 4) for the last common node of *Minjinia* and crown-group gnathostomes. We present the median value of these distributions of the estimations overall log likelihoods, AICs and presence or absence of endochondral bone in Table 1.

## Data availability

The holotype specimen of *Minjinia turgenensis* will be permanently deposited in the collections of the Institute of Paleontology, Mongolian Academy of Sciences. Original tomograms are available at (doi:10.6084/m9.figshare.12301229) and rendered models are available at (doi:10.6084/m9.figshare.12301223). The phylogenetic character list and dataset are available as Supplementary Information S1 and S2. The LifeScience Identifier for *Minjinia turgenensis* is urn:lsid:zoobank.org:act:82A1CEEC-B990-47FF-927A-D2F0B59AEA87

## Code availability

R code for generating partitions based on character fits and code for likelihood ancestral states reconstructions and plots are available in the Supplementary Information.

## Acknowledgements

M. Bolortsetseg generously assisted MDB with contacts and field experience in Mongolia. Fieldwork was supported by National Geographic Society grants CRE 8769-10 and GEFNE35-12 to MDB. AJ’s field contributions were supported by funds from the Anna Maria Lundin’s stipend from Smålands Nation, Uppsala University. RS’s field contributions were supported by a Royal Society Research Grant and the University of Manchester. The majority of this work was supported by the European Research Council (ERC) under the European Union’s Seventh Framework Programme (FP/2007-2013)/ERC Grant Agreement number 311092 to MDB. RPD was also supported by the Île-de-France DIM (domaine d’intérêt majeur) matériaux anciens et patrimoniaux grant PHARE. Stig Walsh is thanked for access and loan of specimen at the National Museums of Scotland. Synchrotron tomography was performed at the ESRF (application LS 2451) with the assistance of Paul Tafforeau. SG was supported by a Royal Society Dorothy Hodgkin Research Fellowship. Matt Friedman is thanked for undertaking the X-ray computed microtomography analysis. This study includes data produced in the CTEES facility at University of Michigan, supported by the Department of Earth & Environmental Sciences and College of Literature, Science, and the Arts. TNT was made available with the support of the Willi Hennig Society.

## Author Contributions

MDB conceived and designed the study. MDB, AJ, YAA, and EZ participated in all field seasons. RPD and AJ undertook preliminary CT scanning and segmentation that revealed the fossil was a ‘placoderm’ and had endochondral bone. RS discovered the first vertebrate remains in the first field season at Yamaat Gol in 2010. SG undertook the segmentation of *Minjinia* with input from MDB. AJ performed segmentation of *Diplacanthus* tissue. MC provided input on occipital comparative morphology of ‘placoderms’. RPD provided data and comparative analyses and data for endoskeletal tissue. YAA provided background on the geology, palaeontology, and stratigraphy of the type location; EZ and YAA organized field logistics and permitting. MDB, SG, MC, RPD, and AJ undertook the anatomical interpretation and prepared the figures. MDB and SG conducted the phylogenetic analyses. RS conducted the parsimony branch support analyses. TG wrote the script for generating MrBayes partitions from TNT’s character fits table and conducted the likelihood and model-fitting analyses. The manuscript was written by MDB, RPD, and SG.

---

**Extended Data Fig 1.**
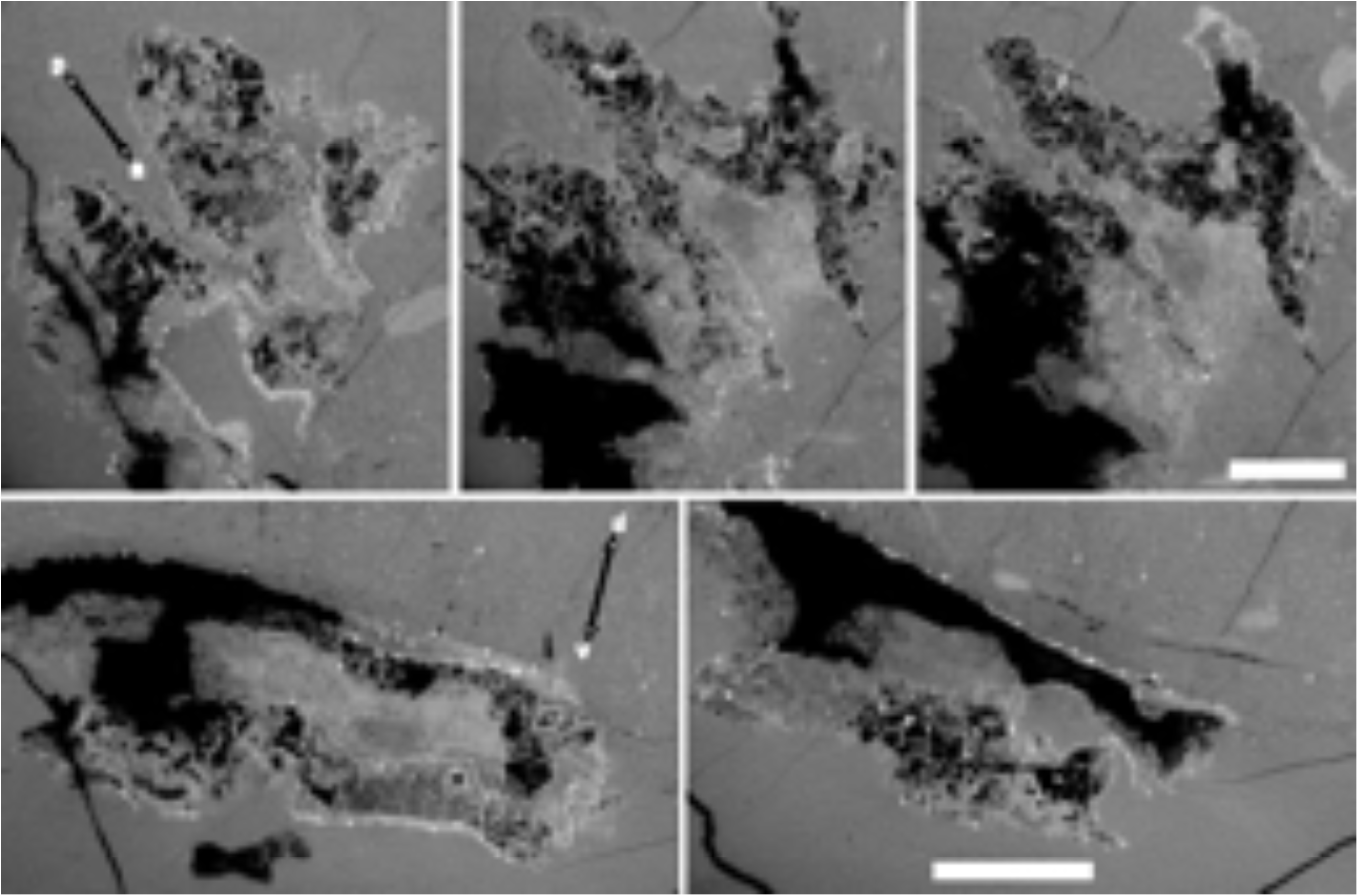
Tomograms of endoskeletal ossification in *Minjinia*. Top row: semi-coronal sections through braincase. Double-headed arrows indicate anterior-posterior (a-p) dorsal-ventral (d-v) axes. Bottom row: semi-transverse sections through posterior part of endocranium. Voids of black space represent mouldic preservation. Scale bars, 10 mm and apply across each row of panels.

**Extended Data Fig 2.**
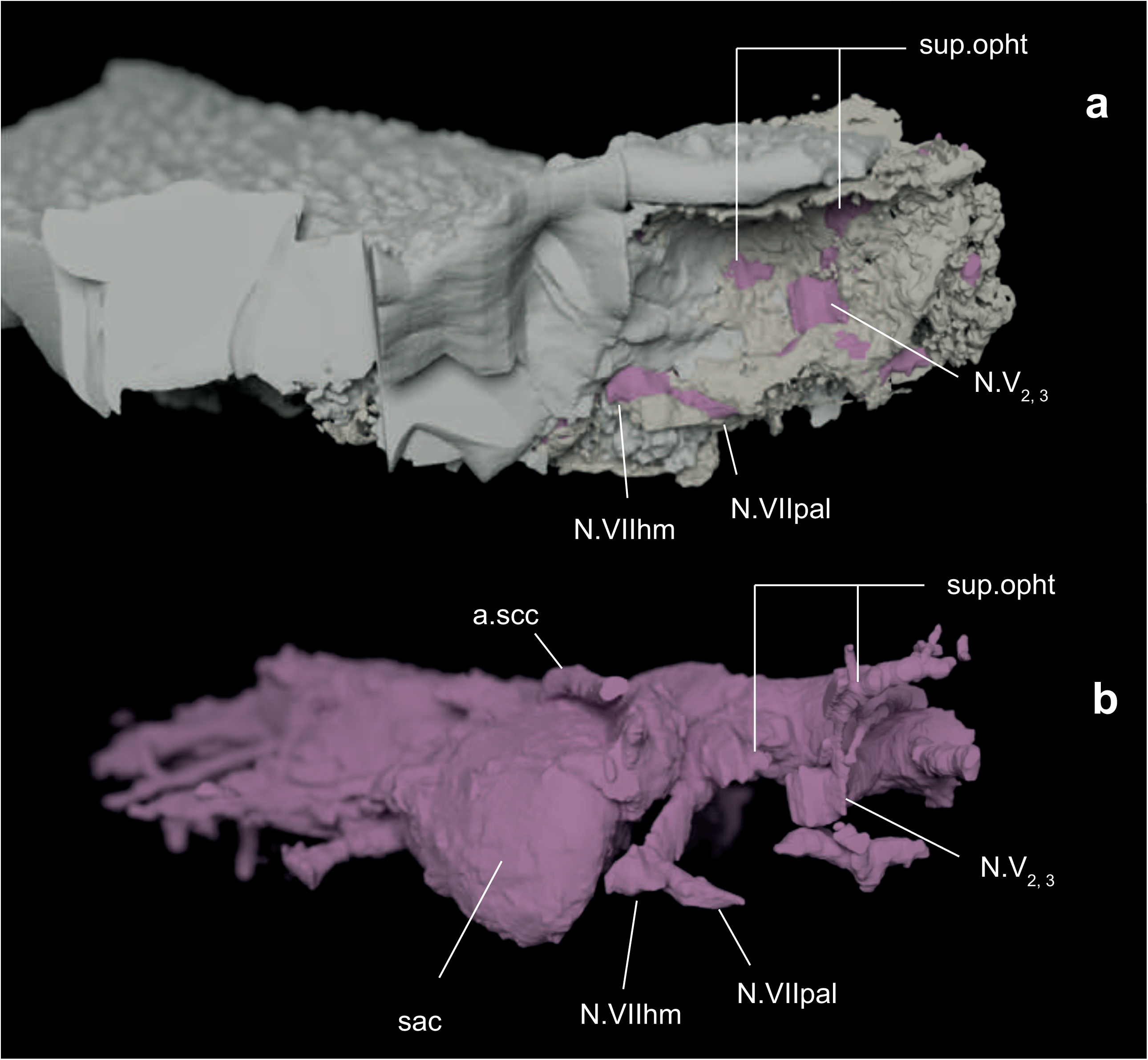
Right orbital wall and innervation pattern of *Minjinia*. **a**, orbit in anterolateral view showing disposition of nerve openings (pink infill). **b**, endocast in the same perspective showing the relationship between nerve canals and endocast. a.scc., anterior semicircular canal; N.V_2,3_ trunk of the trigeminal nerve canal for branches 2 and 3; N.VIIhm, hyomandibular branch of facial nerve canal; N.VIIpal, palatine branch of facial nerve canal; sac., sacculus; sup.opth, canal for supra-ophtalmic nerve. Scale bars, 20 mm (upper scale bar associates with **a**, lower scale bar associates with **b** and **c**).

**Extended Data Fig 3.**
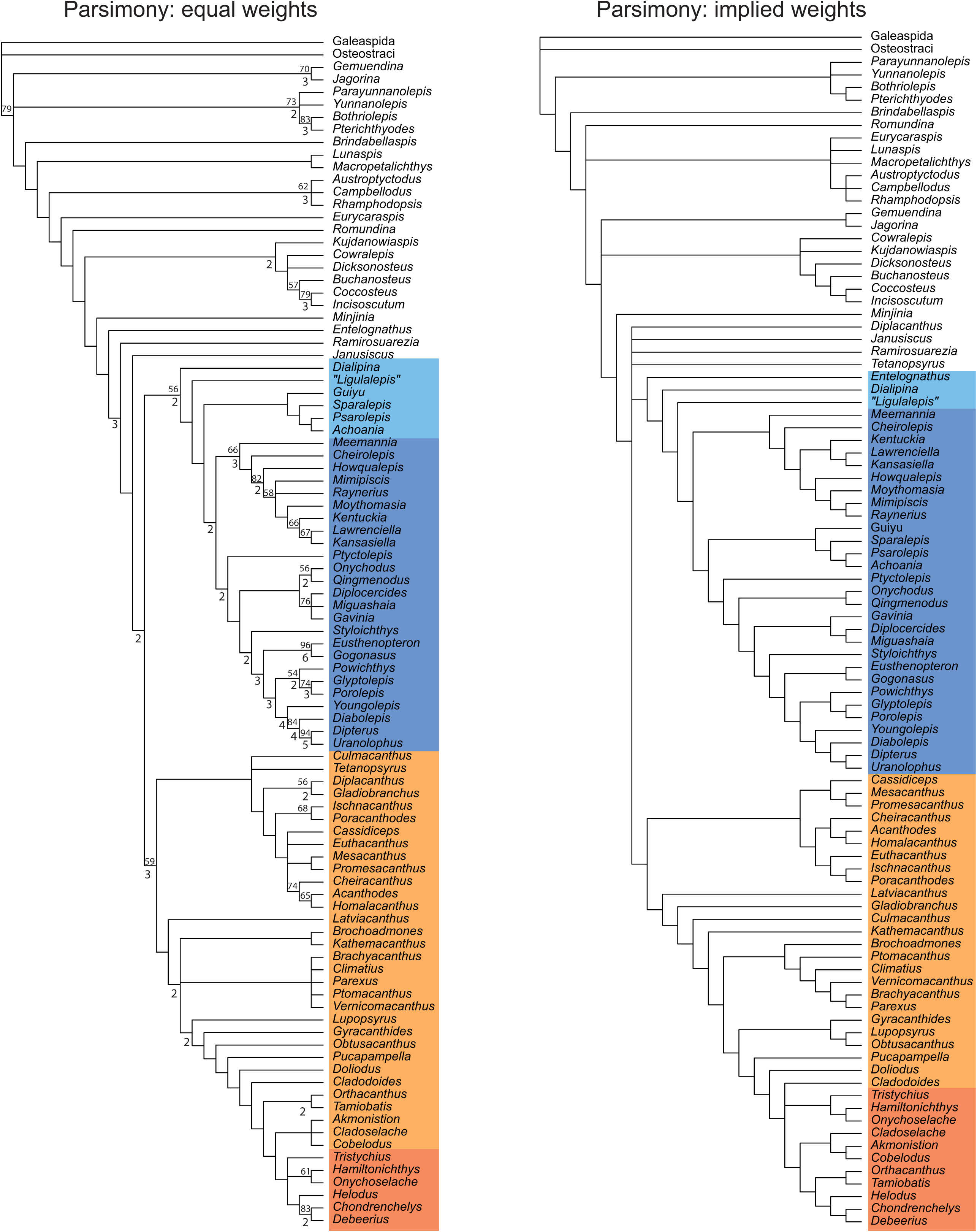
Results of phylogenetic parsimony analysis. Dataset consists of 95 taxa and 284 characters. Both trees are strict consensus topologies. Equal weights parsimony analysis using the ratchet resulted in 240 trees with a length of 831 steps. Implied weights parsimony analysis using random addition sequence + branch-swapping resulted in 8 optimal trees with score 85.20513. Double-digit figures above internal branches are bootstrap values of 50% and over; single-digit figures below branches are Bremer decay index values. Blue shading: osteichthyan total group (dark blue: crown group); orange shading: chondrichthyan total group (dark orange: crown group).

**Extended Data Fig 4.**
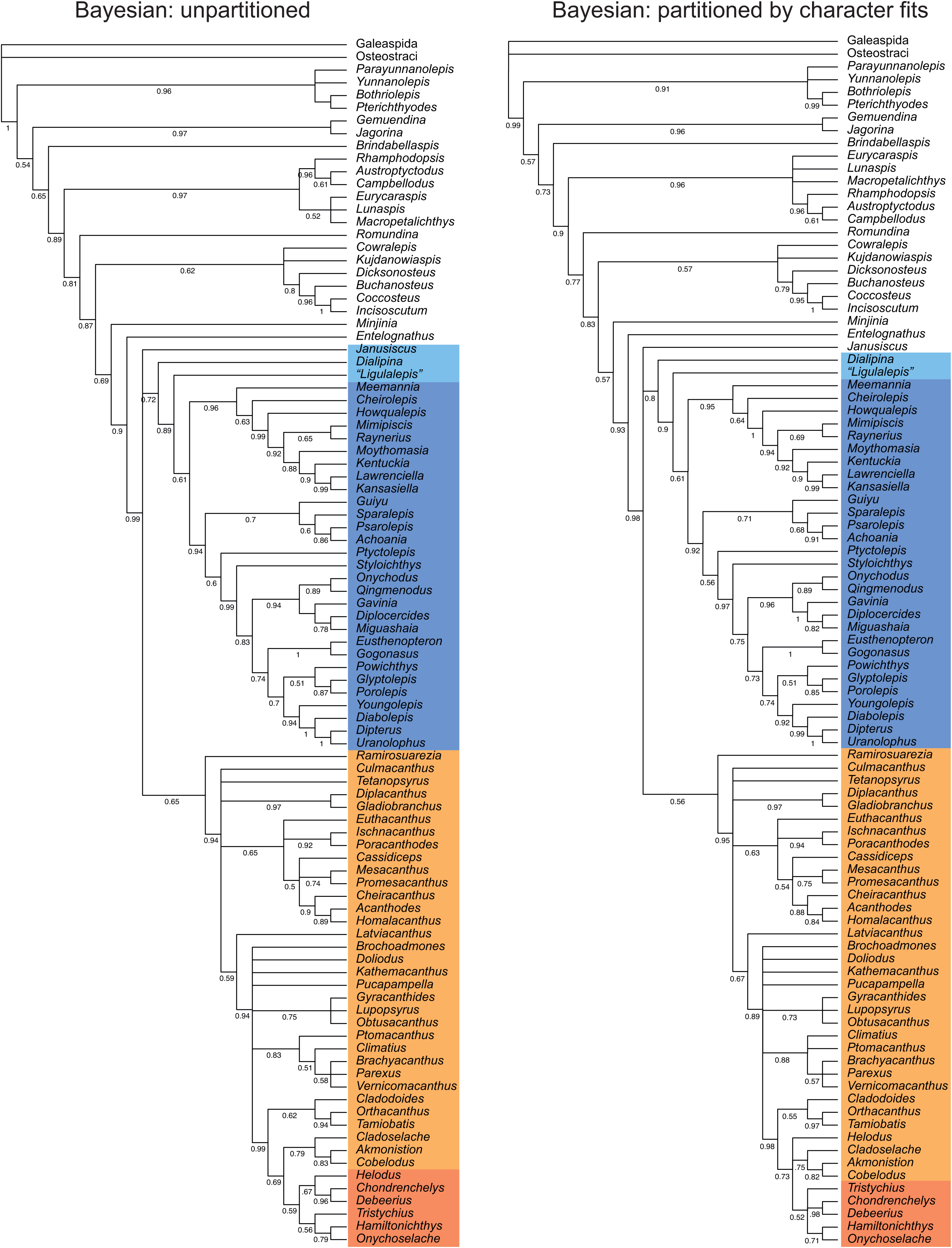
Results of Bayesian phylogenetic analysis using both partitioned and unpartitioned data. Majority-rules consensus trees with posterior probabilities shown along branches. Blue shading: osteichthyan total group (dark blue: crown group); orange shading: chondrichthyan total group (dark orange: crown group).

**Extended Data Fig 5.**
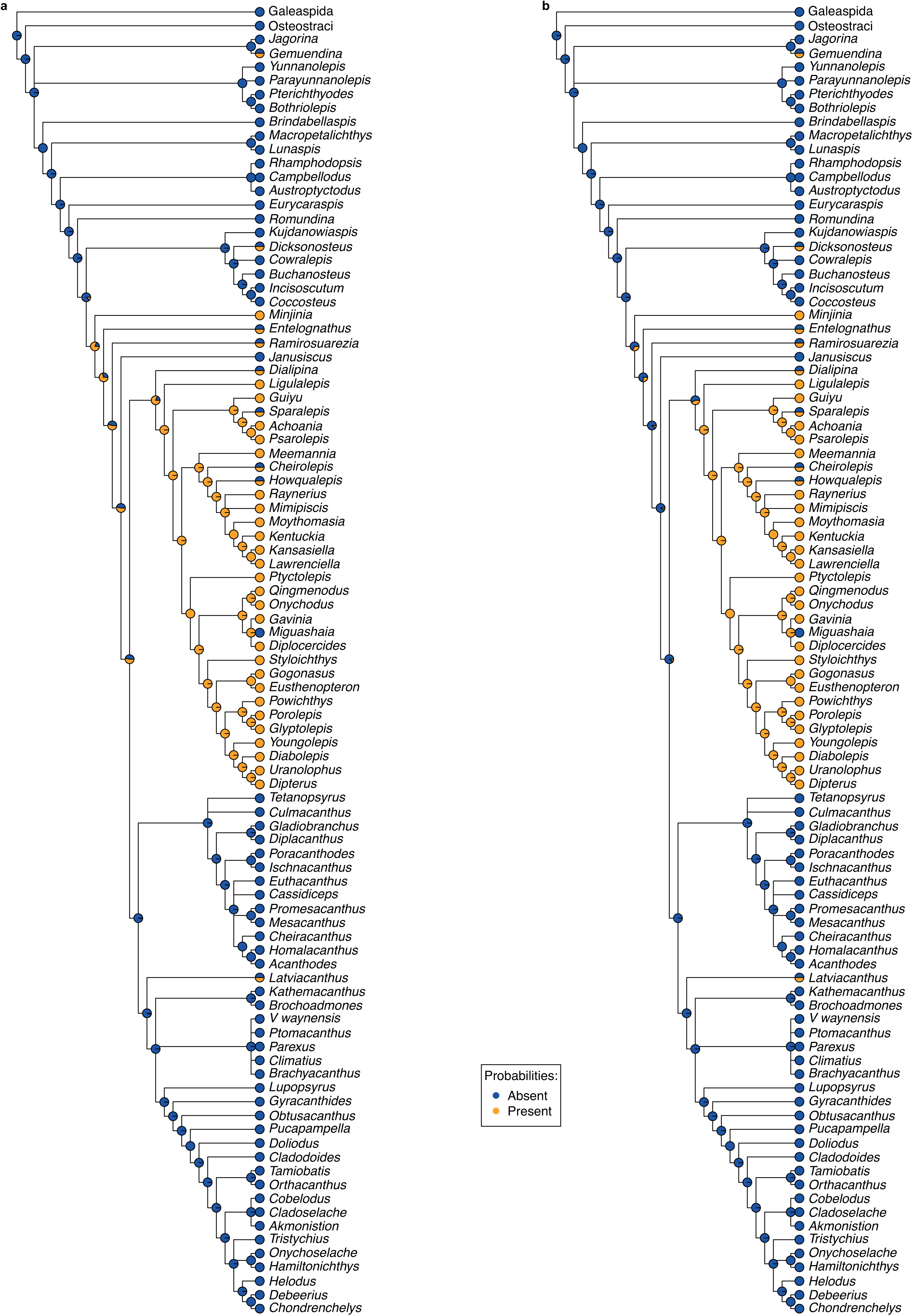
Likelihood ancestral state mapping of endochondral bone on equal weights parsimony results. **a**, ARD, all rates different model; **b**, ER, equal rates model.

**Extended Data Fig 6.**
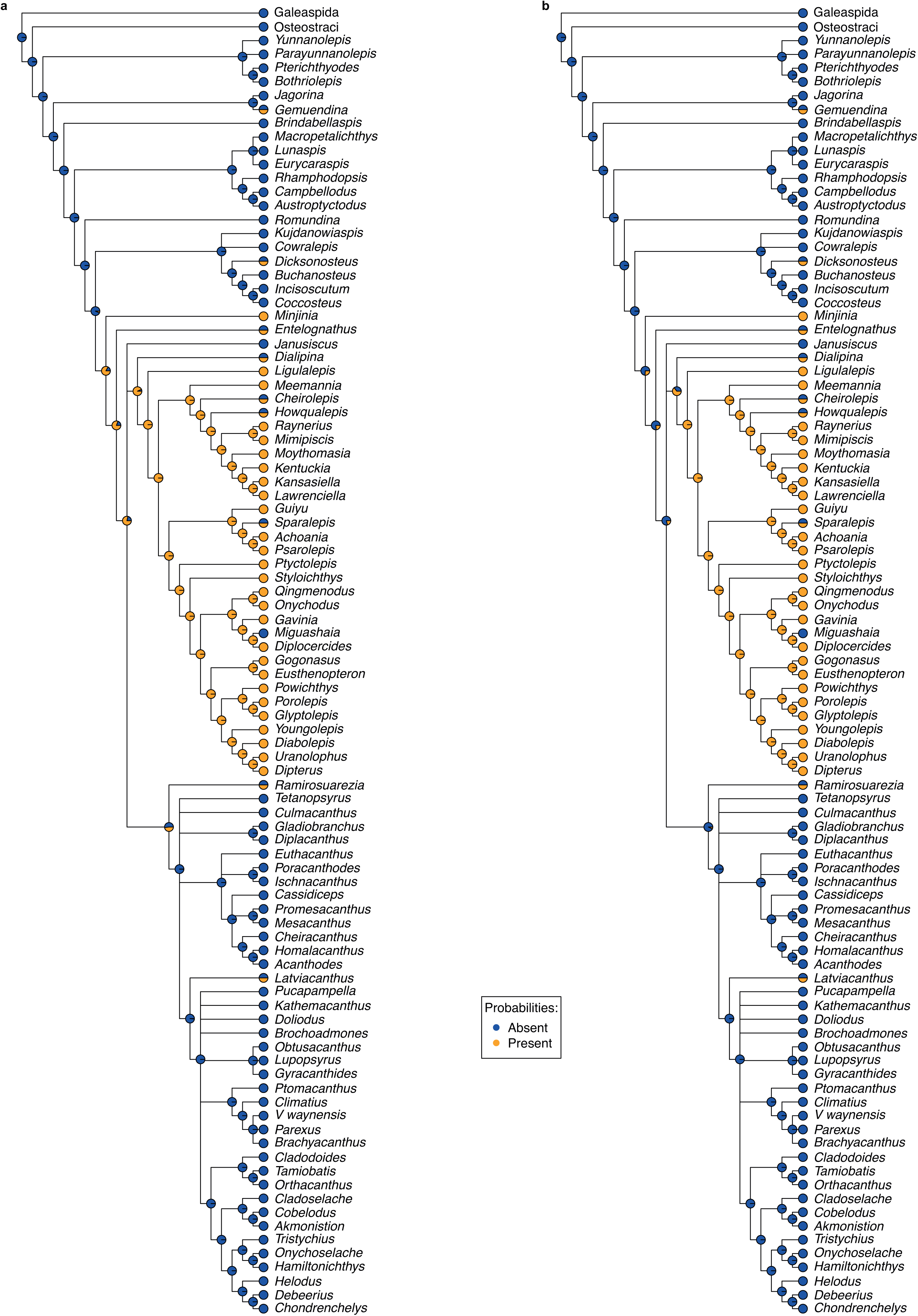
Likelihood ancestral state mapping of endochondral bone on unpartitioned Bayesian analysis results. **a**, ARD, all rates different model; **b**, ER, equal rates model.

