## Supplementary Information for "Endochondral bone in an Early Devonian ‘placoderm’ from Mongolia"

**Supplementary Information Guide**

**Supplementary Information.** Full phylogenetic character list and notes; character-state optimization data for a fully-resolved parsimony tree; Supplementary Table; Supplementary References.

**Supplementary Data 1.** Phylogenetic character data (master Nexus file), command files, R scripts for data partitions, and command lists for phylogenetic analyses in TNT.

**Supplementary Data 2.** R scripts and markdown for generating figures and output tables for ancestral states using maximum likelihood.

**Supplementary Video 1.** Transverse tranche through otic wall of *Minjinia turgenensis* showing endochondral bone corresponding to Fig 1a in the main text. Note that the thickness of the trabeculae is exaggerated: the trabeculae are encrusted in a carbonaceous matrix that does not contrast strongly with bone. This resulted in difficulties separating the trabeculae from the matrix with a single consistent threshold value. Indigo coloured material is cranial exoskeleton. Taupe coloured material is endoskeletal bone.

**Supplementary Video 2.** Transverse tranche through otic wall of *Ligulalepis* showing endochondral bone corresponding to Fig 1c in the main text. Data are from Clement et al. (2018). Indigo coloured material is cranial exoskeleton. Taupe coloured material is endoskeletal bone.
