## Supplementary Data 1 for "Endochondral bone in an Early Devonian ‘placoderm’ from Mongolia"

#### Phylogenetic character list

The character list derives primarily from Clement et al. [1] with some additions. To facilitate interpretation of characters, we have retained descriptive text and extended to them or added descriptions where relevant. Notes from Clement et al. are in parentheses and references within parenthetical text refer to citations in their paper

**1. Tessellate prismatic calcified cartilage.**
0 absent
1 present

**2. Prismatic calcified cartilage.** *Culmacanthus*, *Eurycaraspis*, *Gemuendina*, *Lunaspis*, *Ptomacanthus*, *Ramirosuarezia*, *Rhamphodopsis*, *Ramirosuarezia* changed from inapplicable to missing as all these taxa are scored as missing data for the principal character (absence or presence of calcified cartilage).

0 single layered
1 multi-layered

**3. Perichondral bone.**
0 present
1 absent

**4. Extensive endochondral ossification.**
0 absent
1 present

**5. Enamel(oid) present on dermal bones and scales.** Enameloid is reported in osteostracans but was previously scored polymorphic. Enameloid is well described in trematаspid osteostracans[2-4]. However, this group is highly nested within Osteostraci. The histology of the outer branches of Osteostraci is less clear but appears to be absent in fragments assigned to *Ateleaspis* by Märss et al. [5]. Based on this and arguments from previous workers[6], we have scored this character as absent. *Debeerius* and *Orthacanthus* changed from '-' to '?' to reflect known character hierarchy.

0 absent
1 present

**6. Enamel.** *Brachyacanthus* changed to '-' from '?' in order to be consistent with codings for enameloid. Changed to '-' for Osteostraci. *Onychoselache* set to '?'

0 single-layered
1 multi-layered

**7. Enamel layers.** *Brachyacanthus* changed to '-' from '?' in order to be consistent with codings for enameloid. *Onychoselache* set to '?'. *Guiyu* has been re-coded as '?' because it is unclear from published figures if the enamel layers are closely appressed.

0 applied directly to one another (ganoine)
1 separated by layers of dentine

**8. Pore canal network** [7]. The previous formulation of this character referenced an extensive pore canal network. However, this leaves assessment of presence and absence ambiguous based on subjective interpretations of when such a network might be considered 'extensive'. Instead, we have reformulated this character to capture those instances in which cavities connect with the outer surface of the bone via pores. Changed from '?' to '0' for *Cassidiceps*, *Euthacanthus*, *Helodus*, *Homalacanthus*, *Ischnacanthus*, *Janusiscus*, *Kathemacanthus*, *Kentuckia*, *Latviacanthus*, '*Ligualepis*', *Mesacanthus*, *Onychodus*, *Pterichthyodes*, *Tristychius*; from '1' to '0' in *Dialipina*, *Poracanthodes*; from '0' to '?' in *Guiyu*, *Jagorina*, *Moythomasia*). *Debeerius* and *Orthacanthus* have been recoded from inapplicable to '?' consistent with the scorings for other chondrichthyan-like taxa. This also allows this character to be used as a root in hierarchies of other characters. This character could be made contingent on the presence/absence of dermal bones, in which case the '?' can be inferred as inapplicable if a prior hierarchy is specified.

0 absent
1 present

**9. Dentinous tissue**
0 absent
1 present

**10. Dentine kind**
0 mesodentine
1 semidentine
2 orthodentine

**11. Bone cell lacunae in body scale bases**
0 present
1 absent

**12. Main dentinous tissue forming fin spine.** Changed from '-' to '0' in *Pterichthyodes*. *Austroptyctodus* changed from '-' to '?'.

0 osteodentine
1 orthodentine

**13. Longitudinal scale alignment in fin webs.** Changed from '-' to '0' in *Pterichthyodes*. Changed from inapplicable to '0' in *Parayunnanolepis* and to '?' in *Bothriolepis*, *Yunnanolepis*, and Galeaspida. *Miguashaia* is changed from unknown to present. *Akmonistion*, *Chondrenchelys*, *Cladoselache*, *Cobelodus*, *Debeerius*, *Onychoselache*, *Orthacanthus*, *Tristychius*, have been recoded from absent to '?'. *Cheirolepis* changed from absent to present.

0 absent
1 present

**14. Differentiated lepidotrichia.** This character is contingent on longitudinal alignment of fin web scales. All codings in this character have been re-scored to reflect this contingency.

0 absent
1 present

**15. Body scale growth pattern**
0 comprising single odontode unit/generation ("monodontode")
1 comprising a complex of multipe odontode generations/units ("polyodontode")

**16. Body scale growth concentric**
0 absent
1 present

**17. Generations of odontodes**
0 buried
1 areally growing
2 resorbed

**18. Body scales with peg-and-socket articulation** (Changed from '?' to '1' in *Styloichthys*.). *Botrhiolepis* changed from inapplicable to unknown. *Bothriolepis* body scales do exist [8] but are poorly known.

0 absent
1 present

**19. Scale peg** (Changed from '?' to '0' in Styloichthys.)
0 broad
1 narrow

**20. Anterodorsal process on scale** (Changed from '?' to '1' in *Styloichthys*.)
0 absent
1 present
**21. Body scale profile** (Changed from '?' to '1' in *Styloichthys*.). Osteostraci changed from '?' to '1' (flattened).

0 distinct crown and base demarcated by a constriction ("neck")
1 flattened

**22. Profile of scales with constriction between crown and base**
0 neck similar in width to crown
1 neck greatly constricted, resulting in anvil-like shape

**23. Body scales with bulging base** (Changed from '?' to '0' in *Styloichthys*.)
0 absent
1 present

**24. Body scales with flattened base** (Changed from '?' to '1' in *Styloichthys*.)
0 present
1 absent

**25. Basal pore in scales**
0 absent
1 present

**26. Flank scale alignment** (Changed from '-' to '?' for *Chondrenchelys*).
0 vertical rows oblique rows or hexagonal
1 rhombic packing
2 disorganised

**27. Scute-like ridge scales (basal fulcra)**
0 absent
1 present

**28. Sensory line canal**
0 perforates scales
1 passes between scales
2 C-shaped scales

**29. Dermal ornamentation** (Changed from '1' to '3' for *Glyptolepis*, from '3' to '-' for *Onychoselache*, from '3' to '0' for *Psarolepis*).

0 smooth
1 parallel, vermiform ridges
2 concentric ridges
3 tuberculate

**30. Sensory line network.** *Rhamphodopsis* has been changed from 0 to 1.
0 preserved as open grooves (sulci) in dermal bones

1 sensory lines pass through canals in dermal bones (open as pores)

**31. Sensory canals/grooves**
0 contained within the thickness of dermal bones
1 contained in prominent ridges on visceral surface of bone

**32. Jugal portion of infraorbital canal joins supramaxillary canal**
0 present
1 absent

**33. Dermal skull roof**
0 includes large dermal plates
1 consists of undifferentiated plates or tesserae

**34. Anterior pit line of dermal skull roof** (Changed from ‘1’ to ‘0’ for *Cheirolepis*).
0 absent
1 present

**35. Tessera morphology**
0 large interlocking polygonal plates
1 microsquamose, not larger than body squamation

**36. Cranial spines.** This and the following character have been expanded from the original multistate formulation in Clement et al. (2018).
(0) absent
(1) present

**37. Cranial spines**
(0) monocuspid
(1) multicupsid

**38. Extent of dermatocranial cover**
0 complete
1 incomplete (limited to skull roof)

**39. Openings for endolymphatic ducts in dermal skull roof.** Score for *Gemuendina* changed from “?” to present, based on Gross (1963). Score changed from absent to present for *Eurycaraspis* following reinterpretation of Pan et al. (2015). Score changed from ‘?’ to “0’ in *Climatius* [9]. Score changed from ‘-’ to ‘?’ in *Ischnacanthus, Obtusacanthus, Poracananthodes, Promesacanthus, Tetanopsyrus and Vernicomacanthus*.
0 present
1 absent

**40. Endolymphatic ducts with oblique course through dermal skull bones**
0 absent
1 present

**41. Endolymphatic duct relationship to median skull roof bone (i.e. nuchal plate)**
0 within median bone
1 on bones flanking the median bone (e.g. paranuchals)

**42. Pineal opening perforation in dermal skull roof**

0 present
1 absent

**43. Dermal plate associated with pineal eminence or foramen.** *Buchanosteus* and *Entelognathus* changed to 0 consistent with their descriptions; *Kujdanowiaspis* changed to 'unknown'. Rhenanids can only be scored as '?' because the nature of the underlying dermal bones is usually unknown.

0 contributes to orbital margin (plate(s) excluded from orbital margin by skull roofing bones.)
1 plate bordered laterally by skull roofing bones

**44. Broad supraorbital vaults**

0 absent
1 present

**45. Median commissure between supraorbital sensory lines.** Changed from '0' to '1' for *Eurycaraspis*.

0 absent
1 present

**46. Dermal cranial joint at level of sphenoid-otic junction** (Changed from '?' to '-' for Chondrenchelys.)

0 absent
1 present

**47. Otic canal extends through postparietals**

0 absent
1 present

**48. Number of bones of skull roof lateral to postparietals** (We have modified this character to take into account the condition in *Powichthys* and lungfishes (new additions to the matrix), which show more than two bones lateral to the postparietals.)

0 two
1 one
2 more than two

**49. Suture between paired skull roofing bones (centrals of placoderms postparietals of osteichthyans)**

0 straight
1 sinusoidal

**50. Medial processes of paranuchal wrapping posterolateral corners of nuchal plate.** We have reformulated this as a binary character. The previous version of this character in Clement et al. (2018) added a state for preclusion of paranuchals from nuchals, but we consider this unnecessary: the key variable concerns processes that wrap the posterior margin of the medially bordering bone.

0 absent
1 present

**51. Paired pits on ventral surface of nuchal plate**
0 absent
1 present

**52. Sclerotic ring**
0 absent
1 present

**53. Consolidated cheek plates**
0 absent
1 present

**54. Cheek plate**
0 undivided
1 divided (i.e., squamosal and preopercular)

**55. Subsquamosals in taxa with divided cheek** (Changed from '?' to '-' for *Chondrenchelys*).
0 absent
1 present

**56. Preopercular shape** (Changed from '?' to '-' for *Chondrenchelys*).
0 rhombic
1 bar-shaped

**57. Vertical canal associated with preopercular/suborbital canal**
0 absent
1 present

**58. Enlarged postorbital tessera separate from orbital series**
0 absent
1 present

**59. Extent of maxilla along cheek**
0 to posterior margin of cheek
1 cheek bones exclude maxilla from posterior margin of cheek

**60. Dermal neck joint.** *Jagorina* changed to '-'. *Lunaspis* codings based on attributed material (MB.f.1309.1-2) and radiographs in the Senckenberg Museum's online catalogue (Röntgenarchiv Wilhelm Stürmer: 10825-WS 10825).
0 overlap
1 ginglymoid ('arthrodire'-type)
2 reverse ginglymoid ('antiarch'-type)
3 longitudinal

**61. Sensory line scales/plates on head**
0 unspecialized
1 apposed growth
2 paralleling canal

3 semicylindrical C-shaped ring scales

**62. Bony hyoidean gill-cover series (branchiostegals)**
0 absent
1 present

**63. Branchiostegal plate series along ventral margin of lower jaw**
0 absent
1 present

**64. Branchiostegal ossifications.** Ordered.
0 plate-like
1 narrow and ribbon-like
2 filamentous

**65. Branchiostegal ossifications.** (Changed from '1' to '0' for *Porolepis*, *Gogonasus*).
0 ornamented
1 unornamented

**66. Imbricated branchiostegal ossifications**
0 absent
1 present

**67. Median gular**
0 absent
1 present

**68. Lateral gular**
0 absent
1 present

**69. Opercular (submarginal) ossification**
0 absent
1 present

**70. Shape of opercular (submarginal) ossification**
0 broad plate that tapers towards its proximal end
1 narrow, rod-shaped

**71. Size of lateral gular plates**
0 extending most of length of the lower jaw
1 restricted to the anterior third of the jaw (no longer than the width of three or four branchiostegals)

**72. Gill arches**. (Changed from '?' to '0' in *Chondrenchelys*).
0 largely restricted to region under braincase
1 extend far posterior to braincase

**73. Basihyal**
0 absent

1 present

**74. Interhyal**
0 absent
1 present

**75. Hypohyal**
0 absent
1 present

**76. Endoskeletal urohyal**
0 absent
1 present

**77. Oral dermal tubercles borne on jaw cartilages or at margins of the mouth.** Galeaspida and Osteostraci changed from inapplicable to 0 to allow use of character hierarchies. This is also justifiable in standard non-hierarchical analyses as there is no requirement that jaws precede dentitions (though it seems functionally likely to have done so). There is no necessary dependence between dentition and jaws [10].
0 absent
1 present

**78. Oral dermal tubercles patterned in organised rows (teeth)**
0 absent
1 present

**79. Enamel(oid) on teeth**
0 absent
1 present

**80. Cap of enameloid restricted to upper part of teeth (acrodin)**
0 absent
1 present

**81. Tooth whorls**
0 absent
1 present

**82. Bases of tooth whorls**
0 single, continuous plate
1 some or all whorls consist of separate tooth units

**83. Distribution of tooth whorls** (Changed from '1' to '0' for *Debeerius*; from '-' to '0' for *Chondrenchelys*.)
0 entire length of tooth row
1 restricted to symphysial region

**84. Distribution of tooth whorls.** Ordered. (Changed from '?' to '0' for *Helodus*) State symbols for this character are reassigned and treated as ordered as the previous unordered arrangement violates the triangle inequality. A transition from present in lower only to present in upper only will always require two events: loss of lower plus gain of upper (the converse is also true). Thus, these characters must be ordered. Note that ordering imposes no constraint on which state appears first.

0 lower jaws only
1 upper and lower jaws
2 upper jaws only

**85. Teeth ankylosed to dermal bones**

0 absent
1 present

**86. Plicidentine**

0 absent
1 present

**87. Dermal jaw plates on biting surface of jaw cartilages**

0 absent
1 present

**88. Maxillary and dentary marginal bones of mouth**

0 absent
1 present

**89. Premaxilla**

0 extends under orbit
1 restricted anterior to orbit

**90. Maxilla shape**

0 splint-shaped
1 cleaver-shaped

**91. Pair of tooth plates (anterior supragnathals or vomers) on ethmoidal plate** (Changed from ‘1’ to ‘0’ for *Onychodus*, *Chondrenchelys*.)

0 absent
1 present

**92. Strong posterior flexion of dentary symphysis** (Changed from ‘?’ to ‘-’ for *Chondrenchelys*.)

0 absent
1 present

**93. Extent of infradentaries**

0 along much of ventral margin of dentary
1 restricted to posterior half of dentary

**94. Coronoid fangs**

0 absent
1 present

**95. Position of upper mandibular arch cartilage (and associated cheek plate where present)**
0 entirely suborbital
1 with a postorbital extension

**96. Position of mandibular arch articulations** (Changed from ‘0’ to ‘1’ for *Cheirolepis*; ‘0’ to ‘?’ in *Chondrenchelys*, *Debeerius*.)
0 terminal
1 subterminal

**97. Autopalatine and quadrate** (Changed from ‘-’ to ‘0’ for *Debeerius*.)
0 comineralized
1 separate mineralizations

**98. Large otic process of the palatoquadrate** (Changed from ‘1’ to ‘?’ for *Chondrenchelys*.)
0 absent
1 present

**99. Insertion area for jaw adductor muscles on palatoquadrate**
0 ventral or medial
1 lateral

**100. Palatoquadrate fused with neurocranium**
0 absent
1 present

**101. Oblique ridge or groove along medial face of palatoquadrate**
0 absent
1 present

**102. Fenestration of palatoquadrate at basipterygoid articulation** (Changed from ‘0’ to ‘-’ for *Chondrenchelys*.)
0 absent
1 present

**103. Perforate or fenestrate anterodorsal (metapterygoid) portion of palatoquadrate**
0 absent
1 present

**104. Pronounced dorsal process on Meckelian bone or cartilage**
0 absent
1 present

**105. Number of coronoids**
0 four or more
1 three or fewer

**106. Preglenoid process** (Changed from ‘?’ to ‘1’ for *Helodus*.)
0 absent

1 present

**107. Jaw articulation located on rearmost extremity of mandible**
0 absent
1 present

**108. Precerebral fontanelle**
0 absent
1 present

**109. Median dermal bone of palate (parasphenoid)**
0 absent
1 present

**110. Parasphenoid**
0 lozenge-shaped
1 splint-shaped
2 diamond-shaped

**111. Multifid anterior margin of parasphenoid denticle plate**
0 absent
1 present

**112. Enlarged ascending processes of parasphenoid**
0 absent
1 present

**113. Buccohypophysial canal in parasphenoid**
0 single
1 paired

**114. Nasal opening(s)**
0 dorsal, placed between orbits
1 ventral and anterior to orbit

**115. External opening of posterior nostril and orbit**
0 separated by dermal bone(s)
1 confluent

**116. Olfactory tracts**
0 short, with olfactory capsules situated close to telencephalon cavity
1 elongate and tubular (much longer than wide)

**117. Prominent pre-orbital rostral expansion of the neurocranium.** We have restructured this character to describe three different rostrum constructions that can be individually explained as any combination of two transformations (however, there may be substantial differences in the complexity of these transformations). The purpose is to account for the fact that macropetalichthyids and a number of other placoderms have a prominent pre-orbital expansion of the braincase, covered by dermal bones. In macropetalichthyids, this rostrum contains the nasal capsules, whereas in ‘acanthothoracids’ and antiarchs, the nasal capsules remain nestled between the orbits. The rostrum is instead formed from the 'premedian' or 'upper lip' area (though the upper lip is pushed forward in macropetalichthyids as well) [11].

0 present, formed of subethmoidal platform ('upper lip')
1 absent
2 present, formed of rhinocapsular block

**118. Pronounced sub-ethmoidal keel**

0 absent
1 present

**119. Internasal vacuities**

0 absent
1 present

**120. Discrete division of the ethmoid and more posterior braincase at the level of the optic tract canal**

0 absent
1 present

**121. Position of myodome for superior oblique eye muscles**

0 posterior and dorsal to foramen for nerve II
1 anterior and dorsal to foramen

**122. Endoskeletal intracranial joint**

0 absent
1 present

**123. Spiracular groove on basicranial surface**

0 absent
1 present

**124. Transverse otic process**

0 present
1 absent

**125. Jugular canal** (In the previous formulation, a 'short' jugular canal was described as being anterior to the skeletal labyrinth. However, most taxa with a short jugular canal have this somewhere along the length of the labyrinth rather than exclusively anterior to it. We have therefore modified the terminology to accommodate this pattern. Changed from '?' to '2' for *Chondrenchelys*.). Ordered character.

0 long (invested in otic region along length of skeletal labyrinth)
1 short (restricted to short portion of region of skeletal labyrinth, or anterior to it)
2 absent (jugular vein uninvested in otic region)

**126. Spiracular groove on lateral commissure**

0 absent
1 present

**127. Subpituitary fenestra** (*Onychodus* changed from '?' to '0'.). Changed to absent in *Macropetalichthys*[12].

0 absent
1 present

**128. Supraorbital shelf broad with convex lateral margin**
0 absent
1 present

**129. Orbit dorsal or facing dorsolaterally, surrounded laterally by endocranium.** *Gemuendina* and *Jagorina* are changed to 'missing' based on arguments from Goujet[13] and new work on rhenanid-like fishes from the Hunsrück Slate. Goujet suggested that the hyomandibula in these taxa was, in fact, a hypertrophied lateral otic process ('anterior postorbital process'). Forthcoming work by M.D.B, M.C., and Matt Friedman provides some modest corroboration for this hypothesis.
0 present
1 absent

**130. Eyestalk attachment area**
0 absent
1 present

**131. Postorbital process** (Changed from '?' to '1' for *Chondrenchelys*.)
0 absent
1 present

**132. Canal for jugular in postorbital process** (Changed from '?' to '1' for *Chondrenchelys*.)
0 absent
1 present

**133. Series of perforations for innervation of supraorbital sensory canal in supraorbital shelf**
0 absent
1 present

**134. Extended prehypophysial portion of sphenoid**
0 absent
1 present

**135. Narrow interorbital septum, with outer walls in contact along midline forming a single sheet.** (The previous formulation of this character distinguished 'narrow' and 'broad' interorbital septa. This lacks precision, and is subject to differing opinions. We therefore have reformulated this character to refer to distinguish cases where the lateral walls of the braincase are separate in the orbital region from those where they join as a single sheet along the midline.)
0 absent
1 present

**136. The main trunk of facial nerve (N. VII)**
0 elongate and passes anterolaterally through orbital floor
1 stout, divides within otic capsule at the level of the transverse otic wall

**137. Course of hyoid ramus of facial nerve (N. VII) relative to jugular canal**
0 traverses jugular canal, with separate exit in otic region
1 intersects jugular canal, with exit through posterior jugular foramen

**138. Glossopharyngeal nerve (N. IX) exit**
0 foramen situated posteroventral to otic capsule and anterior to metotic fissure
1 through metotic fissure

**139. Relationship of cranial endocavity to basisphenoid**
0 endocavity occupies full depth of sphenoid
1 enodcavity dorsally restricted

**140. Subcranial ridges**
0 absent
1 present

**141. Ascending basisphenoid pillar pierced by common internal carotid**
0 absent
1 present

**142. Canal for lateral dorsal aorta within basicranial cartilage**
0 absent
1 present

**143. Entrance of internal carotids**
0 through separate openings flanking the hypophyseal opening or recess
1 through a common opening at the central midline of the basicranium

**144. Canal for efferent pseudobranchial artery within basicranial cartilage**
0 absent
1 present

**145. Position of basal/basipterygoid articulation** (Changed from '1' to '?' for *Debeerius*.)
0 same anteroposterior level as hypophysial opening
1 anterior to hypophysial opening
2 posterior to hypophysial opening

**146. Articulation between neurocanium and palatoquadrate posterodorsal to orbit (suprapterygoid articulation)**
0 absent
1 present

**147. Labyrinth cavity**
0 separated from the main neurocranial cavity by a cartilaginous or ossified capsular wall
1 skeletal capsular wall absent

**148. Basipterygoid process (basal articulation) with vertically oriented component**
0 absent

1 present

**149. Pituitary vein canal**
0 dorsal to level of basipterygoid process
1 flanked posteriorly by basipterygoid process

**150. External (horizontal) semicircular canal**
0 absent
1 present

**151. Sinus superior**
0 absent or indistinguishable from union of anterior and posterior canals with saccular chamber
1 present

**152. External (horizontal) semicircular canal** (Changed from '1' to '?' for *Moythomasia*.)
0 joins the vestibular region dorsal to posterior ampulla
1 joins level with posterior ampulla

**153. Horizontal semicircular canal in dorsal view**
0 medial to path of jugular vein
1 dorsal to jugular vein

**154. Lateral cranial canal**
0 absent
1 present

**155. Posterior dorsal fontanelle** (Changed from '?' to '0' for *Chondrenchelys*.)
0 absent
1 present

**156. Shape of posterior dorsal fontanelle** (Changed from '?' to '-' for *Chondrenchelys*.)
0 approximately as long as broad
1 much longer than wide, slot-shaped

**157. Synotic tectum** (Changed from '?' to '-' for *Chondrenchelys*.)
0 absent
1 present

**158. Dorsal ridge**
0 absent
1 present

**159. Shape of median dorsal ridge anterior to endolymphatic fossa**
0 developed as a squared-off ridge or otherwise ungrooved
1 bears a midline groove

**160. Endolymphatic ducts in neurocranium** (Changed code from '?' to '1' for *Chondrenchelys*.)
0 posteriodorsally angled tubes

1 tubes oriented vertically through median endolymphatic fossa

**161. Position of hyomandibula articulation on neurocranium**. Galeaspids are scored as state 1; the presumed hyoid arch is in a postorbital position based on the course of the facial nerve[14]. There is no evidence of an osteostracans-like topology for this articulation.

0 below or anterior to orbit, on ventrolateral angle of braincase
1 on otic capsule, posterior to orbit

**162. Position of hyomandibula articulation relative to structure of skeletal labyrinth**

0 anterior or lateral to skeletal labyrinth
1 at level of posterior semicircular canal

**163. Hyoid arch articulation on braincase**

0 single
1 double

**164. Branchial ridges.** Ordered character.

0 present
1 reduced to vagal process
2 absent (articulation made with bare cranial wall)

**165. Craniospinal process** (Changed from ??? to ?0? for Chondrenchelys.)

0 absent
1 present

**166. Ventral cranial fissure.** Changed from '-' to '0' for *Chondrenchelys*. *Minjinia* is scored as 'present', but this could be argued to be taphonomic.

0 absent
1 present

**167. Basicranial fenestra**

0 absent
1 present

**168. Metotic (otic-occipital) fissure.** *Janusiscus* is scored as 'present' based on ongoing work on high-resolution tomography scanning.

0 absent
1 present

**169. Vestibular fontanelle** (Changed from '1' to '?' for *Porolepis*; '?' to '0' for *Chondrenchelys*.)

0 absent
1 present

**170. Occipital arch wedged in between otic capsules** (Changed from '?' to '0' for *Psarolepis*.)

0 absent
1 present

**171. Spino-occipital nerve foramina**

0 two or more, aligned horizontally
1 one or two, dorsoventrally offset

**172. Ventral notch between parachordals**
0 present or entirely unfused
1 absent

**173. Parachordal shape**
0 forming a broad, flat surface as wide as the otic capsules
1 mediolaterally constricted relative to the otic capsules

**174. Stalk-shaped parachordal/occipital region**
0 absent
1 present

**175. Paired occipital facets**
0 absent
1 present

**176. Size of aperture to notochordal canal**
0 much smaller than foramen magnum
1 as large, or larger, than foramen magnum

**177. Canal for median dorsal aorta within basicranium** (Changed from '?' to '0' for *Chondrenchelys*.)
0 absent
1 present

**178. Hypotic lamina (and dorsally directed glossopharyngeal canal)** (Changed from '?' to '-' for *Chondrenchelys*.)
0 absent
1 present

**179. Macromeric dermal shoulder girdle**
0 present
1 absent

**180. Dermal shoulder girdle composition**
0 ventral and dorsal (scapular) components
1 ventral components only

**181. Shape of dorsal blade of dermal shoulder girdle (either cleithrum or anterolateral plate)**
0 spatulate
1 pointed

**182. Dermal shoulder girdle forming a complete ring around the trunk**
0 present
1 absent

**183. Pectoral fenestra completely encircled by dermal shoulder armour.** *Lunaspis* is re-coded as from absent to "?". We consider the interpretation of the highly flattened and lightly metamorphosed Hunsrückschiefer specimens to be too uncertain to code with confidence. The absence of this connection stems from Stensiö's (1963) reconstruction which has been reproduced by Denison [15] and Goutjet [13]. The encirclement of the pectoral fenestra by the demal thoracic armour was thought to be restricted to the antiarchs and arthrodires. However, since these publications Liu [16] described the pectoral armour of *Eurycaraspis*, a putative petalichthyid from China. Liu described a posterior lateral plate which links the PDL to the PVL plate in that taxon. The connection is not obvious in dorsal view. A possible corresponding plate appears in *Lunaspis* in Gross (1961, fig. 4).

0 present
1 absent

**184. Median dorsal plate**

0 absent
1 present

**185. Posterior dorsolateral (PDL) plate or equivalent**

0 absent
1 present

**186. Pronounced internal median keel on dorsal shoulder girdle** (i.e., crista of median dorsal plate)

0 absent
1 present

**187. Crista internalis of dermal shoulder girdle**

0 absent
1 present

**188. Scapular infundibulum**

0 absent
1 present

**189. Scapular process of shoulder endoskeleton**

0 absent
1 present

**190. Ventral margin of separate scapular ossification**

0 horizontal
1 deeply angled

**191. Cross sectional shape of scapular process**

0 flattened or strongly ovate
1 subcircular

**192. Flange on trailing edge of scapulocoracoid**

0 absent
1 present

**193. Scapular process with posterodorsal angle**
0 absent
1 present

**194. Endoskeletal postbranchial lamina on scapular process**
0 present
1 absent

**195. Mineralisation of internal surface of scapular blade**
0 mineralised all around
1 unmineralised on internal face forming a hemicylindrical cross-section

**196. Coracoid process**
0 absent
1 present

**197. Procoracoid mineralisation**
0 absent
1 present

**198. Fin base articulation on scapulocoracoid**
0 deeper than wide (stenobasal)
1 wider than deep (eurybasal)

**199. Pectoral fin articulation**
0 monobasal
1 polybasal

**200. Number of basals in polybasal pectoral fins**
0 three or more
1 two

**201. Branching radials in paired fins**
0 absent
1 present

**202. Number of mesomeres in metapterygial axis**
0 five or fewer
1 seven or more

**203. Biserial pectoral fin endoskeleton**
0 absent
1 present

**204. Perforate propterygium** (Changed from '-' to '0' for *Chondrenchelys*.)
0 absent
1 present

**205. Filamentous extension of pectoral fin from axillary region**
0 absent

1 present

**206. Pelvic fins** (Changed from '?' to '0' for Galeaspida. )
0 absent
1 present

**207. Pelvic claspers**
0 absent
1 present

**208. Dermal pelvic clasper ossifications**
0 absent
1 present

**209. Pectoral fins covered in macromeric dermal armour**
0 absent
1 present

**210. Pectoral fin base has large, hemispherical dermal component** (Changed from ??? to ?0? for *Styloichthys*.)
0 absent
1 present

**211. Dorsal fin spines.** Osteostraci has been changed to 0/1. These are absent in *Ateleaspis* [17] but a spine or substantial leading-edge scute is found in some genera. Placoderm-grade taxa lacking complete body preservation are uniformly scored as '?'. Dupret [18] suggests spines were present in *Kujdanowiaspis*. While these seem reasonable, there are no spines figured either articulated or isolated and must be scored as unknown based on this evidence.
0 absent
1 present

**212. Anal fin spine**
0 absent
1 present

**213. Paired fin spines**
0 absent
1 present

**214. Median fin spine insertion**
0 shallow, not greatly deeper than dermal bones/scales
1 deep

**215. Intermediate fin spines**
0 absent
1 present

**216. Fin spine cross-section**
0 Round or horseshoe shaped
1 Flat-sided, with rectangular profile

**217. Intermediate spines when present**
0 one pair
1 multiple pairs

**218. Prepectoral fin spines**
0 absent
1 present

**219. Fin spines with ridges**
0 absent
1 present

**220. Fin spines with nodes**
0 absent
1 present

**221. Fin spines with rows of large retrorse denticles**
0 absent
1 present

**222. Expanded spine rib on leading edge of spine**
0 absent
1 present

**223. Spine ridges**
0 converging at the distal apex of the spine
1 converging on leading edge of spine

**224. Synarcual**
0 absent
1 present

**225. Series of thoracic supraneurals**
0 absent
1 present

**226. Number of dorsal fins, if present**
0 one
1 two

**227. Posterior dorsal fin shape**
0 base approximately as broad as tall, not broader than all of other median fins
1 base much longer than the height of the fin, substantially longer than any of the other dorsal fins

**228. Basal plate in dorsal fin**
0 absent
1 present

**229. Branching radial structure articulating with dorsal fin basal plate**
0 absent
1 present

**230. Anal fin.** *Lunaspis* has been scored as missing because this part of the specimens figured by [19] are not complete.
0 absent
1 present

**231. Basal plate in anal fin**
0 absent
1 present

**232. Caudal radials**
0 extend beyond level of body wall and deep into hypochordal lobe
1 radials restricted to axial lobe

**233. Supraneurals in axial lobe of caudal fin**
0 absent
1 present

**234. Epichordal lepidotrichia in caudal fin**
0 absent
1 present

**235. Enamel and pore canals** (Zhu and Schultze 2001; Zhu et al. 2001; Zhu and Yu 2002; Zhu et al. 2006; Friedman 2007a.Taxa that lack pore openings are coded as inapplicable for this character). This character is contingent on both presence of enamel and presence of pore-canal systems in the dermal bones. We have made the character contingent on a pore-canal network, but we have verified that the scoring is consistent with the distribution of enamel.
0 enamel absent from inner surface of pores
1 enamel lines portions of pore canal

**236. Canal-bearing bone of skull roof extends far past posterior margin of parietals.**
0 no
1 yes

**237. Pineal eminence (in taxa lacking pineal foramen)**
0 absent
1 present

**238. Position of anterior pitline**
0 on postparietal
1 on parietal

**239. Opening in dermal skull roof for spiracular bounded by bones carrying otic canal**
0 absent
1 present

**240. Median skull roof bone between postparietals**

0 absent
1 present

**241. Westoll lines**
0 absent
1 present

**242. Preoperculosubmandibular**
0 absent
1 present

**243. Hyomandibula**
0 imperforate
1 perforate

**244. Urohyal shape**
0 absent
1 vertical plate

**245. Maxilla (in taxa with marginal jaw bones)**
0 present
1 absent

**246. Length of dentary**
0 constitutes a majority of jaw length
1 half the length of jaw or less

**247. Labial pit**
0 absent
1 present

**248. Prearticular symphysis**
0 absent
1 present

**249. Mandibular sensory canal**
0 extends through infradentaries
1 extends through infradentaries and dentary

**250. Extensive flange composed of prearticular and Meckelian bone that extends beyond ventral edge of outer dermal series**
0 absent
1 present

**251. Posterior coronoid**
0 similar to anterior coronoids
1 forms expanded coronoid process

**252. Retroarticular process**
0 absent

1 present

**253. Inturned medial process of premaxilla**
0 absent
1 present

**254. Anteriorly directed adductor fossae between neurocranium and skull roof**
0 absent
1 present

**255. Vomerine fangs**
0 absent
1 present

**256. Number of dermopalatines**
0 multiple
1 one

**257. Entopterygoids**
0 separated
1 contact along midline

**258. Rostral tubuli**
0 absent
1 present

**259. Position of anterior nostril**
0 facial
1 at oral margin

**260. Posterior nostril.** Ordered character.
0 facial
1 at margin of oral cavity
2 palatal

**261. Three large pores (in addition to nostrils) associated with each side of ethmoid**
0 absent
1 present

**262. Ventral face of nasal capsule in taxa with mineralized ethmoid** Ordered character.
0 complete
1 fenestra ventrolateralis
2 entire floor unmineralized

**263. Size of profundus canal in postnasal wall**
0 small
1 large

**264. Paired pineal and parapineal tracts**
0 absent

1 present

**265. Posterior of parasphenoid**
0 restricted to ethmosphenoid region
1 extends to otic region

**266. Endoskeletal spiracular canal.** Ordered character.
0 open
1 spiracular bar
2 complete enclosure in canal

**267. Barbed lepidotrichial segments.** All codings in this corrected to match parent character (presence of lepidotrichia).
0 absent
1 present

**268. Relative position of jugular groove/canal and hyomandibular articulation.** Ordered character.
0 hyomandibula dorsal
1 hmd straddles
2 hmd ventral

**269. Optic lobes**
0 narrower than cerebellum
1 same width or wider than cerebellum

**270. Hypophyseal chamber**
0 projects posteroventrally
1 projects ventrally or anteroventrally

**271. Crus commune of anterior and posterior semicircular canals**
0 dorsal to braincase endocavity roof
1 ventral to braincase endocavity roof

**272. Horizontal semicircular canal**
0 obiquely oriented
1 horizontally oriented

**273. Supraotic cavity**
0 absent
1 present

**274. Pelvic girdle with substantial dermal component**
0 present
1 absent

**275. Pelvic fin spine**
0 absent
1 present

**276. Pelvic fin**

0 monobasal
1 polybasal

**277. Postparietals/centrals.** Rhenanids are scored as '1'. This is based on Westoll (1967) and *Jagorina* specimen at MB.f.510.1.

0 absent
1 present

**278. Condition of postparietals/centrals.** *Eurycaraspis* changed to inapplicable. Rhenanids are scored as '1'. This is based on Westoll (1967) and *Jagorina* specimen at MB.f.510.1.

0 do not meet in midline
1 meet in midline
2 single midline bone

**279. Parietals**

0 absent
1 present

**280. Condition of parietals**

0 do not meet in midline
1 meet in midline

**281. Endoskeletal lamina (postnasal wall) separating posterior nostril and orbit**

0 absent
1 present

**282. Pituitary vein canal**

0 discontinuous, enters the cranial cavity
1 discontinuous, enters hypophysial recess
2 continuous transverse vein

**283. Sutures between dermal bones.** New character.

0 absent
1 present

**284. Interolateral/clavicular margin.** New character.

0 Angled anterolaterally
1 Mediolaterally straight

***Deleted characters***

We have deleted c.99 (Palatoquadrate relationship to dermal cheek bones: (0) articulation narrow and restricted; (1) broad articulation) from the matrix of Clement et al. (2018). It originated in the matrix of Giles et al. [20], however we consider it too vaguely stated to be of much value as the terms 'narrow' and 'broad' are not defined. It is beyond the data available to us to re-define this character based on a clear, standardised metric.

### Character state transformations

```
Tree length = 831
Consistency index (CI) = 0.3682
Homoplasy index (HI) = 0.6318
CI excluding uninformative characters = 0.3667
HI excluding uninformative characters = 0.6333
Retention index (RI) = 0.7871
Rescaled consistency index (RC) = 0.2898
```

```
/--------------------------------------------------------------------------------- Galeaspida
|
+--------------------------------------------------------------------------------- Osteostraci
|
|                                                                             /--- Gemuendina
|                                                                     /-----185
|                                                                     |       \--- Jagorina
188/----------------------------------------------------------------186
|  |                                                                  |  /-------- Yunnanolepis
|  |                                                                  |  |
|  |                                                                  \184 /------ Parayunnanolepis
|  |                                                                     | |
|  |                                                                     183  /--- Bothriolepis
|  |                                                                       \182
\187                                                                          \--- Pterichthyodes
   |
   |  /--------------------------------------------------------------------------- Brindabellaspis
   |  |
   |  |                                                                       /--- Lunaspis
   |  | /-------------------------------------------------------------------179
   \181 |                                                                     \--- Macropetalichthys
      | |
      | |                                                                  /------ Rhamphodopsis
      180                                                                  |
        |  /-------------------------------------------------------------177  /--- Austroptyctodus
        |  |                                                               \176
        |  |                                                                  \--- Campbellodus
        |  |
        \178  /------------------------------------------------------------------- Eurycaraspis
           |  |
           |  |  /---------------------------------------------------------------- Romundina
           |  |  |
           \175  |                                                 /-------------- Kujdanowiaspis
              |  |                                                 |
              |  |  /--------------------------------------------172  /----------- Dicksonosteus
              |  |  |                                              |  |
              \174  |                                              \171  /-------- Cowralepis
                 |  |                                                 |  |
                 |  |                                                 \170 /------ Buchanosteus
                 |  |                                                    | |
                 |  |                                                    169  /--- Coccosteus
                 \173                                                      \168
                    |                                                         \--- Incisoscutum
                    |
                    | /----------------------------------------------------------- Minjinia
                    | |
                    | |  /-------------------------------------------------------- Entelognathus
                    | |  |
                    167  |  /----------------------------------------------------- Ramirosuarezia
                      |  |  |
                      |  |  |  /-------------------------------------------------- Janusiscus
                      |  |  |  |
                      |  |  |  |               /---------------------------------- Dialipina
                      \166  |  |               |
                         |  |  |               |  /------------------------------- "Ligulalepis"
                         |  |  |               |  |
                         |  |  |  /----------162  |                      /-------- Guiyu
                         |  |  |  |            |  |                      |
                         |  |  |  |            |  |  /-----------------159 /------ Sparalepis
                         |  |  |  |            |  |  |                   | |
                         \165  |  |            \161  |                   158  /--- Psarolepis
                            |  |  |               |  |                     \157
                            |  |  |               |  |                        \--- Achoania
                            |  |  |               |  |
                            |  |  |               |  |       /-------------------- Meemannia
                            |  |  |               |  |       |
                            |  |  |               |  |  /--140  /----------------- Cheirolepis
                            |  |  |               \160  |    |  |
                            |  |  |                  |  |    \139  /-------------- Howqualepis
                            |  |  |                  |  |       |  |
                            |  |  |                  |  |       |  |          /--- Mimipiscis
                            \164  |                  |  |       \138  /-----136
                               |  |                  |  |          |  |       \--- Raynerius
                               |  |                  |  |          \137
                               |  |                  |  |             |  /-------- Moythomasia
                               |  |                  |  |             |  |
                               |  |                  |  |             \135 /------ Kentuckia
                               |  |                  \156                | |
                               |  |                     |                134  /--- Lawrenciella
                               |  |                     |                  \133
                               |  |                     |                     \--- Kansasiella
                               |  |                     |
                               |  |                     |  /---------------------- Ptyctolepis
                               |  |                     |  |
                               |  |                     |  |                  /--- Onychodus
                               |  |                     |  |             /--150
```

```
| |             | |         |  \--- Qingmenodus
| |             \155 /-----153
| |               | |       | /------ Miguashaia
| |               | |       | |
| |               | |      152  /--- Diplocercides
| |               | |        \151
| |               154           \--- Gavinia
| |                 |
\163                | /----------------- Styloichthys
  |                 | |
  |                 | |             /--- Eusthenopteron
  |                 \149 /--------141
  |                   |  |          \--- Gogonasus
  |                   |  |
  |                   |  |       /------ Powichthys
  |                   \148       |
  |                     |  /--143   /--- Glyptolepis
  |                     |  |   \142
  |                     |  |        \--- Porolepis
  |                     \147
  |                       | /-------- Youngolepis
  |                       | |
  |                       \146 /------ Diabolepis
  |                         |  |
  |                        145  /--- Dipterus
  |                          \144
  |                             \--- Uranolophus
  |
  |                             /--- Culmacanthus
  |               /-----------106
  |               |             \--- Tetanopsyrus
  |               |
  |               |             /--- Diplacanthus
  | /-----------107 /---------104
  | |             | |           \--- Gladiobranchus
  | |             | |
  | |           105             /--- Ischnacanthus
  | |             | /---------102
  | |             | |           \--- Poracanthodes
  | |             | |
  | |           \103 /-------------- Cassidiceps
  | |               | |
  | |             \101 /------------ Euthacanthus
  | |                 | |
  | |                 | |       /--- Mesacanthus
  132               \100 /----98
    |                   |  |    \--- Promesacanthus
    |                   \-99
    |                      | /------ Cheiracanthus
    |                      | |
    |                     \97   /--- Acanthodes
    |                        \-96
    |                           \--- Homalacanthus
    |
    | /------------------------------ Latviacanthus
    | |
    | |                         /--- Brochoadmones
    | | /-----------------------129
  \131  |                       \--- Kathemacanthus
      | |
      | |                 /--------- V waynensis
      | |                 |
    \130 /---------------127 /------ Ptomacanthus
       | |                |  |
       | |              \126 /------ Climatius
       | |                 | |
       | |                125   /--- Brachyacanthus
     \128                    \124
         |                      \--- Parexus
         |
         | /------------------------ Lupopsyrus
         | |
        123 /----------------------- Gyracanthides
          | |
        \122 /---------------------- Obtusacanthus
            | |
          \121 /-------------------- Pucapampella
              | |
            \120 /------------------ Doliodus
                | |
                | | /--------------- Cladodoides
              \119 |
                 | |            /--- Orthacanthus
                 | | /----------116
                118  |          \--- Tamiobatis
                   | |
                   | |       /------ Cobelodus
                 \117        |
                     | /----109 /--- Akmonistion
                     | |     \108
                     | |        \--- Cladoselache
                   \115
                       | /---------- Tristychius
                       | |
                       | |      /--- Hamiltonichthys
                     \114 /---112
                         | |    \--- Onychoselache
                       \113
                           | /------ Helodus
                           | |
```

```
111  /--- Chondrenchelys
  \110
     \--- Debeerius
```

Apomorphy lists:

| Branch | Character | Steps | CI | Change |
|---|---|---|---|---|
| node_188 --> Galeaspida | 3 | 1 | 0.167 | 0 ==> 1 |
| | 9 | 1 | 0.333 | 1 ==> 0 |
| | 17 | 1 | 0.400 | 0 ==> 1 |
| | 23 | 1 | 0.125 | 0 --> 1 |
| | 26 | 1 | 0.400 | 0 ==> 1 |
| | 27 | 1 | 0.167 | 1 ==> 0 |
| | 31 | 1 | 0.091 | 0 --> 1 |
| | 52 | 1 | 0.200 | 1 ==> 0 |
| | 151 | 1 | 0.500 | 0 --> 1 |
| | 161 | 1 | 0.333 | 0 ==> 1 |
| node_188 --> Osteostraci | 8 | 1 | 0.167 | 0 ==> 1 |
| | 21 | 1 | 0.143 | 0 --> 1 |
| | 24 | 1 | 0.091 | 0 --> 1 |
| | 177 | 1 | 0.200 | 0 --> 1 |
| | 230 | 1 | 0.200 | 0 ==> 1 |
| node_188 --> node_187 | 10 | 1 | 0.333 | 0 --> 1 |
| | 33 | 1 | 0.500 | 1 ==> 0 |
| | 77 | 1 | 0.167 | 0 ==> 1 |
| | 120 | 1 | 0.500 | 0 ==> 1 |
| | 150 | 1 | 1.000 | 0 ==> 1 |
| | 164 | 1 | 0.400 | 0 ==> 1 |
| | 175 | 1 | 0.500 | 0 ==> 1 |
| | 194 | 1 | 0.500 | 0 ==> 1 |
| | 201 | 1 | 1.000 | 0 --> 1 |
| | 206 | 1 | 1.000 | 0 ==> 1 |
| | 224 | 1 | 0.250 | 0 --> 1 |
| | 282 | 1 | 0.500 | 0 ==> 2 |
| | 283 | 1 | 0.500 | 0 ==> 1 |
| node_187 --> node_181 | 31 | 1 | 0.091 | 0 --> 1 |
| | 35 ( tessera morph) | 1 | 0.200 | 0 --> 1 |
| | 42 | 1 | 0.111 | 0 ==> 1 |
| | 60 | 1 | 0.750 | 2 --> 3 |
| | 199 | 1 | 0.500 | 0 --> 1 |
| | 207 | 1 | 0.333 | 0 --> 1 |
| | 227 | 1 | 0.250 | 0 --> 1 |
| | 279 | 1 | 0.500 | 0 --> 1 |
| node_181 --> node_180 | 21 | 1 | 0.143 | 0 --> 1 |
| | 24 | 1 | 0.091 | 0 --> 1 |
| | 30 | 1 | 0.333 | 0 ==> 1 |
| | 41 | 1 | 1.000 | 0 ==> 1 |
| | 43 | 1 | 0.250 | 0 ==> 1 |
| | 45 | 1 | 0.250 | 0 ==> 1 |
| | 114 | 1 | 0.333 | 0 ==> 1 |
| | 117 | 1 | 0.286 | 0 --> 1 |
| | 125 | 1 | 0.667 | 0 ==> 1 |
| node_180 --> node_178 | 127 | 1 | 0.500 | 0 --> 1 |
| | 142 | 1 | 0.200 | 1 ==> 0 |
| | 211 | 1 | 0.167 | 0 ==> 1 |
| node_178 --> node_175 | 31 | 1 | 0.091 | 1 --> 0 |
| | 91 | 1 | 0.333 | 0 --> 1 |
| | 129 (orbit dorsal or dorsolat facing) | 1 | 0.500 | 0 ==> 1 |
| | 161 | 1 | 0.333 | 0 --> 1 |
| | 230 | 1 | 0.200 | 0 --> 1 |
| | 274 | 1 | 0.500 | 0 --> 1 |
| node_175 --> node_174 | 30 | 1 | 0.333 | 1 ==> 0 |
| | 45 | 1 | 0.250 | 1 --> 0 |
| | 60 | 1 | 0.750 | 3 ==> 0 |
| | 284 | 1 | 0.200 | 1 --> 0 |
| node_174 --> node_173 | 95 | 1 | 0.500 | 0 ==> 1 |
| | 96 | 1 | 1.000 | 0 ==> 1 |
| | 136 | 1 | 0.500 | 0 ==> 1 |
| | 198 | 1 | 0.200 | 0 ==> 1 |
| | 278 | 1 | 0.400 | 0 ==> 1 |
| | 280 | 1 | 0.250 | 0 ==> 1 |
| node_173 --> node_167 | 10 | 1 | 0.333 | 1 --> 2 |
| | 21 | 1 | 0.143 | 1 --> 0 |
| | 26 | 1 | 0.400 | 0 --> 1 |
| | 39 (endolymph in sr) | 1 | 0.333 | 0 --> 1 |
| | 62 | 1 | 0.200 | 0 --> 1 |
| | 68 | 1 | 0.500 | 0 --> 1 |
| | 73 | 1 | 0.500 | 0 --> 1 |
| | 120 | 1 | 0.500 | 1 --> 0 |
| | 121 | 1 | 1.000 | 0 --> 1 |
| | 127 | 1 | 0.500 | 1 --> 0 |
| | 137 | 1 | 0.333 | 0 --> 1 |
| | 141 | 1 | 0.500 | 0 --> 1 |

| | | | | |
|---|---|---|---|---|
| | 147 (labyrinth cavity) | 1 | 1.000 | 0 ==> 1 |
| | 153 | 1 | 0.333 | 0 --> 1 |
| | 164 | 1 | 0.400 | 1 --> 2 |
| | 175 | 1 | 0.500 | 1 --> 0 |
| | 196 | 1 | 0.333 | 1 --> 0 |
| | 207 | 1 | 0.333 | 1 --> 0 |
| | 220 | 1 | 0.125 | 1 --> 0 |
| | 224 | 1 | 0.250 | 1 --> 0 |
| | 227 | 1 | 0.250 | 1 --> 0 |
| | 272 | 1 | 0.250 | 0 --> 1 |
| | 281 | 1 | 0.500 | 0 --> 1 |
| | 282 | 1 | 0.500 | 2 --> 1 |
| node_167 --> node_166 | 29 | 1 | 0.214 | 3 ==> 1 |
| | 151 | 1 | 0.500 | 0 --> 1 |
| | 176 | 1 | 1.000 | 0 --> 1 |
| node_166 --> node_165 | 53 | 1 | 0.333 | 1 ==> 0 |
| | 78 | 1 | 0.167 | 0 ==> 1 |
| | 98 | 1 | 0.333 | 0 --> 1 |
| | 99 | 1 | 0.500 | 0 --> 1 |
| | 134 | 1 | 0.250 | 0 --> 1 |
| | 165 | 1 | 0.500 | 1 ==> 0 |
| | 173 | 1 | 0.250 | 0 ==> 1 |
| | 181 | 1 | 0.250 | 0 --> 1 |
| | 182 | 1 | 1.000 | 0 --> 1 |
| | 183 (pectoral fenestra) | 1 | 0.250 | 0 --> 1 |
| | 185 | 1 | 0.333 | 1 --> 0 |
| | 219 | 1 | 0.200 | 0 --> 1 |
| node_165 --> node_164 | 132 | 1 | 0.333 | 0 --> 1 |
| | 168 | 1 | 0.250 | 0 ==> 1 |
| node_164 --> node_163 | 123 | 1 | 0.200 | 0 --> 1 |
| | 155 | 1 | 0.250 | 0 ==> 1 |
| | 166 (ventral cranial fissure) | 1 | 0.250 | 0 ==> 1 |
| node_163 --> node_132 | 27 | 1 | 0.167 | 1 ==> 0 |
| | 28 | 1 | 0.500 | 0 ==> 1 |
| | 29 | 1 | 0.214 | 1 --> 3 |
| | 33 | 1 | 0.500 | 0 ==> 1 |
| | 39 (endolymph in sr) | 1 | 0.333 | 1 --> 0 |
| | 52 | 1 | 0.200 | 1 ==> 0 |
| | 63 | 1 | 0.500 | 1 --> 0 |
| | 68 | 1 | 0.500 | 1 --> 0 |
| | 69 | 1 | 1.000 | 1 ==> 0 |
| | 72 | 1 | 0.500 | 0 ==> 1 |
| | 81 | 1 | 0.143 | 0 ==> 1 |
| | 91 | 1 | 0.333 | 1 ==> 0 |
| | 108 | 1 | 0.500 | 0 --> 1 |
| | 124 | 1 | 0.200 | 0 ==> 1 |
| | 125 | 1 | 0.667 | 1 ==> 2 |
| | 138 | 1 | 1.000 | 0 --> 1 |
| | 146 | 1 | 0.200 | 0 ==> 1 |
| | 153 | 1 | 0.333 | 1 ==> 0 |
| | 162 | 1 | 1.000 | 0 ==> 1 |
| | 179 | 1 | 0.167 | 0 ==> 1 |
| | 180 | 1 | 0.500 | 0 ==> 1 |
| | 189 | 1 | 1.000 | 0 ==> 1 |
| | 197 | 1 | 0.143 | 0 ==> 1 |
| | 212 | 1 | 0.500 | 0 ==> 1 |
| | 214 | 1 | 0.167 | 0 ==> 1 |
| | 215 | 1 | 0.167 | 0 --> 1 |
| | 222 | 1 | 0.500 | 0 --> 1 |
| | 232 | 1 | 1.000 | 1 ==> 0 |
| | 268 (rel position of jugular/hmd) | 2 | 0.667 | 0 ==> 2 |
| | 275 | 1 | 0.200 | 0 ==> 1 |
| | 283 | 1 | 0.500 | 1 ==> 0 |
| node_132 --> node_107 | 12 | 1 | 0.500 | 0 --> 1 |
| | 16 | 1 | 0.333 | 0 ==> 1 |
| | 22 | 1 | 0.500 | 0 ==> 1 |
| | 23 | 1 | 0.125 | 0 ==> 1 |
| | 24 | 1 | 0.091 | 1 --> 0 |
| | 77 | 1 | 0.167 | 1 ==> 0 |
| | 106 | 1 | 0.500 | 0 --> 1 |
| | 132 | 1 | 0.333 | 1 --> 0 |
| | 143 | 1 | 0.143 | 0 --> 1 |
| node_107 --> node_105 | 61 | 1 | 0.429 | 0 --> 1 |
| | 190 | 1 | 0.333 | 0 ==> 1 |
| node_105 --> node_103 | 63 | 1 | 0.500 | 0 --> 1 |
| | 64 | 1 | 1.000 | 0 ==> 1 |
| | 65 | 1 | 0.500 | 0 ==> 1 |
| | 191 | 1 | 0.500 | 0 ==> 1 |
| | 195 | 1 | 0.250 | 0 --> 1 |
| | 216 | 1 | 0.500 | 0 ==> 1 |
| node_103 --> node_101 | 29 | 1 | 0.214 | 3 ==> 0 |
| | 52 | 1 | 0.200 | 0 ==> 1 |
| | 87 | 1 | 0.333 | 1 ==> 0 |
| | 214 | 1 | 0.167 | 1 ==> 0 |

| | | | | |
|---|---|---|---|---|
| node_100 --> node_99 | 11 | 1 | 0.333 | 0 ==> 1 |
| | 38 (extent of dermatocr) | 1 | 0.200 | 0 ==> 1 |
| | 61 | 1 | 0.429 | 1 --> 0 |
| | 195 | 1 | 0.250 | 1 --> 0 |
| | 217 | 1 | 0.500 | 1 --> 0 |
| | 226 | 1 | 0.143 | 1 ==> 0 |
| node_99 --> node_97 | 61 | 1 | 0.429 | 0 --> 2 |
| | 103 | 1 | 1.000 | 0 ==> 1 |
| | 214 | 1 | 0.167 | 0 ==> 1 |
| | 215 | 1 | 0.167 | 1 ==> 0 |
| node_97 --> node_96 | 64 | 1 | 1.000 | 1 ==> 2 |
| node_96 --> Acanthodes | 97 | 1 | 0.250 | 0 ==> 1 |
| | 101 | 1 | 0.250 | 0 ==> 1 |
| | 190 | 1 | 0.333 | 1 ==> 0 |
| node_99 --> node_98 | 35 ( tessera morph) | 1 | 0.200 | 1 ==> 0 |
| | 65 | 1 | 0.500 | 1 ==> 0 |
| node_98 --> Promesacanthus | 66 | 1 | 0.500 | 1 ==> 0 |
| | 218 | 1 | 0.167 | 0 ==> 1 |
| node_100 --> Euthacanthus | 10 | 1 | 0.333 | 2 ==> 0 |
| | 24 | 1 | 0.091 | 0 ==> 1 |
| | 179 | 1 | 0.167 | 1 ==> 0 |
| | 216 | 1 | 0.500 | 1 ==> 0 |
| | 218 | 1 | 0.167 | 0 ==> 1 |
| node_101 --> Cassidiceps | 35 ( tessera morph) | 1 | 0.200 | 1 ==> 0 |
| node_103 --> node_102 | 77 | 1 | 0.167 | 0 ==> 1 |
| | 101 | 1 | 0.250 | 0 --> 1 |
| | 215 | 1 | 0.167 | 1 --> 0 |
| | 275 | 1 | 0.200 | 1 --> 0 |
| node_102 --> Poracanthodes | 16 | 1 | 0.333 | 1 ==> 0 |
| node_105 --> node_104 | 11 | 1 | 0.333 | 0 ==> 1 |
| | 35 ( tessera morph) | 1 | 0.200 | 1 ==> 0 |
| | 61 | 1 | 0.429 | 1 --> 2 |
| | 78 | 1 | 0.167 | 1 --> 0 |
| | 180 | 1 | 0.500 | 1 --> 0 |
| | 192 | 1 | 1.000 | 0 ==> 1 |
| | 194 | 1 | 0.500 | 1 ==> 0 |
| | 217 | 1 | 0.500 | 1 --> 0 |
| node_104 --> Diplacanthus | 3 | 1 | 0.167 | 0 ==> 1 |
| | 10 | 1 | 0.333 | 2 ==> 0 |
| | 62 | 1 | 0.200 | 1 ==> 0 |
| | 179 | 1 | 0.167 | 1 ==> 0 |
| node_104 --> Gladiobranchus | 104 | 1 | 0.333 | 0 ==> 1 |
| | 196 | 1 | 0.333 | 0 ==> 1 |
| | 197 | 1 | 0.143 | 1 ==> 0 |
| | 218 | 1 | 0.167 | 0 ==> 1 |
| | 220 | 1 | 0.125 | 0 ==> 1 |
| node_107 --> node_106 | 29 | 1 | 0.214 | 3 --> 1 |
| | 62 | 1 | 0.200 | 1 ==> 0 |
| | 98 | 1 | 0.333 | 1 --> 0 |
| | 104 | 1 | 0.333 | 0 --> 1 |
| | 215 | 1 | 0.167 | 1 --> 0 |
| node_106 --> Culmacanthus | 53 | 1 | 0.333 | 0 ==> 1 |
| | 179 | 1 | 0.167 | 1 ==> 0 |
| | 197 | 1 | 0.143 | 1 ==> 0 |
| node_106 --> Tetanopsyrus | 28 | 1 | 0.500 | 1 ==> 0 |
| | 214 | 1 | 0.167 | 1 ==> 0 |
| | 220 | 1 | 0.125 | 0 ==> 1 |
| node_132 --> node_131 | 10 | 1 | 0.333 | 2 --> 0 |
| | 17 | 1 | 0.400 | 0 --> 1 |
| | 83 | 1 | 1.000 | 1 ==> 0 |
| | 84 | 1 | 0.667 | 0 ==> 1 |
| | 87 | 1 | 0.333 | 1 ==> 0 |
| | 123 | 1 | 0.200 | 1 --> 0 |
| | 141 | 1 | 0.500 | 1 --> 0 |
| | 173 | 1 | 0.250 | 1 --> 0 |
| | 178 | 1 | 1.000 | 0 --> 1 |
| | 202 | 1 | 0.250 | 0 --> 1 |
| node_131 --> node_130 | 66 | 1 | 0.500 | 1 --> 0 |
| | 196 | 1 | 0.333 | 0 ==> 1 |
| | 218 | 1 | 0.167 | 0 ==> 1 |
| | 220 | 1 | 0.125 | 0 --> 1 |
| | 222 | 1 | 0.500 | 1 --> 0 |
| node_130 --> node_128 | 106 | 1 | 0.500 | 0 ==> 1 |
| | 223 | 1 | 0.250 | 0 --> 1 |
| node_128 --> node_123 | 13 | 1 | 0.333 | 1 ==> 0 |
| | 25 | 1 | 0.500 | 0 ==> 1 |
| | 26 | 1 | 0.400 | 1 ==> 2 |
| | 79 | 1 | 0.250 | 0 --> 1 |
| | 142 | 1 | 0.200 | 0 --> 1 |
| | 144 | 1 | 0.333 | 0 --> 1 |
| | 193 | 1 | 0.500 | 0 --> 1 |
| node_123 --> node_122 | 10 | 1 | 0.333 | 0 --> 2 |
| | 62 | 1 | 0.200 | 1 ==> 0 |
| | 207 | 1 | 0.333 | 0 --> 1 |

| | | | | |
|---|---|---|---|---|
| | 215 | 1 | 0.167 | 1 ==> 0 |
| node_122 --> node_121 | 3 | 1 | 0.167 | 0 ==> 1 |
| node_121 --> node_120 | 1 | 1 | 1.000 | 0 ==> 1 |
| | 28 | 1 | 0.500 | 1 --> 2 |
| | 38 (extent of dermatocr) | 1 | 0.200 | 0 --> 1 |
| | 61 | 1 | 0.429 | 0 --> 3 |
| | 212 | 1 | 0.500 | 1 --> 0 |
| | 223 | 1 | 0.250 | 1 --> 0 |
| node_120 --> node_119 | 143 | 1 | 0.143 | 0 ==> 1 |
| | 145 | 1 | 0.667 | 0 --> 1 |
| | 156 | 1 | 0.333 | 0 ==> 1 |
| | 160 | 1 | 1.000 | 0 --> 1 |
| | 166 (ventral cranial fissure) | 1 | 0.250 | 1 ==> 0 |
| | 172 | 1 | 0.200 | 0 --> 1 |
| node_119 --> node_118 | 75 | 1 | 0.500 | 0 --> 1 |
| | 82 | 1 | 0.500 | 0 ==> 1 |
| | 133 | 1 | 0.333 | 0 --> 1 |
| | 213 | 1 | 0.200 | 1 --> 0 |
| | 218 | 1 | 0.167 | 1 --> 0 |
| | 275 | 1 | 0.200 | 1 --> 0 |
| node_118 --> node_117 | 157 | 1 | 0.500 | 0 ==> 1 |
| | 269 | 1 | 0.500 | 0 ==> 1 |
| node_117 --> node_115 | 17 | 1 | 0.400 | 1 --> 0 |
| | 133 | 1 | 0.333 | 1 --> 0 |
| | 139 | 1 | 0.500 | 0 --> 1 |
| | 220 | 1 | 0.125 | 1 ==> 0 |
| | 270 | 1 | 0.250 | 0 --> 1 |
| | 271 | 1 | 0.500 | 0 --> 1 |
| | 272 | 1 | 0.250 | 1 --> 0 |
| node_115 --> node_109 | 123 | 1 | 0.200 | 0 ==> 1 |
| | 177 | 1 | 0.200 | 0 ==> 1 |
| | 205 | 1 | 0.500 | 0 --> 1 |
| | 219 | 1 | 0.200 | 1 --> 0 |
| | 226 | 1 | 0.143 | 1 --> 0 |
| | 230 | 1 | 0.200 | 1 ==> 0 |
| | 233 | 1 | 0.250 | 0 ==> 1 |
| node_109 --> node_108 | 128 | 1 | 0.500 | 0 ==> 1 |
| | 156 | 1 | 0.333 | 1 --> 0 |
| node_108 --> Akmonistion | 3 | 1 | 0.167 | 1 ==> 0 |
| | 15 | 1 | 0.200 | 1 ==> 0 |
| | 36 ( cranial spines abs/pres) | 1 | 0.500 | 0 ==> 1 |
| | 124 | 1 | 0.200 | 1 ==> 0 |
| | 226 | 1 | 0.143 | 0 --> 1 |
| node_108 --> Cladoselache | 197 | 1 | 0.143 | 1 ==> 0 |
| | 202 | 1 | 0.250 | 1 ==> 0 |
| | 205 | 1 | 0.500 | 1 --> 0 |
| node_109 --> Cobelodus | 135 | 1 | 0.500 | 0 ==> 1 |
| | 211 | 1 | 0.167 | 1 ==> 0 |
| | 227 | 1 | 0.250 | 0 ==> 1 |
| | 228 | 1 | 0.200 | 1 ==> 0 |
| node_115 --> node_114 | 37 (cranial spines monocuspid/multicuspid) | 1 | 1.000 | 0 --> 1 |
| | 98 | 1 | 0.333 | 1 ==> 0 |
| | 142 | 1 | 0.200 | 1 --> 0 |
| | 146 | 1 | 0.200 | 1 ==> 0 |
| | 159 | 1 | 1.000 | 0 ==> 1 |
| | 168 | 1 | 0.250 | 1 ==> 0 |
| | 171 | 1 | 0.200 | 0 --> 1 |
| | 197 | 1 | 0.143 | 1 ==> 0 |
| | 221 | 1 | 0.200 | 0 ==> 1 |
| | 229 | 1 | 0.333 | 0 --> 1 |
| node_114 --> node_113 | 116 | 1 | 0.250 | 0 --> 1 |
| | 143 | 1 | 0.143 | 1 --> 0 |
| | 158 | 1 | 0.500 | 1 ==> 0 |
| | 202 | 1 | 0.250 | 1 ==> 0 |
| node_113 --> node_111 | 22 | 1 | 0.500 | 0 --> 1 |
| | 23 | 1 | 0.125 | 0 --> 1 |
| | 72 | 1 | 0.500 | 1 --> 0 |
| | 100 | 1 | 0.333 | 0 --> 1 |
| | 107 | 1 | 0.500 | 0 ==> 1 |
| | 155 | 1 | 0.250 | 1 --> 0 |
| | 156 | 1 | 0.333 | 1 --> 0 |
| | 200 | 1 | 0.500 | 0 ==> 1 |
| | 227 | 1 | 0.250 | 0 ==> 1 |
| | 229 | 1 | 0.333 | 1 --> 0 |
| | 230 | 1 | 0.200 | 1 ==> 0 |
| node_111 --> node_110 | 15 | 1 | 0.200 | 1 ==> 0 |
| | 108 | 1 | 0.500 | 1 ==> 0 |
| | 117 | 1 | 0.286 | 1 ==> 0 |
| | 203 | 1 | 0.333 | 0 ==> 1 |
| node_110 --> Chondrenchelys | 82 | 1 | 0.500 | 1 ==> 0 |
| | 198 | 1 | 0.200 | 1 ==> 0 |
| | 211 | 1 | 0.167 | 1 ==> 0 |
| | 226 | 1 | 0.143 | 1 ==> 0 |
| | 228 | 1 | 0.200 | 1 ==> 0 |

| | | | | |
|---|---|---|---|---|
| node_110 --> Debeerius | 100 | 1 | 0.333 | 1 --> 0 |
| | 155 | 1 | 0.250 | 0 --> 1 |
| node_111 --> Helodus | 79 | 1 | 0.250 | 1 ==> 0 |
| | 142 | 1 | 0.200 | 0 --> 1 |
| | 144 | 1 | 0.333 | 1 ==> 0 |
| | 177 | 1 | 0.200 | 0 ==> 1 |
| | 193 | 1 | 0.500 | 1 ==> 0 |
| | 219 | 1 | 0.200 | 1 ==> 0 |
| | 221 | 1 | 0.200 | 1 ==> 0 |
| | 224 | 1 | 0.250 | 0 ==> 1 |
| node_113 --> node_112 | 5 | 1 | 0.200 | 0 --> 1 |
| | 28 | 1 | 0.500 | 2 --> 1 |
| | 36 ( cranial spines abs/pres) | 1 | 0.500 | 0 ==> 1 |
| | 118 | 1 | 0.500 | 0 --> 1 |
| | 170 | 1 | 0.333 | 0 ==> 1 |
| node_112 --> Hamiltonichthys | 15 | 1 | 0.200 | 1 ==> 0 |
| | 38 (extent of dermatocr) | 1 | 0.200 | 1 ==> 0 |
| | 233 | 1 | 0.250 | 0 ==> 1 |
| node_114 --> Tristychius | 95 | 1 | 0.500 | 1 ==> 0 |
| | 252 | 1 | 0.500 | 0 ==> 1 |
| node_117 --> node_116 | 2 | 1 | 1.000 | 0 ==> 1 |
| | 12 | 1 | 0.500 | 0 --> 1 |
| | 25 | 1 | 0.500 | 1 --> 0 |
| | 79 | 1 | 0.250 | 1 --> 0 |
| | 101 | 1 | 0.250 | 0 --> 1 |
| | 118 | 1 | 0.500 | 0 --> 1 |
| | 124 | 1 | 0.200 | 1 ==> 0 |
| | 200 | 1 | 0.500 | 0 --> 1 |
| | 203 | 1 | 0.333 | 0 --> 1 |
| | 228 | 1 | 0.200 | 1 --> 0 |
| | 231 | 1 | 0.500 | 1 --> 0 |
| node_116 --> Orthacanthus | 130 | 1 | 0.333 | 1 ==> 0 |
| | 170 | 1 | 0.333 | 0 ==> 1 |
| node_116 --> Tamiobatis | 221 | 1 | 0.200 | 0 ==> 1 |
| node_118 --> Cladodoides | 170 | 1 | 0.333 | 0 ==> 1 |
| node_119 --> Doliodus | 128 | 1 | 0.500 | 0 ==> 1 |
| | 129 (orbit dorsal or dorsolat facing) | 1 | 0.500 | 1 ==> 0 |
| | 140 | 1 | 0.333 | 0 ==> 1 |
| | 215 | 1 | 0.167 | 0 ==> 1 |
| node_121 --> Obtusacanthus | 15 | 1 | 0.200 | 1 ==> 0 |
| | 78 | 1 | 0.167 | 1 ==> 0 |
| node_122 --> Gyracanthides | 179 | 1 | 0.167 | 1 ==> 0 |
| | 195 | 1 | 0.250 | 0 ==> 1 |
| | 218 | 1 | 0.167 | 1 ==> 0 |
| node_123 --> Lupopsyrus | 15 | 1 | 0.200 | 1 ==> 0 |
| | 77 | 1 | 0.167 | 1 ==> 0 |
| | 117 | 1 | 0.286 | 1 ==> {02} |
| node_128 --> node_127 | 35 ( tessera morph) | 1 | 0.200 | 1 ==> 0 |
| | 52 | 1 | 0.200 | 0 --> 1 |
| | 134 | 1 | 0.250 | 1 --> 0 |
| | 140 | 1 | 0.333 | 0 --> 1 |
| | 179 | 1 | 0.167 | 1 ==> 0 |
| | 214 | 1 | 0.167 | 1 ==> 0 |
| node_127 --> node_126 | 197 | 1 | 0.143 | 1 ==> 0 |
| | 223 | 1 | 0.250 | 1 --> 0 |
| node_126 --> node_125 | 58 | 1 | 1.000 | 0 ==> 1 |
| node_125 --> node_124 | 191 | 1 | 0.500 | 0 ==> 1 |
| | 223 | 1 | 0.250 | 0 --> 1 |
| node_124 --> Brachyacanthus | 27 | 1 | 0.167 | 0 ==> 1 |
| node_124 --> Parexus | 190 | 1 | 0.333 | 0 ==> 1 |
| | 214 | 1 | 0.167 | 0 ==> 1 |
| | 221 | 1 | 0.200 | 0 ==> 1 |
| node_125 --> Climatius | 13 | 1 | 0.333 | 1 ==> 0 |
| | 84 | 1 | 0.667 | 1 ==> 0 |
| node_126 --> Ptomacanthus | 16 | 1 | 0.333 | 0 ==> 1 |
| | 23 | 1 | 0.125 | 0 ==> 1 |
| node_130 --> node_129 | 3 | 1 | 0.167 | 0 ==> 1 |
| | 61 | 1 | 0.429 | 0 --> 1 |
| | 62 | 1 | 0.200 | 1 ==> 0 |
| | 197 | 1 | 0.143 | 1 --> 0 |
| node_129 --> Brochoadmones | 23 | 1 | 0.125 | 0 ==> 1 |
| | 29 | 1 | 0.214 | 3 ==> 0 |
| | 221 | 1 | 0.200 | 0 ==> 1 |
| node_129 --> Kathemacanthus | 77 | 1 | 0.167 | 1 ==> 0 |
| | 220 | 1 | 0.125 | 1 --> 0 |
| node_131 --> Latviacanthus | 61 | 1 | 0.429 | 0 ==> 3 |
| | 84 | 1 | 0.667 | 1 ==> 2 |
| | 195 | 1 | 0.250 | 0 ==> 1 |
| node_163 --> node_162 | 4 | 1 | 0.333 | 0 --> 1 |
| | 5 | 1 | 0.200 | 0 ==> 1 |
| | 18 | 1 | 0.200 | 0 ==> 1 |
| | 20 | 1 | 0.200 | 0 --> 1 |
| | 21 | 1 | 0.143 | 0 --> 1 |
| | 67 | 1 | 0.250 | 0 --> 1 |

| | | | | |
|---|---|---|---|---|
| | 74 | 1 | 1.000 | 0 --> 1 |
| | 75 | 1 | 0.500 | 0 --> 1 |
| | 85 (teeth ankyolosed) | 1 | 0.500 | 1 ==> 0 |
| | 88 | 1 | 0.500 | 0 ==> 1 |
| | 101 | 1 | 0.250 | 0 --> 1 |
| | 109 | 1 | 0.500 | 0 --> 1 |
| | 126 | 1 | 0.333 | 0 --> 1 |
| | 139 | 1 | 0.500 | 0 --> 1 |
| | 171 | 1 | 0.200 | 0 --> 1 |
| | 172 | 1 | 0.200 | 0 --> 1 |
| | 213 | 1 | 0.200 | 1 --> 0 |
| | 225 | 1 | 0.500 | 0 --> 1 |
| node_162 --> node_161 | 6 | 1 | 0.500 | 0 --> 1 |
| | 14 | 1 | 1.000 | 0 --> 1 |
| | 30 | 1 | 0.333 | 0 ==> 1 |
| | 42 | 1 | 0.111 | 1 ==> 0 |
| | 53 | 1 | 0.333 | 0 --> 1 |
| | 90 | 1 | 0.500 | 0 --> 1 |
| node_161 --> node_160 | 8 | 1 | 0.167 | 0 ==> 1 |
| | 46 | 1 | 0.250 | 0 --> 1 |
| | 122 | 1 | 0.333 | 0 --> 1 |
| node_160 --> node_156 | 79 | 1 | 0.250 | 0 ==> 1 |
| | 110 | 1 | 0.500 | 0 ==> 1 |
| | 116 | 1 | 0.250 | 0 ==> 1 |
| | 130 | 1 | 0.333 | 1 ==> 0 |
| | 184 | 1 | 1.000 | 1 ==> 0 |
| | 211 | 1 | 0.167 | 1 ==> 0 |
| node_156 --> node_140 | 19 | 1 | 0.500 | 0 --> 1 |
| | 31 | 1 | 0.091 | 0 ==> 1 |
| | 32 | 1 | 1.000 | 0 --> 1 |
| | 46 | 1 | 0.250 | 1 --> 0 |
| | 71 | 1 | 0.500 | 0 --> 1 |
| | 93 | 1 | 1.000 | 0 --> 1 |
| | 102 | 1 | 1.000 | 0 --> 1 |
| | 122 | 1 | 0.333 | 1 --> 0 |
| | 152 | 1 | 0.333 | 0 --> 1 |
| | 154 | 1 | 1.000 | 0 ==> 1 |
| | 158 | 1 | 0.500 | 1 --> 0 |
| | 169 | 1 | 0.250 | 0 --> 1 |
| | 226 | 1 | 0.143 | 1 --> 0 |
| | 228 | 1 | 0.200 | 1 --> 0 |
| | 231 | 1 | 0.500 | 1 --> 0 |
| | 236 | 1 | 1.000 | 0 ==> 1 |
| | 238 | 1 | 0.500 | 1 --> 0 |
| | 239 | 1 | 0.500 | 0 --> 1 |
| | 266 | 1 | 0.333 | 0 --> 1 |
| node_140 --> node_139 | 7 | 1 | 1.000 | 1 ==> 0 |
| | 165 | 1 | 0.500 | 0 --> 1 |
| node_139 --> Cheirolepis | 21 | 1 | 0.143 | 1 ==> 0 |
| | 23 | 1 | 0.125 | 0 ==> 1 |
| | 24 | 1 | 0.091 | 1 ==> 0 |
| node_139 --> node_138 | 8 | 1 | 0.167 | 1 --> 0 |
| | 34 | 1 | 0.200 | 0 ==> 1 |
| | 80 | 1 | 1.000 | 0 --> 1 |
| | 111 | 1 | 0.500 | 0 ==> 1 |
| | 115 | 1 | 1.000 | 0 ==> 1 |
| | 135 | 1 | 0.500 | 0 --> 1 |
| | 204 | 1 | 1.000 | 0 ==> 1 |
| | 234 | 1 | 0.500 | 1 ==> 0 |
| node_138 --> Howqualepis | 48 | 1 | 0.286 | 0 ==> 1 |
| | 81 | 1 | 0.143 | 0 ==> 1 |
| | 92 | 1 | 0.333 | 0 ==> 1 |
| node_138 --> node_137 | 31 | 1 | 0.091 | 1 --> 0 |
| | 177 | 1 | 0.200 | 0 ==> 1 |
| | 243 | 1 | 0.333 | 0 ==> 1 |
| node_137 --> node_135 | 112 | 1 | 0.250 | 0 ==> 1 |
| | 269 | 1 | 0.500 | 0 --> 1 |
| | 271 | 1 | 0.500 | 0 --> 1 |
| node_135 --> node_134 | 111 | 1 | 0.500 | 1 --> 0 |
| | 266 | 1 | 0.333 | 1 ==> 2 |
| node_134 --> Kentuckia | 31 | 1 | 0.091 | 0 --> 1 |
| node_134 --> node_133 | 42 | 1 | 0.111 | 0 --> 1 |
| | 49 | 1 | 0.500 | 0 --> 1 |
| | 171 | 1 | 0.200 | 1 ==> 0 |
| node_133 --> Lawrenciella | 133 | 1 | 0.333 | 0 ==> 1 |
| node_137 --> node_136 | 172 | 1 | 0.200 | 1 --> 0 |
| | 266 | 1 | 0.333 | 1 --> 0 |
| node_136 --> Mimipiscis | 142 | 1 | 0.200 | 0 ==> 1 |
| node_136 --> Raynerius | 17 | 1 | 0.400 | 0 ==> 2 |
| | 24 | 1 | 0.091 | 1 ==> 0 |
| | 71 | 1 | 0.500 | 1 ==> 0 |
| | 73 | 1 | 0.500 | 1 ==> 0 |
| | 114 | 1 | 0.333 | 1 ==> 0 |
| node_140 --> Meemannia | 29 | 1 | 0.214 | 1 ==> 0 |

| | | | | |
|---|---|---|---|---|
| node_156 --> node_155 | 17 | 1 | 0.400 | 0 --> 2 |
| | 20 | 1 | 0.200 | 1 --> 0 |
| | 27 | 1 | 0.167 | 1 --> 0 |
| | 34 | 1 | 0.200 | 0 --> 1 |
| | 54 | 1 | 0.333 | 0 --> 1 |
| | 59 | 1 | 1.000 | 0 --> 1 |
| | 76 | 1 | 1.000 | 0 --> 1 |
| | 89 | 1 | 0.333 | 0 --> 1 |
| | 126 | 1 | 0.333 | 1 ==> 0 |
| | 146 | 1 | 0.200 | 0 --> 1 |
| | 163 | 1 | 0.500 | 0 ==> 1 |
| | 199 | 1 | 0.500 | 1 --> 0 |
| | 243 | 1 | 0.333 | 0 --> 1 |
| | 249 | 1 | 1.000 | 1 --> 0 |
| | 258 | 1 | 0.333 | 0 --> 1 |
| | 272 | 1 | 0.250 | 1 --> 0 |
| | 276 | 1 | 1.000 | 1 --> 0 |
| node_155 --> node_154 | 29 | 1 | 0.214 | 1 --> 0 |
| | 167 | 1 | 0.500 | 0 ==> 1 |
| | 268 (rel position of jugular/hmd) | 1 | 0.667 | 0 ==> 1 |
| node_154 --> node_149 | 86 | 1 | 0.500 | 0 --> 1 |
| | 90 | 1 | 0.500 | 1 ==> 0 |
| | 94 | 1 | 0.500 | 0 --> 1 |
| | 105 | 1 | 1.000 | 0 ==> 1 |
| | 237 | 1 | 0.200 | 0 ==> 1 |
| | 255 | 1 | 0.500 | 0 --> 1 |
| | 262 | 1 | 0.500 | 0 ==> 1 |
| | 272 | 1 | 0.250 | 0 --> 1 |
| | 273 | 1 | 1.000 | 0 --> 1 |
| node_149 --> node_148 | 6 | 1 | 0.500 | 1 ==> 0 |
| | 131 | 1 | 0.333 | 1 ==> 0 |
| | 148 | 1 | 0.500 | 0 --> 1 |
| | 169 | 1 | 0.250 | 0 ==> 1 |
| | 181 | 1 | 0.250 | 1 ==> 0 |
| node_148 --> node_141 | 18 | 1 | 0.200 | 1 ==> 0 |
| | 56 | 1 | 1.000 | 0 ==> 1 |
| | 149 | 1 | 1.000 | 0 ==> 1 |
| | 152 | 1 | 0.333 | 0 ==> 1 |
| | 198 | 1 | 0.200 | 1 ==> 0 |
| | 244 | 1 | 1.000 | 0 ==> 1 |
| | 260 | 2 | 0.500 | 0 ==> 2 |
| node_141 --> Eusthenopteron | 5 | 1 | 0.200 | 1 ==> 0 |
| | 8 | 1 | 0.167 | 1 ==> 0 |
| | 29 | 1 | 0.214 | 0 ==> 3 |
| | 112 | 1 | 0.250 | 0 ==> 1 |
| | 157 | 1 | 0.500 | 0 ==> 1 |
| | 258 | 1 | 0.333 | 1 --> 0 |
| | 266 | 1 | 0.333 | 0 ==> 1 |
| node_141 --> Gogonasus | 264 | 1 | 0.500 | 0 ==> 1 |
| node_148 --> node_147 | 31 | 1 | 0.091 | 0 ==> 1 |
| | 46 | 1 | 0.250 | 1 --> 0 |
| | 52 | 1 | 0.200 | 1 ==> 0 |
| | 55 | 1 | 1.000 | 0 ==> 1 |
| | 202 | 1 | 0.250 | 0 ==> 1 |
| | 203 | 1 | 0.333 | 0 ==> 1 |
| | 229 | 1 | 0.333 | 0 ==> 1 |
| | 235 | 1 | 1.000 | 0 ==> 1 |
| | 242 | 1 | 1.000 | 0 ==> 1 |
| node_147 --> node_143 | 48 | 1 | 0.286 | 0 --> 1 |
| | 81 | 1 | 0.143 | 0 --> 1 |
| | 119 | 1 | 0.250 | 0 ==> 1 |
| | 258 | 1 | 0.333 | 1 --> 0 |
| | 263 | 1 | 1.000 | 0 ==> 1 |
| | 264 | 1 | 0.500 | 0 ==> 1 |
| node_143 --> node_142 | 46 | 1 | 0.250 | 0 --> 1 |
| | 47 | 1 | 0.500 | 0 ==> 1 |
| | 112 | 1 | 0.250 | 0 ==> 1 |
| | 131 | 1 | 0.333 | 0 ==> 1 |
| node_142 --> Glyptolepis | 8 | 1 | 0.167 | 1 ==> 0 |
| | 18 | 1 | 0.200 | 1 ==> 0 |
| | 24 | 1 | 0.091 | 1 ==> 0 |
| | 29 | 1 | 0.214 | 0 ==> 3 |
| | 42 | 1 | 0.111 | 0 ==> 1 |
| | 67 | 1 | 0.250 | 1 ==> 0 |
| node_142 --> Porolepis | 198 | 1 | 0.200 | 1 ==> 0 |
| | 234 | 1 | 0.500 | 1 ==> 0 |
| node_143 --> Powichthys | 48 | 1 | 0.286 | 1 --> 2 |
| | 143 | 1 | 0.143 | 0 ==> 1 |
| | 146 | 1 | 0.200 | 1 ==> 0 |
| | 266 | 1 | 0.333 | 0 ==> 1 |
| node_147 --> node_146 | 42 | 1 | 0.111 | 0 ==> 1 |
| | 122 | 1 | 0.333 | 1 ==> 0 |
| | 148 | 1 | 0.500 | 1 --> 0 |
| | 167 | 1 | 0.500 | 1 ==> 0 |

| | | | | |
|---|---|---|---|---|
| | 254 | 1 | 1.000 | 0 ==> 1 |
| | 284 | 1 | 0.200 | 0 --> 1 |
| node_146 --> Youngolepis | 20 | 1 | 0.200 | 0 ==> 1 |
| | 54 | 1 | 0.333 | 1 ==> 0 |
| | 146 | 1 | 0.200 | 1 ==> 0 |
| node_146 --> node_145 | 24 | 1 | 0.091 | 1 --> 0 |
| | 67 | 1 | 0.250 | 1 --> 0 |
| | 85 (teeth ankyolosed) | 1 | 0.500 | 0 ==> 1 |
| | 86 | 1 | 0.500 | 1 ==> 0 |
| | 137 | 1 | 0.333 | 1 --> 0 |
| | 155 | 1 | 0.250 | 1 --> 0 |
| | 163 | 1 | 0.500 | 1 --> 0 |
| | 169 | 1 | 0.250 | 1 --> 0 |
| | 171 | 1 | 0.200 | 1 --> 0 |
| | 240 | 1 | 1.000 | 0 ==> 1 |
| | 241 | 1 | 1.000 | 0 ==> 1 |
| | 245 | 1 | 0.500 | 0 --> 1 |
| | 246 | 1 | 0.333 | 0 ==> 1 |
| | 247 | 1 | 1.000 | 0 ==> 1 |
| | 248 | 1 | 1.000 | 0 ==> 1 |
| | 255 | 1 | 0.500 | 1 --> 0 |
| | 260 | 1 | 0.500 | 0 ==> 1 |
| node_145 --> Diabolepis | 48 | 1 | 0.286 | 0 ==> 2 |
| | 112 | 1 | 0.250 | 0 ==> 1 |
| node_145 --> node_144 | 100 | 1 | 0.333 | 0 ==> 1 |
| | 117 | 1 | 0.286 | 1 ==> 0 |
| | 123 | 1 | 0.200 | 1 ==> 0 |
| | 166 (ventral cranial fissure) | 1 | 0.250 | 1 ==> 0 |
| | 168 | 1 | 0.250 | 1 --> 0 |
| | 237 | 1 | 0.200 | 1 ==> 0 |
| | 257 | 1 | 1.000 | 0 ==> 1 |
| | 259 | 1 | 1.000 | 0 ==> 1 |
| | 260 | 1 | 0.500 | 1 ==> 2 |
| | 262 | 1 | 0.500 | 1 ==> 2 |
| | 265 | 1 | 1.000 | 0 ==> 1 |
| node_144 --> Dipterus | 18 | 1 | 0.200 | 1 ==> 0 |
| node_144 --> Uranolophus | 27 | 1 | 0.167 | 0 ==> 1 |
| | 34 | 1 | 0.200 | 1 ==> 0 |
| | 78 | 1 | 0.167 | 1 ==> 0 |
| | 107 | 1 | 0.500 | 0 ==> 1 |
| | 172 | 1 | 0.200 | 1 ==> 0 |
| node_149 --> Styloichthys | 20 | 1 | 0.200 | 0 --> 1 |
| | 54 | 1 | 0.333 | 1 --> 0 |
| | 81 | 1 | 0.143 | 0 ==> 1 |
| | 119 | 1 | 0.250 | 0 ==> 1 |
| | 126 | 1 | 0.333 | 0 ==> 1 |
| | 130 | 1 | 0.333 | 0 ==> 1 |
| | 246 | 1 | 0.333 | 0 ==> 1 |
| | 250 | 1 | 0.500 | 0 ==> 1 |
| node_154 --> node_153 | 5 | 1 | 0.200 | 1 --> 0 |
| | 8 | 1 | 0.167 | 1 --> 0 |
| | 18 | 1 | 0.200 | 1 ==> 0 |
| | 24 | 1 | 0.091 | 1 ==> 0 |
| | 29 | 1 | 0.214 | 0 --> 3 |
| | 42 | 1 | 0.111 | 0 ==> 1 |
| | 67 | 1 | 0.250 | 1 ==> 0 |
| | 91 | 1 | 0.333 | 1 --> 0 |
| | 97 | 1 | 0.250 | 0 ==> 1 |
| | 132 | 1 | 0.333 | 1 --> 0 |
| | 198 | 1 | 0.200 | 1 --> 0 |
| | 225 | 1 | 0.500 | 1 --> 0 |
| | 233 | 1 | 0.250 | 0 --> 1 |
| | 253 | 1 | 0.500 | 0 --> 1 |
| | 270 | 1 | 0.250 | 0 --> 1 |
| node_153 --> node_150 | 57 | 1 | 0.333 | 0 --> 1 |
| | 81 | 1 | 0.143 | 0 ==> 1 |
| | 89 | 1 | 0.333 | 1 --> 0 |
| | 110 | 1 | 0.500 | 1 ==> 0 |
| | 116 | 1 | 0.250 | 1 --> 0 |
| | 119 | 1 | 0.250 | 0 ==> 1 |
| | 124 | 1 | 0.200 | 0 --> 1 |
| | 131 | 1 | 0.333 | 1 --> 0 |
| | 134 | 1 | 0.250 | 1 ==> 0 |
| | 137 | 1 | 0.333 | 1 --> 0 |
| | 143 | 1 | 0.143 | 0 --> 1 |
| | 169 | 1 | 0.250 | 0 ==> 1 |
| node_150 --> Onychodus | 92 | 1 | 0.333 | 0 ==> 1 |
| | 243 | 1 | 0.333 | 1 ==> 0 |
| | 280 | 1 | 0.250 | 1 ==> 0 |
| node_150 --> Qingmenodus | 29 | 1 | 0.214 | 3 --> 0 |
| | 45 | 1 | 0.250 | 0 ==> 1 |
| node_153 --> node_152 | 31 | 1 | 0.091 | 0 --> 1 |
| | 47 | 1 | 0.500 | 0 ==> 1 |
| | 48 | 1 | 0.286 | 0 --> 1 |

| | | | | |
|---|---|---|---|---|
| | 123 | 1 | 0.200 | 1 --> 0 |
| | 164 | 2 | 0.400 | 2 --> 0 |
| | 168 | 1 | 0.250 | 1 --> 0 |
| | 238 | 1 | 0.500 | 1 --> 0 |
| | 245 | 1 | 0.500 | 0 ==> 1 |
| | 246 | 1 | 0.333 | 0 ==> 1 |
| | 250 | 1 | 0.500 | 0 ==> 1 |
| | 251 | 1 | 1.000 | 0 ==> 1 |
| | 252 | 1 | 0.500 | 0 ==> 1 |
| | 261 | 1 | 1.000 | 0 ==> 1 |
| | 267 | 1 | 1.000 | 0 ==> 1 |
| node_152 --> node_151 | 181 | 1 | 0.250 | 1 --> 0 |
| node_151 --> Gavinia | 81 | 1 | 0.143 | 0 ==> 1 |
| node_152 --> Miguashaia | 4 | 1 | 0.333 | 1 ==> 0 |
| | 34 | 1 | 0.200 | 1 ==> 0 |
| | 48 | 1 | 0.286 | 1 --> 2 |
| | 117 | 1 | 0.286 | 1 ==> 0 |
| node_160 --> node_159 | 57 | 1 | 0.333 | 0 --> 1 |
| | 81 | 1 | 0.143 | 0 ==> 1 |
| | 92 | 1 | 0.333 | 0 ==> 1 |
| | 94 | 1 | 0.500 | 0 --> 1 |
| | 119 | 1 | 0.250 | 0 ==> 1 |
| | 134 | 1 | 0.250 | 1 ==> 0 |
| | 213 | 1 | 0.200 | 0 --> 1 |
| | 274 | 1 | 0.500 | 1 --> 0 |
| | 275 | 1 | 0.200 | 0 ==> 1 |
| node_159 --> Guiyu | 20 | 1 | 0.200 | 1 --> 0 |
| node_159 --> node_158 | 21 | 1 | 0.143 | 1 ==> 0 |
| | 34 | 1 | 0.200 | 0 --> 1 |
| | 253 | 1 | 0.500 | 0 --> 1 |
| node_158 --> node_157 | 29 | 1 | 0.214 | 1 ==> 0 |
| node_157 --> Achoania | 45 | 1 | 0.250 | 0 ==> 1 |
| | 89 | 1 | 0.333 | 0 ==> 1 |
| | 145 | 1 | 0.667 | 0 ==> 2 |
| node_161 --> "Ligulalepis" | 31 | 1 | 0.091 | 0 ==> 1 |
| | 270 | 1 | 0.250 | 0 ==> 1 |
| | 278 | 1 | 0.400 | 1 ==> 2 |
| | 281 | 1 | 0.500 | 1 --> 0 |
| | 282 | 1 | 0.500 | 1 --> 2 |
| node_162 --> Dialipina | 11 | 1 | 0.333 | 0 ==> 1 |
| | 19 | 1 | 0.500 | 0 --> 1 |
| | 48 | 1 | 0.286 | 0 ==> 1 |
| | 239 | 1 | 0.500 | 0 ==> 1 |
| node_164 --> Janusiscus | 140 | 1 | 0.333 | 0 ==> 1 |
| | 143 | 1 | 0.143 | 0 ==> 1 |
| | 266 | 1 | 0.333 | 0 ==> 1 |
| node_165 --> Ramirosuarezia | 104 | 1 | 0.333 | 0 ==> 1 |
| | 117 | 1 | 0.286 | 1 ==> 0 |
| | 124 | 1 | 0.200 | 0 ==> 1 |
| | 171 | 1 | 0.200 | 0 ==> 1 |
| node_166 --> Entelognathus | 24 | 1 | 0.091 | 1 --> 0 |
| | 43 | 1 | 0.250 | 1 ==> 0 |
| | 57 | 1 | 0.333 | 0 ==> 1 |
| | 70 | 1 | 0.250 | 0 ==> 1 |
| | 77 | 1 | 0.167 | 1 ==> 0 |
| | 88 | 1 | 0.500 | 0 ==> 1 |
| | 172 | 1 | 0.200 | 0 ==> 1 |
| | 177 | 1 | 0.200 | 0 ==> 1 |
| | 278 | 1 | 0.400 | 1 ==> 2 |
| | 279 | 1 | 0.500 | 1 ==> 0 |
| node_167 --> Minjinia | 4 | 1 | 0.333 | 0 ==> 1 |
| | 166 (ventral cranial fissure) | 1 | 0.250 | 0 ==> 1 |
| node_173 --> node_172 | 23 | 1 | 0.125 | 0 --> 1 |
| | 40 (endo w oblique) | 1 | 1.000 | 0 ==> 1 |
| | 109 | 1 | 0.500 | 0 ==> 1 |
| | 211 | 1 | 0.167 | 1 --> 0 |
| | 226 | 1 | 0.143 | 1 --> 0 |
| | 228 | 1 | 0.200 | 1 --> 0 |
| | 262 | 2 | 0.500 | 0 ==> 2 |
| | 284 | 1 | 0.200 | 0 --> 1 |
| node_172 --> node_171 | 24 | 1 | 0.091 | 1 --> 0 |
| | 60 | 1 | 0.750 | 0 --> 1 |
| | 237 | 1 | 0.200 | 0 ==> 1 |
| node_171 --> node_170 | 49 | 1 | 0.500 | 0 --> 1 |
| | 50 (medial process of paranuchal binary) | 1 | 0.500 | 0 ==> 1 |
| | 70 | 1 | 0.250 | 0 --> 1 |
| | 125 | 1 | 0.667 | 1 --> 0 |
| node_170 --> node_169 | 51 (paired pits) | 1 | 1.000 | 0 ==> 1 |
| | 186 | 1 | 1.000 | 0 ==> 1 |
| node_169 --> Buchanosteus | 43 | 1 | 0.250 | 1 ==> 0 |
| | 70 | 1 | 0.250 | 1 --> 0 |
| | 220 | 1 | 0.125 | 1 ==> 0 |
| node_169 --> node_168 | 44 | 1 | 0.500 | 0 ==> 1 |
| | 78 | 1 | 0.167 | 0 ==> 1 |

| | | | | |
|---|---|---|---|---|
| | 97 | 1 | 0.250 | 0 --> 1 |
| | 113 | 1 | 1.000 | 0 ==> 1 |
| | 284 | 1 | 0.200 | 1 ==> 0 |
| node_168 --> Incisoscutum | 42 | 1 | 0.111 | 1 ==> 0 |
| | 213 | 1 | 0.200 | 1 ==> 0 |
| | 280 | 1 | 0.250 | 1 ==> 0 |
| node_170 --> Cowralepis | 29 | 1 | 0.214 | 3 ==> 2 |
| | 31 | 1 | 0.091 | 0 ==> 1 |
| | 38 (extent of dermatocr) | 1 | 0.200 | 0 ==> 1 |
| | 39 (endolymph in sr) | 1 | 0.333 | 0 ==> 1 |
| | 60 | 1 | 0.750 | 1 --> 0 |
| | 173 | 1 | 0.250 | 0 ==> 1 |
| | 174 | 1 | 0.500 | 0 ==> 1 |
| | 183 (pectoral fenestra) | 1 | 0.250 | 0 ==> 1 |
| | 230 | 1 | 0.200 | 1 --> 0 |
| | 278 | 1 | 0.400 | 1 ==> 2 |
| | 280 | 1 | 0.250 | 1 ==> 0 |
| node_172 --> Kujdanowiaspis | 42 | 1 | 0.111 | 1 ==> 0 |
| node_174 --> Romundina | 5 | 1 | 0.200 | 0 ==> 1 |
| | 31 | 1 | 0.091 | 0 --> 1 |
| | 43 | 1 | 0.250 | 1 ==> 0 |
| | 114 | 1 | 0.333 | 1 ==> 0 |
| | 143 | 1 | 0.143 | 0 ==> 1 |
| | 145 | 1 | 0.667 | 0 --> 1 |
| | 152 | 1 | 0.333 | 0 ==> 1 |
| | 173 | 1 | 0.250 | 0 ==> 1 |
| | 270 | 1 | 0.250 | 0 --> 1 |
| node_175 --> Eurycaraspis | 44 | 1 | 0.500 | 0 ==> 1 |
| | 50 (medial process of paranuchal binary) | 1 | 0.500 | 0 ==> 1 |
| node_178 --> node_177 | 38 (extent of dermatocr) | 1 | 0.200 | 0 ==> 1 |
| | 70 | 1 | 0.250 | 0 ==> 1 |
| | 97 | 1 | 0.250 | 0 --> 1 |
| | 99 | 1 | 0.500 | 0 --> 1 |
| | 183 (pectoral fenestra) | 1 | 0.250 | 0 --> 1 |
| | 185 | 1 | 0.333 | 1 ==> 0 |
| | 208 | 1 | 1.000 | 0 --> 1 |
| | 233 | 1 | 0.250 | 0 --> 1 |
| node_177 --> node_176 | 220 | 1 | 0.125 | 1 --> 0 |
| | 237 | 1 | 0.200 | 0 --> 1 |
| node_176 --> Austroptyctodus | 181 | 1 | 0.250 | 0 ==> 1 |
| node_177 --> Rhamphodopsis | 219 | 1 | 0.200 | 0 ==> 1 |
| | 275 | 1 | 0.200 | 0 ==> 1 |
| node_180 --> node_179 | 87 | 1 | 0.333 | 1 --> 0 |
| | 116 | 1 | 0.250 | 0 --> 1 |
| | 117 | 1 | 0.286 | 1 --> 2 |
| | 144 | 1 | 0.333 | 0 --> 1 |
| | 174 | 1 | 0.500 | 0 ==> 1 |
| | 282 | 1 | 0.500 | 2 --> 0 |
| node_179 --> Lunaspis | 9 | 1 | 0.333 | 1 ==> 0 |
| | 29 | 1 | 0.214 | 3 ==> 2 |
| | 42 | 1 | 0.111 | 1 ==> 0 |
| | 219 | 1 | 0.200 | 0 ==> 1 |
| | 237 | 1 | 0.200 | 0 ==> 1 |
| node_181 --> Brindabellaspis | 23 | 1 | 0.125 | 0 --> 1 |
| node_187 --> node_186 | 13 | 1 | 0.333 | 1 ==> 0 |
| | 136 | 1 | 0.500 | 0 --> 1 |
| | 153 | 1 | 0.333 | 0 --> 1 |
| | 164 | 1 | 0.400 | 1 --> 2 |
| | 226 | 1 | 0.143 | 1 --> 0 |
| | 278 | 1 | 0.400 | 0 --> 1 |
| node_186 --> node_184 | 21 | 1 | 0.143 | 0 --> 1 |
| | 187 | 1 | 1.000 | 0 ==> 1 |
| | 188 | 1 | 1.000 | 0 ==> 1 |
| | 209 | 1 | 1.000 | 0 ==> 1 |
| | 224 | 1 | 0.250 | 1 --> 0 |
| | 277 | 1 | 1.000 | 1 ==> 0 |
| node_184 --> node_183 | 9 | 1 | 0.333 | 1 --> 0 |
| node_183 --> node_182 | 3 | 1 | 0.167 | 0 ==> 1 |
| | 78 | 1 | 0.167 | 0 --> 1 |
| | 210 | 1 | 1.000 | 0 ==> 1 |
| | 213 | 1 | 0.200 | 1 ==> 0 |
| node_182 --> Pterichthyodes | 211 | 1 | 0.167 | 0 ==> 1 |
| node_183 --> Parayunnanolepis | 26 | 1 | 0.400 | 0 ==> 1 |
| node_186 --> node_185 | 26 | 1 | 0.400 | 0 --> 2 |
| | 27 | 1 | 0.167 | 1 --> 0 |
| | 161 | 1 | 0.333 | 0 ==> 1 |
| | 183 (pectoral fenestra) | 1 | 0.250 | 0 --> 1 |
| | 185 | 1 | 0.333 | 1 ==> 0 |
| | 284 | 1 | 0.200 | 1 ==> 0 |

### Supplementary Tables

**Supplementary Table 1.** Model likelihoods and confidence intervals for ancestral states reconstructions on results from parsimony and Bayesian phylogenetic analysis. ER, equal rates model; ARD, all-rates-different model. Reports for marginal likelihoods of presence or absence of endochondral bone are given for the gnathostome crown node and the last common ancestor of *Minjinia* and the gnathostome crown.

| Trees | Model | Median | 2.5% | 25% | 75% | 97.5% |
|---|---|---|---|---|---|---|
| Parsimony Implied Weights | | log.lik. | | | | |
| | ER | -28.27 | -28.28 | -28.27 | -28.24 | -28.15 |
| | ARD | -26.15 | -26.16 | -26.16 | -26.12 | -26.02 |
| | | AIC | | | | |
| | ER | 58.53 | 58.3 | 58.47 | 58.55 | 58.55 |
| | ARD | 56.31 | 56.04 | 56.25 | 56.31 | 56.31 |
| | | *Minjinia*:Gnathostome absent | | | | |
| | ER | 0.86 | 0.85 | 0.85 | 0.86 | 0.86 |
| | ARD | 0.34 | 0.32 | 0.34 | 0.34 | 0.34 |
| | | *Minjinia*:Gnathostome present | | | | |
| | ER | 0.14 | 0.14 | 0.14 | 0.15 | 0.15 |
| | ARD | 0.66 | 0.66 | 0.66 | 0.66 | 0.68 |
| | | Gnathostome crown absent | | | | |
| | ER | 0.96 | 0.96 | 0.96 | 0.96 | 0.96 |
| | ARD | 0.59 | 0.59 | 0.59 | 0.59 | 0.59 |
| | | Gnathostome crown present | | | | |
| | ER | 0.04 | 0.04 | 0.04 | 0.04 | 0.04 |
| | ARD | 0.41 | 0.41 | 0.41 | 0.41 | 0.41 |
| Parsimony Equal Weights | | log.lik. | | | | |
| | ER | -28.91 | -28.97 | -28.93 | -28.88 | -28.84 |
| | ARD | -26.54 | -26.61 | -26.57 | -26.52 | -26.48 |
| | | AIC | | | | |
| | ER | 59.82 | 59.68 | 59.75 | 59.85 | 59.95 |
| | ARD | 57.09 | 56.96 | 57.05 | 57.13 | 57.23 |
| | | *Minjinia*:Gnathostome absent | | | | |
| | ER | 0.67 | 0.66 | 0.66 | 0.67 | 0.68 |
| | ARD | 0.19 | 0.18 | 0.18 | 0.19 | 0.2 |

| | | | | | | |
|---|---|---|---|---|---|---|
| | | *Minjinia*:Gnathostome present | | | | |
| | ER | 0.33 | 0.32 | 0.33 | 0.34 | 0.34 |
| | ARD | 0.81 | 0.8 | 0.81 | 0.82 | 0.82 |
| | | Gnathostome crown absent | | | | |
| | ER | 0.91 | 0.91 | 0.91 | 0.91 | 0.91 |
| | ARD | 0.46 | 0.45 | 0.46 | 0.47 | 0.48 |
| | | Gnathostome crown present | | | | |
| | ER | 0.09 | 0.09 | 0.09 | 0.09 | 0.09 |
| | ARD | 0.54 | 0.52 | 0.53 | 0.54 | 0.55 |
| Bayesian Partitioned | | log.lik. | | | | |
| | ER | -29.61 | -32.35 | -30.32 | -29.08 | -28.12 |
| | ARD | -27.34 | -29.71 | -27.95 | -26.81 | -25.62 |
| | | AIC | | | | |
| | ER | 61.22 | 58.24 | 60.17 | 62.65 | 66.71 |
| | ARD | 58.67 | 55.24 | 57.62 | 59.9 | 63.43 |
| | | *Minjinia*:Gnathostome absent | | | | |
| | ER | 0.9 | 0.08 | 0.58 | 1 | 1 |
| | ARD | 0.37 | 0 | 0.06 | 0.94 | 1 |
| | | *Minjinia*:Gnathostome present | | | | |
| | ER | 0.1 | 0 | 0 | 0.42 | 0.92 |
| | ARD | 0.63 | 0 | 0.06 | 0.94 | 1 |
| | | Gnathostome crown absent | | | | |
| | ER | 0.88 | 0.05 | 0.69 | 0.95 | 0.99 |
| | ARD | 0.36 | 0 | 0.13 | 0.67 | 0.91 |
| | | Gnathostome crown present | | | | |
| | ER | 0.12 | 0.01 | 0.05 | 0.31 | 0.95 |
| | ARD | 0.64 | 0.09 | 0.33 | 0.87 | 1 |
| Bayesian Unpartitioned | | log.lik. | | | | |
| | ER | -29.66 | -32.09 | -30.28 | -28.64 | -27.6 |
| | ARD | -27.03 | -29.2 | -27.88 | -26.31 | -24.93 |
| | | AIC | | | | |
| | ER | 61.32 | 57.21 | 59.27 | 62.56 | 66.19 |
| | ARD | 58.06 | 53.86 | 56.62 | 59.75 | 62.4 |
| | | *Minjinia*:Gnathostome absent | | | | |

| | | | | | |
|---|---|---|---|---|---|
| ER | 0.73 | 0.03 | 0.34 | 0.98 | 1 |
| ARD | 0.17 | 0 | 0.02 | 0.76 | 1 |
| | *Minjinia*:Gnathostome present | | | | |
| ER | 0.27 | 0 | 0.02 | 0.66 | 0.97 |
| ARD | 0.83 | 0 | 0.24 | 0.98 | 1 |
| | Gnathostome crown absent | | | | |
| ER | 0.79 | 0.05 | 0.47 | 0.92 | 0.99 |
| ARD | 0.22 | 0 | 0.05 | 0.46 | 0.89 |
| | Gnathostome crown present | | | | |
| ER | 0.21 | 0.01 | 0.08 | 0.53 | 0.95 |
| ARD | 0.78 | 0.11 | 0.54 | 0.95 | 1 |
