## Supplementary Data 1 for "Endochondral bone in an Early Devonian ‘placoderm’ from Mongolia": convert.homoplasy.R

```
#' @title Convert homoplasy
#'
#' @description converts a "matrix-list" (TNT output) of homoplasic characters into a a list of homoplasies
#'
#' @param input the TNT file, tab delimited.
#' @param header whether the file has a header (\code{TRUE} - default) or not (\code{FALSE}).
#' @param first.character.zero whether the first character is 0 (\code{TRUE}) or 1 (\code{FALSE} - default).
#' @param mb.out whether to output (print) a MrBayes constraint (\code{TRUE}) or not (\code{FALSE} - default).
#' @param combine.uninfo whether to combine unfinformative characters into the same partition (\code{TRUE}) or into different ones (\code{FALSE} - default).
#' @param export optional, the name/path of a file where to export the MrBayes partitions.
#'  
#' @examples
#'
#' @seealso
#' 
#' @author Thomas Guillerme
#' @export

# input <- "tnt_homo.txt"

convert.homoplasy <- function(input, header = TRUE, first.character.zero = FALSE, mb.out = FALSE, combine.uninfo = FALSE, export) {

    ## Get the input
    input_mess <- read.csv(input)

    ## Clean the input
    ## Remove the rows that are just spaces
    empty_rows <- !apply(input_mess, 1, function(x)(length(grep("[0-9]", x)) > 0))
    if(any(empty_rows)) {
        input_mess <- as.list(input_mess[-empty_rows,])
    }

    ## Split the characters
    split.fun <- function(x) {
        ## Get the digits only
        splits <- strsplit(as.character(x), split = "\\s+")[[1]]

        options(warn = -1)
        ## Remove the first two elements and convert to numeric
        splits <- as.numeric(splits[-c(1,2)])
        options(warn = 0)

        ## Remove the first digit
        return(splits)
    }

    ## Get all characters homoplasy score
    homoplasy_vector <- unlist(lapply(input_mess, split.fun))

    ## Sort characters
    sort.homoplasy <- function(homoplasy, characters) {
        return(which(characters == homoplasy))
    }

    ## Get the list of characters sorted per homoplasy score
    homoplasy_list <- lapply(as.list(na.omit(unique(homoplasy_vector))), sort.homoplasy, characters = homoplasy_vector)
    names(homoplasy_list) <- na.omit(unique(homoplasy_vector))

    ## Add uniformative characters
    if(any(is.na(homoplasy_vector))) {
        ## Get the uninformative characters
        uninformative_characters <- which(is.na(homoplasy_vector))
        if(combine.uninfo) {
            homoplasy_list <- c(homoplasy_list, list("uninformative" = c(uninformative_characters)))
        } else {
            homoplasy_list <- c(homoplasy_list, "uninformative" = uninformative_characters)
        }
    }

    ## Convert into 0 base (i.e. first character is called character 0)
    if(first.character.zero) {
        homoplasy_list <- lapply(homoplasy_list, function(x) return(x-1))
    }

    ## Check if export is required
    if(!missing(export)) {
        export_path <- export
        do_export <- TRUE
    } else {
        do_export <- FALSE
    }


    ## Make the MrBayes partitions
    if(mb.out){
        for(score in 1:length(homoplasy_list)) {
            if(do_export) sink(export_path, append = TRUE)
            cat(paste0(paste0("charset homoplasy_", names(homoplasy_list)[score], " = "), paste(homoplasy_list[[score]], collapse = " "), ";\n"))
            if(do_export) sink()
        }
    }

    return(homoplasy_list)
}
```
