## Supplementary Data 2 for "Endochondral bone in an Early Devonian ‘placoderm’ from Mongolia": ACE.R

```
## plotting the ancestral character estimation
#' @param tree the tree to plot
#' @param character one character as output from read.nexus.data()
#' @param algorithm whether to use the "phytools" one (rerootingMethod) or the "castor" one (asr_mk_model - which is the same as "ape::ace" but much faster!).
#' @param model the ancestral character estimation model to be passed to phytools::rerootinMethod or castor::asr_mk_model. This can be a model name or a transition matrix.
#' 
#' 
## running the ancestral character estimation
run.ACE <- function(tree, character, algorithm, model = "ER") {

    ## Order the character to match the one from the input tree
    character <- character[match(tree$tip.label, names(character))]

    ## Get the character's tokens
    tokens <- unique(character)
    options(warn = -1)
    token_value <- as.numeric(tokens)
    options(warn = 0)

    ## Make the state table
    state_table <- matrix(0, ncol = length(na.omit(token_value)), nrow = length(character))
    colnames(state_table) <- na.omit(token_value)

    ## Fill the state table
    fill.table <- function(taxa, tokens) {
        ## Match the taxa character with the right token
        taxa_char <- as.numeric(tokens %in% taxa)
        ## If no match, make equiprobalbe
        if(sum(taxa_char) == 0) {
            return(rep(1/length(tokens), length(tokens)))
            #return(rep(1.0, length(tokens))) # This is the corrected tip partials
        } else {
            return(taxa_char)
        }
    }
    state_table <- t(sapply(character, fill.table, tokens = as.character(na.omit(token_value))))
    colnames(state_table) <- as.character(na.omit(token_value))

    ## Do the ancestral estimation
    if(algorithm == "phytools") {
        ancestral_estimates <- phytools::rerootingMethod(tree, state_table, model = model, tips = FALSE)
        ## Get the node probabilities
        ancestral_nodes <- ancestral_estimates$marginal.anc
        ## Get the model likelihood
        log.lik <- ancestral_estimates$loglik
        ## Get the transition matrix
        transition <- ancestral_estimates$Q
    }
    if(algorithm == "castor") {
        ancestral_estimates <- castor::asr_mk_model(tree = tree, tip_states = NULL, Nstates = length(na.omit(token_value)), rate_model = model, Ntrials = 3, tip_priors = state_table)
        ## Get the node probabilities
        ancestral_nodes <- ancestral_estimates$ancestral_likelihoods
        ## Get the model likelihood
        log.lik <- ancestral_estimates$loglikelihood
        ## Get the transition matrix
        transition <- ancestral_estimates$transition_matrix
    }

    return(list("tips" = state_table, "nodes" = ancestral_nodes, "log.lik" = log.lik, "transition" = transition))
}

## plotting the ancestral character estimation
#' @param tree_plot the tree to plot
#' @param ace the tips and nodes probabilities from run.ACE()
#' @param tree_ace the tree used for the ACE
#' @param col the colours for the different tokens (should match the number of tokens for clarity!)
#' @param type how to represent the tips and nodes (either "pie", or "thermo" - see ?ape::nodelabels)
#' @param type.size the size of the pies or thermos
#' @param tip.size the size of the tip labels
#' @param state.names optional, the name of the states (rather than 0 and 1!)
#' @param legend.pos the x,y coordinates for the legend (is bottom left by default). If set to NULL, it doesn't plot the legend anywhere.
#' @param ... any option to be passed to plot.phylo (tip labels, main, etc...)
plot.ACE <- function(tree_plot, ace, trees_ace, col = c("blue", "orange"), type = "pie", type.size = 0.5, tip.size = 0.5, state.names, legend.pos, ...) {

    ## Plot the tree_plot
    # Don't plot with branch lengths to retain readability (requires plot.phylo disambiguation, I think)
    ## ATTN: THOMAS: See if there is a way to adjust tip labels so they aren't overlapped by the pies.
    plot(tree_plot, cex = 1.0, use.edge.length=FALSE, label.offset=1.0, node.depth = 2)

    ## Match the tip labels with the ace tips
    # ace$tip <- ace$tip[match(tree_plot$tip.label, rownames(ace$tip)), ]

    ## Node in tree_plots
    get.node.ace <- function(ace, tree_plot, trees_ace) {
        ## Extract the tips per clade
        get.tips.clade <- function(node, tree_plot) {
            return(extract.clade(tree_plot, node = Ntip(tree_plot) + node)$tip.label)
        }

        ## Get all tips
        all_tips <- lapply(as.list(1:Nnode(tree_plot)), get.tips.clade, tree_plot)

        ## Get the nodes of interest in the tree_ace
        get.MRCAs <- function(phy, tips) {
            return(unlist(lapply(tips, function(tip, phy) getMRCA(phy, tip), phy = phy)) - Ntip(phy))
        }
        nodes_list <- lapply(trees_ace, get.MRCAs, tips = all_tips)

        ## Getting the ace values per nodes
        get.ace <- function(one_ace, nodes) {return(one_ace$node[nodes, 1])}

        ## Get the median values for these nodes
        return(apply(do.call(rbind, mapply(get.ace, ace, nodes_list, SIMPLIFY = FALSE)), 2, median))
    }

    ## Add the nodes and tips values
    if(type == "pie") {
        tiplabels(pie = ace[[1]]$tip[,1][match(tree_plot$tip.label, names(ace[[1]]$tip[,1]))], piecol = col, cex = 0.5)
        nodelabels(pie = get.node.ace(ace, tree_plot, trees_ace), piecol = col, cex = type.size)
    }

    if(type == "thermo") {
        tiplabels(thermo = ace[[1]]$tip[,1][match(tree_plot$tip.label, names(ace[[1]]$tip[,1]))], piecol = col, cex = type.size)
        nodelabels(thermo = get.node.ace(ace, tree_plot, trees_ace), piecol = col, cex = type.size)
    }

    ## Add the legend
    if(missing(state.names)) {
        legend_text <- paste("state", colnames(ace$tip))
    } else {
        legend_text <- state.names
    }
    if(missing(legend.pos)) {
        legend.pos <- "bottomleft"
    }

    if(!is.null(legend.pos)) {
        legend(legend.pos, legend = legend_text, pch = 19, col = col, title = "Probabilities:")
    }
}
```
