## Supplementary Data 2 for "Endochondral bone in an Early Devonian ‘placoderm’ from Mongolia": Minjinia.ace.Rmd

````
---
title: "Ancestral character estimations of endochondral bone presence (character 4)"
author: "Thomas Guillerme"
date: "`r Sys.Date()`"
output:
  pdf_document:
    fig_width: 8
    fig_height: 8
    keep_tex: false
  self_contained: true
---

# Data and package

```{r, echo = FALSE}
library(ape)
library(knitr)
## Optional install of castor 1.5.7
#castorurl<-"https://cran.r-project.org/src/contrib/Archive/castor/castor_1.5.7.tar.gz"
#install.packages(castorurl, repos=NULL, type="source")
library(castor)
source("ACE.R")
```

```{r, echo = FALSE}
set.seed(42)
## Replacing the spaces in the polymorphisms (e.g. {0 1} -> {01})
system("sed -E 's/{([0-9]) /{\\1/g' ../Data/Minjinia_matrix.nex > ../Data/Minjinia_matrix.tmp")
## This creates a Minjinia_matrix.tmp file that's removed a couple of lines below

## The matrix
matrix <- do.call(rbind, read.nexus.data("../Data/Minjinia_matrix.tmp"))

## Remove the temporary file
system("rm ../Data/Minjinia_matrix.tmp")

## Correct the Ligulalepis name
wrong_label <- grep("Ligulalepis", rownames(matrix))
if(length(wrong_label) != 0) {
    rownames(matrix)[wrong_label] <- "Ligulalepis"
}

## The trees
pars_impweigh_conse <- read.nexus("../Data/parsimony_trees/piwe3consens.tre")
pars_standard_conse <- read.nexus("../Data/parsimony_trees/stdparsconsens.tre")
pars_impweigh_trees <- read.nexus("../Data/parsimony_trees/piwe3brlens.tre")
pars_standard_trees <- read.nexus("../Data/parsimony_trees/stdparsbrlens.tre")
baye_part_conse     <- read.nexus("../Data/partitioned/Minjinia_MrBayes_partitioned.nex.con.tre")
baye_unpa_conse     <- read.nexus("../Data/unpartitioned/Minjinia_MrBayes_unpartitioned.nex.con.tre")
baye_part_trees1    <- read.nexus("../Data/partitioned/Minjinia_MrBayes_partitioned.nex.run1.t")
baye_part_trees2    <- read.nexus("../Data/partitioned/Minjinia_MrBayes_partitioned.nex.run2.t")
baye_unpa_trees1    <- read.nexus("../Data/unpartitioned/Minjinia_MrBayes_unpartitioned.nex.run1.t")
baye_unpa_trees2    <- read.nexus("../Data/unpartitioned/Minjinia_MrBayes_unpartitioned.nex.run2.t")

## Make everything multiphylo
as.multiPhylo <- function(tree) {
    tree <- list(tree)
    class(tree) <- "multiPhylo"
    return(tree)
}
pars_impweigh_conse <- as.multiPhylo(pars_impweigh_conse)
pars_standard_conse <- as.multiPhylo(pars_standard_conse)
baye_part_conse     <- as.multiPhylo(baye_part_conse)
baye_unpa_conse     <- as.multiPhylo(baye_unpa_conse)

## Select 100 trees from the parsimony distribution
pars_standard_trees <- pars_standard_trees[sample(1:length(pars_standard_trees), 100)]

## Select 2*50 trees from the Bayesian distributions (ignoring the 25% first trees each time)
sample.bayesian <- function(trees1, trees2, burnin = 0.25, out = 100) {
    return(c(trees1[sample(burnin*length(trees1):length(trees1), 50)],
             trees2[sample(burnin*length(trees2):length(trees2), 50)]))
}
baye_part_trees <- sample.bayesian(baye_part_trees1, baye_part_trees2)
baye_unpa_trees <- sample.bayesian(baye_unpa_trees1, baye_unpa_trees2)

## Root and dichotomise the trees (just for ACE, they're actually already rooted)
## And also fix the Ligulalepis name
root.tree <- function(tree, outgroup = "Galeaspida") {
    fix.name <- function(tree) {
        wrong_label <- grep("Ligulalepis", tree$tip.label)
        if(length(wrong_label) != 0) {
            tree$tip.label[wrong_label] <- "Ligulalepis"
        }
        return(tree)
    }

    tree <- lapply(tree, fix.name)
    class(tree) <- "multiPhylo"

    add.node.label <- function(tree) {
        tree$node.label <- seq(Ntip(tree):(Ntip(tree)+Nnode(tree)))
        return(tree)
    }

    tree <- lapply(tree, add.node.label)
    class(tree) <- "multiPhylo"    

    return(root(tree, outgroup = outgroup, resolve.root = TRUE))
}
pars_impweigh_conse <- root.tree(pars_impweigh_conse)
pars_standard_conse <- root.tree(pars_standard_conse)
pars_impweigh_trees <- root.tree(pars_impweigh_trees)
pars_standard_trees <- root.tree(pars_standard_trees)
baye_part_conse     <- root.tree(baye_part_conse)
baye_unpa_conse     <- root.tree(baye_unpa_conse)
baye_part_trees     <- root.tree(baye_part_trees)
baye_unpa_trees     <- root.tree(baye_unpa_trees)
```

# Running the ancestral character estimation (ACE)

Here we run the ancestral character estimation with an equal rates (ER) and all rates different (ARD) on a random parsimony tree and a random resolution (dichotomous) of the Bayesian consensus tree.

```{r, echo = FALSE}
## ACE Wrapper
run.models <- function(tree, matrix) {
    ER  <- lapply(tree, run.ACE, character = matrix[, 4], algorithm = "castor", model = "ER")
    ARD <- lapply(tree, run.ACE, character = matrix[, 4], algorithm = "castor", model = "ARD")
    return(list("ER" = ER, "ARD" = ARD))
}

ace_pars_impweigh_trees <- run.models(pars_impweigh_trees, matrix)
ace_pars_standard_trees <- run.models(pars_standard_trees, matrix)
ace_baye_part_trees     <- run.models(baye_part_trees, matrix)
ace_baye_unpa_trees     <- run.models(baye_unpa_trees, matrix)
ace_pars_impweigh_conse <- run.models(pars_impweigh_conse, matrix)
ace_pars_standard_conse <- run.models(pars_standard_conse, matrix)
ace_baye_part_conse     <- run.models(baye_part_conse, matrix)
ace_baye_unpa_conse     <- run.models(baye_unpa_conse, matrix)
```

## Extracting the models informations

Extracting the AICs, logged likelihood (global) and the scaled likelihood for each state at the node of interest (MRCA between _Minjinia_ and _Helodus_) for all the ancestral character estimations.

```{r, echo = FALSE}
## Function for extracting the values of a list of ACEs
extract.ace <- function(ace.list, what, ...) {
    return(unlist(lapply(ace.list, what, ...)))
}
## Calculating both AICs
get.AIC <- function(data, param) {
    return(2*param - 2*data$log.lik)
}
## Get the model likelihoods
get.log.lik <- function(data) {
    return(data$log.lik)
}
## Get the node likelihoods
get.node.lik <- function(data, tree) {
    node_Minjinia <- getMRCA(tree, tip = c("Minjinia", "Helodus"))-Ntip(tree)
    node_Eusthenopteron <- getMRCA(tree, tip = c("Eusthenopteron", "Helodus"))-Ntip(tree)
    return(data$nodes[c(node_Minjinia, node_Eusthenopteron), ])
}

## Extract everything form the results
extract.results <- function(ace_results, trees) {
    ## log. lik.
    loglik <- lapply(ace_results, extract.ace, what = get.log.lik)
    ## AICs
    AICs <- list("ER"  = extract.ace(ace_results$ER, what = get.AIC, param = 1),
                 "ARD" = extract.ace(ace_results$ARD, what = get.AIC, param = 2))    
    ## Node lik.
    nodlik <- list("ER"  = mapply(get.node.lik, ace_results$ER, trees),
                   "ARD" = mapply(get.node.lik, ace_results$ARD, trees))
    nodlik_Minjinia0 <- lapply(nodlik, function(x) return(x[1,]))
    nodlik_Minjinia1 <- lapply(nodlik, function(x) return(x[3,]))
    nodlik_Eusthenopteron0 <- lapply(nodlik, function(x) return(x[2,]))
    nodlik_Eusthenopteron1 <- lapply(nodlik, function(x) return(x[4,]))

    return(list("log.lik." = loglik, "AIC" = AICs,
                "Minjinia:Gnathostome absent" = nodlik_Minjinia0,
                "Minjinia:Gnathostome present" = nodlik_Minjinia1,
                "Gnathostome crown absent" = nodlik_Eusthenopteron0,
                "Gnathostome crown present" = nodlik_Eusthenopteron1
                ))
}

results_pars_impweigh <- extract.results(ace_pars_impweigh_trees, pars_impweigh_trees)
results_pars_standard <- extract.results(ace_pars_standard_trees, pars_standard_trees)
results_baye_part     <- extract.results(ace_baye_part_trees, baye_part_trees)
results_baye_unpa     <- extract.results(ace_baye_unpa_trees, baye_unpa_trees)
```

## Result tables

Displaying the results into a table

```{r, echo = FALSE}
## Summarising the distributions
summary.results <- function(results, digits = 2, full = TRUE){
    ## Summarise the distributions
    summary.distribution <- function(distribution, cent.tend = median,
                                     probs = c(0.025, 0.25, 0.75, 0.975)) {
        return(c(cent.tend(distribution), quantile(distribution, prob = probs)))
    }

    ## Run the summaries
    if(full) {
        summarised_results <- lapply(lapply(results, lapply, summary.distribution),
                                     function(x) do.call(rbind, x))

        ## Make it into output tables
        table_results <- lapply(summarised_results, function(x, digits) round(x, digits = digits), digits)
        table_names <- lapply(as.list(names(summarised_results)), function(x) c(x, rep("", 4)))
        return(do.call(rbind, mapply(rbind, table_names, table_results, SIMPLIFY = FALSE)))
    } else {
        summarised_results <- do.call(cbind, lapply(lapply(results, lapply, median), function(x) do.call(rbind, x)))
        colnames(summarised_results) <- names(results)
        return(summarised_results)
    }
}

table_pars_impweigh <- summary.results(results_pars_impweigh)
table_pars_standard <- summary.results(results_pars_standard)
table_baye_part     <- summary.results(results_baye_part)
table_baye_unpa     <- summary.results(results_baye_unpa)

## Combine all results
all_results <- rbind(table_pars_impweigh, table_pars_standard, table_baye_part, table_baye_unpa)

## Add the models, trees nodes
models <- rownames(all_results)
trees <- c("Parsimony Implied Weights", "", "", rep("", 3*5),
           "Parsimony Equal Weights", "", "", rep("", 3*5),
           "Bayesian Partitioned", "", "", rep("", 3*5),
           "Bayesian Unpartitioned", "", "", rep("", 3*5))
nodes <- c("Minjinia:crown gnathostomes", rep("", nrow(all_results)-1))


## Full table
full_results <- cbind(do.call(cbind, list(trees, models)), all_results)
rownames(full_results) <- NULL
colnames(full_results)[1:3] <- c("Trees", "Model", "Median")

# ## Neat results (old)
# neat_results <- round(do.call(rbind,list(summary.results(results_pars_impweigh, full = FALSE),
#                                    summary.results(results_pars_standard, full = FALSE),
#                                    summary.results(results_baye_part, full = FALSE),
#                                    summary.results(results_baye_unpa, full = FALSE))), digits = 2)
# ## Add log lik ratio
# log.ratio <- function(logs, neat_results) return(c(2*abs(diff(neat_results[logs,1])), ""))
# neat_results <- cbind(cbind("log.lik."  = neat_results[, 1],
#                             "log.ratio" = unlist(lapply(list(c(1,2), c(3,4), c(5,6), c(7,8)), log.ratio, neat_results))),
#                             neat_results[,-1])

# models <- rownames(neat_results)
# trees <- c("Parsimony Implied Weights", "",
#            "Parsimony Equal Weights", "",
#            "Bayesian Partitioned", "",
#            "Bayesian Unpartitioned", "")
# neat_results <- cbind(do.call(cbind, list(trees, models)), neat_results)
# rownames(neat_results) <- NULL
# colnames(neat_results)[1:2] <- c("Trees", "Model")


## Neat results (to be similar format than in the manuscript)
parsimony <- round(summary.results(results_pars_standard, full = FALSE), 2)
bayesian  <- round(summary.results(results_baye_unpa, full = FALSE), 2)

## Get the node values and tree diagnosis
neat_table <- cbind(## Tree diagnosis
                    rbind(rbind(parsimony[,1:2], matrix("", 2, 2)),
                          rbind(bayesian[,1:2], matrix("", 2, 2))
                          ),
                    ## Node values
                    rbind(rbind(parsimony[,3:4], parsimony[,5:6]),
                          rbind(bayesian[,3:4], bayesian[,5:6])
                          )
                    )
colnames(neat_table)[c(3,4)] <- c("Absent", "Present")
rownames(neat_table) <- NULL

## Add the log.lik rations
log.ratio <- function(logs, neat_results) return(c(2*abs(diff(as.numeric(neat_results[logs,1]))), rep("", 3)))
neat_table <- cbind(cbind("log.lik."  = neat_table[, 1],
                            "log.ratio" = unlist(lapply(list(c(1,2), c(5,6)), log.ratio, neat_table))),
                            neat_table[,-1])

## Add the model, node and tree names
neat_table <- cbind("trees" = c("Parsimony", rep("", 3), "Bayesian", rep("", 3)),
                    "model" = rep(c("ER", "ARD"), 4),
                    neat_table[, c(1,2,3)],
                    "node"  = rep(c("Minjinia:Gnathostomes", "", "Crown Gnathostomes", ""), 2),
                    neat_table[, -c(1,2,3)])

## Results tables
caption1 <- "Tree distribution (100) ancestral states estimation results. ER = Equal rates model; ARD = All Rates Different model. The columns AIC and log.lik represent the median AIC and log.lik across the 100 parsimony and bayesian trees (for both models). The columns Absent and Present represent the median scaled likelihood for the endochondral bone state."
caption2 <- "The 2.5%, 25%, 75% and 97.5% represent the confidence intervals values for each estimated parameter (log.lik, AIC, Present and Absent endochondral bone)."
knitr::kable(full_results, caption = paste(caption1, caption2))
knitr::kable(neat_table, caption = caption1)
```


```{r eval = TRUE, echo = FALSE}
## This snippet exports the tables in latex format and converts them in docx

## Get the tables
write(knitr::kable(full_results, digits = 2, format = "latex"), "full_results.tex")
write(knitr::kable(neat_table, digits = 2, format = "latex"), "neat_results.tex")

## Make a pandoc converter snippet
system("echo \"pandoc -i table_latex_convert.tex -o result.tables.docx\" > run.pandoc.sh")
system("sh run.pandoc.sh")
system("rm run.pandoc.sh")
```

## Visualisation on the consensus trees

We can then visualise all the results on the consensus trees

```{r, fig.height = 18, fig.width = 14, echo = FALSE}
## Make this true for printing
par(mfrow = c(1,2))

## vis parameters
label_offset = 
tip_size = 0.5
type_size = 0.5

## Plotting a nicer tree
plot.ACE(tree_plot = ladderize(pars_impweigh_conse[[1]]),
         use.edge.length = FALSE,
         label.offset = 2,
         tip.size = tip_size,
         type.size = type_size,
         ace = ace_pars_impweigh_trees$ARD,
         trees_ace = pars_impweigh_trees,
         main = paste0("Endochondral bone estimation\n(parsimony implied weights - ARD)"),
         state.names = c("Absent", "Present")
         )
## Plotting a nicer tree
plot.ACE(tree_plot = ladderize(pars_impweigh_conse[[1]]),
         use.edge.length = FALSE,
         label.offset = 2,
         tip.size = tip_size,
         type.size = type_size,
         ace = ace_pars_impweigh_trees$ER,
         trees_ace = pars_impweigh_trees,
         main = paste0("Endochondral bone estimation\n(parsimony implied weights - ER)"),
         state.names = c("Absent", "Present")
         )
```


```{r, fig.height = 18, fig.width = 14, echo = FALSE}
par(mfrow = c(1,2))
## Plotting a nicer tree
plot.ACE(tree_plot = ladderize(pars_standard_conse[[1]]),
         use.edge.length = FALSE,
         label.offset = 2,
         tip.size = tip_size,
         type.size = type_size,
         ace = ace_pars_standard_trees$ARD,
         trees_ace = pars_standard_trees,
         main = paste0("Endochondral bone estimation\n(parsimony unweighted - ARD)"),
         state.names = c("Absent", "Present")
         )
## Plotting a nicer tree
plot.ACE(tree_plot = ladderize(pars_standard_conse[[1]]),
         use.edge.length = FALSE,
         label.offset = 2,
         tip.size = tip_size,
         type.size = type_size,
         ace = ace_pars_standard_trees$ER,
         trees_ace = pars_standard_trees,
         main = paste0("Endochondral bone estimation\n(parsimony unweighted - ER)"),
         state.names = c("Absent", "Present")
         )
```

```{r, fig.height = 18, fig.width = 14, echo = FALSE}
## Make this true for printing
par(mfrow = c(1,2))
## Plotting a nicer tree
plot.ACE(tree_plot = ladderize(baye_part_conse[[1]]),
         use.edge.length = FALSE,
         label.offset = 2,
         tip.size = tip_size,
         type.size = type_size,
         ace = ace_baye_part_trees$ARD,
         trees_ace = baye_part_trees,
         main = paste0("Endochondral bone estimation\n(Bayesian partitioned - ARD)"),
         state.names = c("Absent", "Present")
         )
## Plotting a nicer tree
plot.ACE(tree_plot = ladderize(baye_part_conse[[1]]),
         use.edge.length = FALSE,
         label.offset = 2,
         tip.size = tip_size,
         type.size = type_size,
         ace = ace_baye_part_trees$ER,
         trees_ace = baye_part_trees,
         main = paste0("Endochondral bone estimation\n(Bayesian partitioned - ER)"),
         state.names = c("Absent", "Present")
         )
```

```{r, fig.height = 18, fig.width = 14, echo = FALSE}
## Make this true for printing
par(mfrow = c(1,2))
## Plotting a nicer tree
plot.ACE(tree_plot = ladderize(baye_unpa_conse[[1]]),
         use.edge.length = FALSE,
         label.offset = 2,
         tip.size = tip_size,
         type.size = type_size,
         ace = ace_baye_unpa_trees$ARD,
         trees_ace = baye_unpa_trees,
         main = paste0("Endochondral bone estimation\n(Bayesian unpartitioned - ARD)"),
         state.names = c("Absent", "Present")
         )
## Plotting a nicer tree
plot.ACE(tree_plot = ladderize(baye_unpa_conse[[1]]),
         use.edge.length = FALSE,
         label.offset = 2,
         tip.size = tip_size,
         type.size = type_size,
         ace = ace_baye_unpa_trees$ER,
         trees_ace = baye_unpa_trees,
         main = paste0("Endochondral bone estimation\n(Bayesian unpartitioned - ER)"),
         state.names = c("Absent", "Present")
         )
```


```{r, fig.height = 20, fig.width = 14, echo = FALSE}
pdf("Parsimony_implied_weights.pdf", height = 20, width = 14)
## Make this true for printing
par(mfrow = c(1,2))
## Plotting a nicer tree
plot.ACE(tree_plot = ladderize(pars_impweigh_conse[[1]]),
         use.edge.length = FALSE,
         label.offset = 2,
         tip.size = tip_size,
         type.size = type_size,
         ace = ace_pars_impweigh_trees$ARD,
         trees_ace = pars_impweigh_trees,
         main = paste0("Endochondral bone estimation\n(parsimony implied weights - ARD)"),
         state.names = c("Absent", "Present")
         )
## Plotting a nicer tree
plot.ACE(tree_plot = ladderize(pars_impweigh_conse[[1]]),
         use.edge.length = FALSE,
         label.offset = 2,
         tip.size = tip_size,
         type.size = type_size,
         ace = ace_pars_impweigh_trees$ER,
         trees_ace = pars_impweigh_trees,
         main = paste0("Endochondral bone estimation\n(parsimony implied weights - ER)"),
         state.names = c("Absent", "Present")
         )
dev.off()
```


```{r, fig.height = 20, fig.width = 14, echo = FALSE}
pdf("Parsimony_equal_weights.pdf", height = 20, width = 14)
par(mfrow = c(1,2))
## Plotting a nicer tree
plot.ACE(tree_plot = ladderize(pars_standard_conse[[1]]),
         use.edge.length = FALSE,
         label.offset = 2,
         tip.size = tip_size,
         type.size = type_size,
         ace = ace_pars_standard_trees$ARD,
         trees_ace = pars_standard_trees,
         main = paste0("Endochondral bone estimation\n(parsimony unweighted - ARD)"),
         state.names = c("Absent", "Present")
         )
## Plotting a nicer tree
plot.ACE(tree_plot = ladderize(pars_standard_conse[[1]]),
         use.edge.length = FALSE,
         label.offset = 2,
         tip.size = tip_size,
         type.size = type_size,
         ace = ace_pars_standard_trees$ER,
         trees_ace = pars_standard_trees,
         main = paste0("Endochondral bone estimation\n(parsimony unweighted - ER)"),
         state.names = c("Absent", "Present")
         )
dev.off()
```


```{r, fig.height = 20, fig.width = 14, echo = FALSE}
pdf("Bayesian_parititioned.pdf", height = 20, width = 14)
## Make this true for printing
par(mfrow = c(1,2))
## Plotting a nicer tree
plot.ACE(tree_plot = ladderize(baye_part_conse[[1]]),
         use.edge.length = FALSE,
         label.offset = 2,
         tip.size = tip_size,
         type.size = type_size,
         ace = ace_baye_part_trees$ARD,
         trees_ace = baye_part_trees,
         main = paste0("Endochondral bone estimation\n(Bayesian partitioned - ARD)"),
         state.names = c("Absent", "Present")
         )
## Plotting a nicer tree
plot.ACE(tree_plot = ladderize(baye_part_conse[[1]]),
         use.edge.length = FALSE,
         label.offset = 2,
         tip.size = tip_size,
         type.size = type_size,
         ace = ace_baye_part_trees$ER,
         trees_ace = baye_part_trees,
         main = paste0("Endochondral bone estimation\n(Bayesian partitioned - ER)"),
         state.names = c("Absent", "Present")
         )
dev.off()
```


```{r, fig.height = 20, fig.width = 14, echo = FALSE}
pdf("Bayesian_unparititioned.pdf", height = 20, width = 14)
## Make this true for printing
par(mfrow = c(1,2))
## Plotting a nicer tree
plot.ACE(tree_plot = ladderize(baye_unpa_conse[[1]]),
         use.edge.length = FALSE,
         label.offset = 2,
         tip.size = tip_size,
         type.size = type_size,
         ace = ace_baye_unpa_trees$ARD,
         trees_ace = baye_unpa_trees,
         main = paste0("Endochondral bone estimation\n(Bayesian unpartitioned - ARD)"),
         state.names = c("Absent", "Present")
         )
## Plotting a nicer tree
plot.ACE(tree_plot = ladderize(baye_unpa_conse[[1]]),
         use.edge.length = FALSE,
         label.offset = 2,
         tip.size = tip_size,
         type.size = type_size,
         ace = ace_baye_unpa_trees$ER,
         trees_ace = baye_unpa_trees,
         main = paste0("Endochondral bone estimation\n(Bayesian unpartitioned - ER)"),
         state.names = c("Absent", "Present")
         )
dev.off()
```
````
